# Comparative gene pathway analysis during adolescent binge-EtOH exposure, withdrawal, and following abstinence

**DOI:** 10.1101/2020.11.02.365841

**Authors:** Alejandro Q. Nato, Hafiz Ata Ul Mustafa, Hannah G. Sexton, Scott D. Moore, James Denvir, Donald A. Primerano, Mary-Louise Risher

**Affiliations:** Department of Biomedical Sciences, Joan C. Edwards School of Medicine, Marshall University, Huntington, WV, USA; Hershel ‘Woody’ Williams Veterans Affairs Medical Center, Huntington WV, USA; Department of Psychiatry and Behavioral Sciences, Duke University Medical Center, Durham, NC, USA; Durham Veterans Affairs Medical Center, Durham, NC, USA

**Keywords:** identification of disease genes, RNA-Seq, GSEA, neuroinflammation, remodeling

## Abstract

**Introduction:** Binge drinking is common among adolescents and young adults and is associated with an increased risk of developing alcohol use disorder (AUD) and long-term cognitive deficits. We analyzed RNA-seq data from male Sprague Dawley rats to identify candidate genes that may play a role in the acute and chronic changes in cognitive function during binge-like adolescent alcohol/EtOH exposure and after a period of abstinence.

**Methods:** At postnatal day (PND) 30, male rats received chronic intermittent EtOH across 16 days. RNA was extracted from hippocampal tissue and sequenced at two acute timepoints, PND 35 and PND 46, and after 24 days forced abstinence (PND 70). We processed RNA-seq data, compiled gene counts, and performed normalization and differential expression analysis (DESeq2). Gene set enrichment analysis was performed through the R package fgsea. Gene sets of the Molecular Signatures Database (MSigDB) collections were used to identify gene pathways that were dysregulated following EtOH exposure. We also evaluated overlapping gene pathways that were affected across all timepoints.

**Results:** Multiple gene pathway analyses revealed that EtOH has robust effects on neuroinflammation, cellular remodeling, sleep, and bioenergetics. Changes were heavily dependent on whether gene expression was assessed during acute EtOH exposure or after abstinence. Genes involved in sleep regulation were selectively impacted during the acute timepoints, whereas dysregulation of genes involved in bioenergetics were only impacted after abstinence. The most striking changes occurred in genes that regulate neuroinflammatory processes and cellular remodeling.

**Conclusion:** These data reveal acute and chronic effects of EtOH on multiple gene pathways that persist across analytic approaches and identify genes that have increased sensitivity to EtOH. These findings contribute to our understanding of the temporal effects of adolescent EtOH exposure and how gene pathway dysregulation contributes to the protracted emergence of neuronal remodeling in the hippocampus during a critical period of brain maturation.

## INTRODUCTION

Alcohol misuse is the third leading preventable cause of death in the USA (CDC, 2022). In 2010, alcohol misuse cost the United States economy ∼$249 billion (Sacks et al., 2015), primarily due to loss of workplace productivity, hospitalizations for acute and chronic alcohol-related injury and disease, and costs relating to the criminal justice system. Binge drinking (4-5 drinks in a 2-hour period in males and females, respectively) comprises ∼75% of the cost of alcohol misuse (Sacks et al., 2015). Alcohol consumption typically begins in adolescence with 90% of alcohol being consumed in a binge drinking manner (DOJ, 2005). The high prevalence of binge drinking coincides with late-stage adolescent brain maturation and a period in which select brain regions are highly vulnerable to the cytotoxic effects of EtOH and other drugs of abuse (Harper and Matsumoto, 2005).

Alcohol use disorder (AUD) is a chronic and complex psychiatric disease, manifested as having uncontrollable drinking patterns. Epidemiological studies reveal that early onset (13-18 years of age) binge drinking is associated with increased likelihood of developing AUD (DeWit et al., 2000, Grant and Dawson, 1997, Hanson et al., 2011, Chin et al., 2010). Early onset binge drinking is also correlated with the emergence of cognitive deficits, including verbal and non-verbal skills, attention, visuospatial function, and learning (Hanson et al., 2011, Chin et al., 2010, Tapert and Brown, 1999, Tapert et al., 2002). Over the last decade, a concerted effort has been made to understand how adolescent onset binge drinking contributes to the development of long-lasting changes in neuronal structure, function and subsequent cognitive changes that may be associated with the emergence of alcohol use disorder.

Parallels are beginning to emerge between human findings and rat models of binge drinking that are consistent across laboratories (Spear and Swartzwelder, 2014, Crews et al., 2019). Hippocampal-dependent tasks, which change the goal location, include the administration of acute ethanol (EtOH) challenge, or have a higher cognitive load, reveal significant deficits in learning processes that persist into adulthood (Risher et al., 2013, Acheson et al., 2013, Coleman et al., 2014, Vetreno and Crews, 2015, Chandler et al., 2022). These behavioral deficits coincide with long-term changes in hippocampal neuronal structure and function (Fujii et al., 2008, Risher et al., 2015a, Sabeti and Gruol, 2008, Swartzwelder et al., 2016), indicative of neuronal remodeling. However, the underlying mechanisms that drive these protracted changes at the synaptic and circuit level remain somewhat elusive. Here we use RNA-Seq to understand the temporal changes that occur in response to repeated adolescent binge EtOH exposure with the goal of elucidating acute and long-term changes in gene regulation that may contribute to synaptic and neuronal remodeling and the development of cognitive dysfunction.

## METHODS

### Adolescent Intermittent Alcohol Exposure

Sprague Dawley rats (*Rattus norvegicus*) SPF grade were obtained from Charles River (NC, USA). At postnatal day (PND) 30 (adolescence), 18 rats received chronic intermittent EtOH (5g/kg intragastrically (i.g.)) or water (H_2_O) 10 times across 16 days (Risher et al., 2015b, Walker et al., 2022). Rats were anesthetized using isoflurane and then euthanized by decapitation. The brains were removed and the hippocampus was rapidly dissected, frozen, and stored at stored at ^-^80°C. Hippocampal tissue was collected from 6 rats each at three time points (Figure 1 and Table S1: Sample ID): 1) 24 hours after 4th dose (PND 35), 2) 24 hours after the tenth dose (PND 46), and 3) 24 days forced abstinence period (PND 70; adult). These time points correlate with adolescence (PND 35 and PND 46) and young adulthood (PND 70) in humans (Sengupta, 2013). All experiments were conducted in accordance with guidelines for care and use of animals provided by the National Institutes of Health and were approved by the Durham Veterans Affairs Medical Center and Duke University IACUC, Durham, NC, USA.

**Figure 1.**
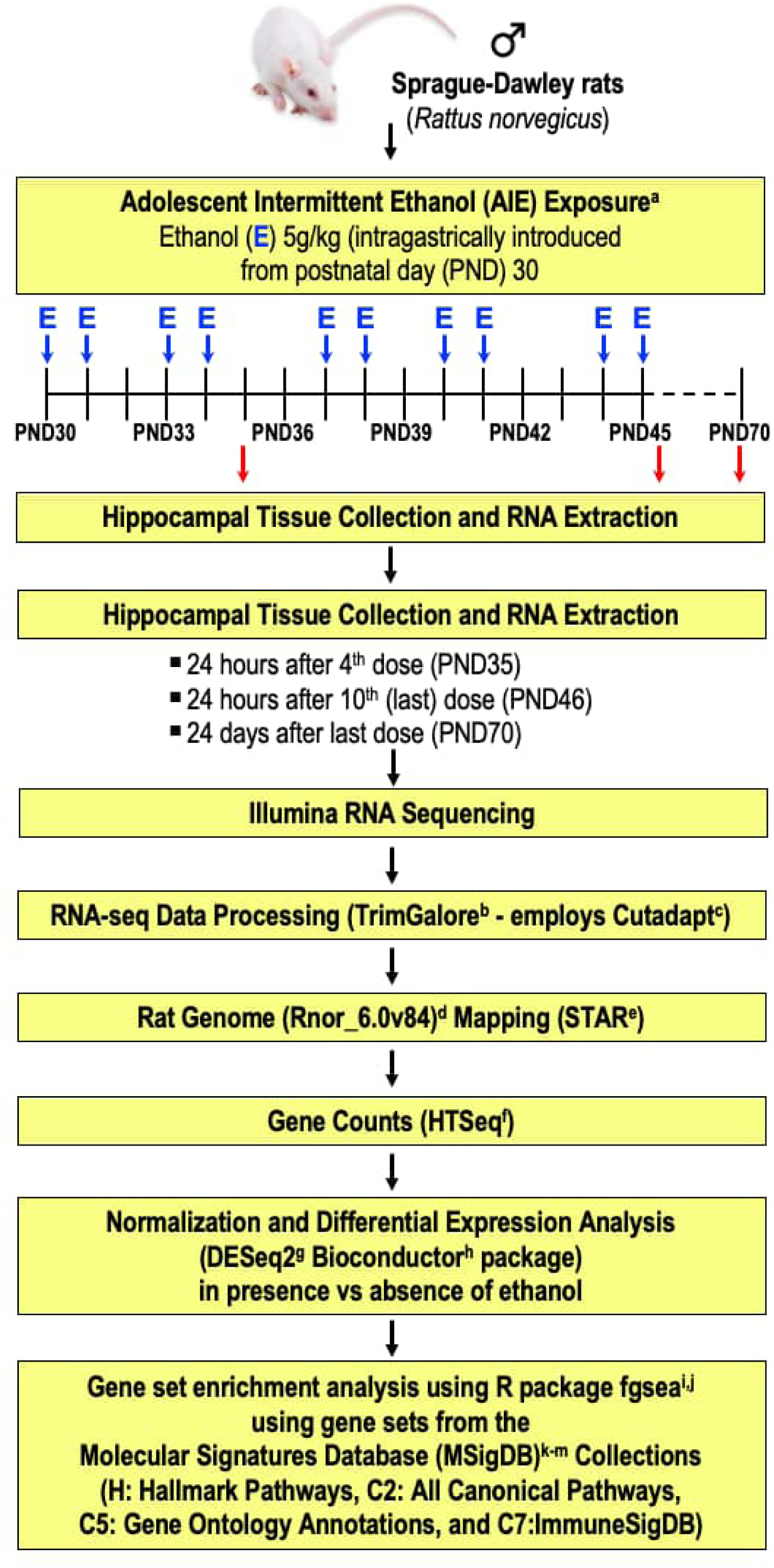
Schematic diagram of methods. Sprague Dawley rats (*Rattus norvegicus*) received chronic intermittent EtOH (5g/kg intragastrically (i.g.)) or water (H2O) 10 times across 16 days (Risher et al., 2015b)^a^. Hippocampal tissue was collected at three time points (PND35, PND46, and PND70) followed by RNA Extraction and RNA Sequencing. RNA-seq data was processed using TrimGalore^b^ that employs Cutadapt^c^. Reads were mapped to the rat genome (Rnor 6.0v84)^d^ through STAR^e^ and gene counts were compiled using HTSeq^f^. Normalization and differential expression analysis were performed using the DESeq2 Bioconductor package^g-i^. Gene set enrichment analysis was performed using R package fgsea^j,k^ using gene sets from the Molecular Signatures Database (MSigDB)^k-m^ Collections. a. Risher et al., 2015b. b. Krueger, 2020. c. Martin, 2011. d. Kersey et al., 2012. e. Dobin et al., 2013. f. Anders et al., 2015. g. Huber et al., 2015. h. Love et al., 2014. i. R Core Team, 2021. j. Korotkevich et al., 2021. k. Subramanian et al., 2005b. l. Liberzon et al., 2011. m. Liberzon et al., 2015.

### RNA Extraction and Sequencing

We extracted total RNA from rat hippocampal tissue using RNeasy mini kit (Qiagen, Germantown, MD) following the manufacturer’s instructions. RNA sample quantity and quality were determined using a NanoDrop 2000 (Thermo Fisher Scientific, Waltham, MA). We used the Roche Kapa mRNA Hyper prep kit (Indianapolis, IN) to prepare libraries following the manufacturer’s instructions. Purified libraries were quantified using Qubit 2.0 Fluorometer (Life Technologies, Carlsbad, CA) and Agilent 2000 Bioanalyzer (Agilent Technologies, Santa Clara, CA). Clusters were generated by cBot with the purified library and sequenced on an Illumina HiSeq 2000 platform (Illumina, Inc., San Diego, CA). All sequencing was performed at the Sequencing and Genomic Technologies Service Center, Duke University, Durham, NC.

### Differential Expression Analysis

We processed RNA-seq data using TrimGalore toolkit (Krueger, 2020), a Perl wrapper that employs Cutadapt (Martin, 2011) and FASTQC (Andrews, 2010) to reliably trim low quality bases and Illumina sequencing adapters from the FASTQ files. Reads that were at least 20 nucleotides in length were aligned and mapped to the Rnor 6.0v84 version of the rat genome and transcriptome (Kersey et al., 2012) using STAR (Dobin et al., 2013). Only reads that mapped to a single genomic location were included in subsequent analyses. Gene counts were compiled using HTSeq (Anders et al., 2015) and filtered to retain genes that had at least 10 reads in any given library. Normalization and differential expression analysis for comparisons at the 3 time points in the presence of EtOH vs. H_2_O, i.e., PND 35 EtOH vs H_2_O, PND 46 EtOH vs H_2_O, and PND 70 EtOH vs H_2_O, respectively, were performed using the DESeq2 Bioconductor package in R (Huber et al., 2015, Love et al., 2014, R Core Team, 2021). To control for multiple hypothesis testing, the false discovery rate (FDR) was calculated using the Benjamini-Hochberg method (Benjamini and Hochberg, 1995). Using all genes in each of conditions above (i.e., presence or absence of EtOH), genes were hierarchically clustered (unsupervised) by correlation distance with complete linkage. Heatmaps were generated using the differentially expressed genes (DEGs) with FDR≤0.05.

### Enrichment Analyses to Identify Candidate Pathways and Genes

Gene set enrichment analysis (GSEA) (Mootha et al., 2003, Subramanian et al., 2005a) computationally assesses whether a defined set of genes statistically distinguishes the difference between two biological states. For each of the comparisons (PND 35 EtOH vs H_2_O, PND 46 EtOH vs H_2_O, and PND 70 EtOH vs H_2_O), we performed GSEA through the R package **`fgsea`** using: (1) all genes (using Ensembl ID) from the differential expression data pre-ranked by −log_10_(p-value)*sign(log_2_(fold change)) (supplementary table 1: PND35, PND46, PND70) and (2) defined annotated human gene sets from the Molecular Signatures Database (MSigDB) Collections, namely: (a) H: Hallmark Pathways, (b) C2: Curated (All Canonical Pathways (CP) subset), (c) C5: Gene Ontology (GO) Annotations, and (d) C7: Immunologic Signatures (ImmuneSigDB subset) (see Table 1 for pathway definitions). We included the top 25 pathways for rat annotated gene sets of Canonical Pathways, GO Annotations, and ImmuneSigDB. For gene sets from Hallmark Pathways, which only has 50 gene sets, we listed the top 10 gene sets. This approach allowed us to determine differentially regulated hallmark, canonical, or immune-related pathways as well as GO terms (biological process, molecular function, and cellular component) leading us to identify candidate pathways and genes associated with binge-drinking and related cognitive dysfunction. We also identified overlapping pathways across timepoints and calculated a global p-value based on their combined adjusted p-values using Fisher’s method. Finally, to understand how these data translate to human health we entered differentially regulated rat genes into the European Bioinformatics Institute (EBI) Experimental Factor Ontology (EFO) database in order to identify human genes whose variants are associated with specific traits from the Genome-Wide Association Studies Catalog (GWAS Catalog) (Buniello et al., 2019). A schematic diagram of the Methods is shown in Figure 1.

**Table 1.** The Molecular Signatures Database (MSigDB) was originally developed for use with GSEA (Liberzon et al., 2015, Liberzon et al., 2011, Köhler et al., 2020, Godec et al., 2016). The latest version of MSigDB consists of seven collections C1-C7 grouped by their location in the human genome. Here we define the various pathways used in the current analysis. For further information see: www.GSEA-msigdb.org.

| Pathway | Definition |
| --- | --- |
| Hallmark (H) | The H collection summarizes most of the relevant information of the original founder gene sets by emphasizing genes that display coordinate expression and represent well-defined biological processes. Hallmark reduces variation and redundancy to provide more refined and concise inputs for gene set enrichment analysis (Liberzon et al., 2011) |
| Canonical (C2) | The C2 collection is divided into the following two sub-collections: Chemical and genetic perturbations and Canonical pathways. The Canonical gene sets are curated from the following online databases: BioCarta, KEGG, Matrisome Project, Pathway Interaction Database, Reactome, SigmaAldrich, Signaling Gateway, SuperArray SABiosciences, WikiPathways |
| Gene Ontology (C5) | The C5 collection is derived from ontology resources. Gene sets in this collection are prefixed with "GOBP" (Biological Process), "GOMF" (Molecular Function), or "GOCC" (Cellular Component) to indicate their source ontology (Köhler et al., 2021). |
| ImmuneSigDB (C7) | The C7 collection represents cell types, states, and perturbations within the immune system. The signatures were generated by manual curation of published studies in human and mouse immunology as part of a collaboration with the Haining Lab at Dana-Farber Cancer Institute and the Human Immunology Project Consortium (HIPC) (Godec et al., 2016). |

## RESULTS

From the differential expression analysis, out of 15,772 genes, there are 132 (0.84%), 143 (0.91%), and 37 (0.23%) genes with |log2FC|≥1 (p<0.05), at timepoints PND35, PND46, and PND70, respectively. After multiple testing correction, only 11 (0.070%), 1 (0.006%), and 1 (0.006%) genes have FDR<0.05 at PND35, PND46, and PND70, respectively (Table S1: PND35, PND46, PND70).

### GSEA using Gene Sets of MSigDB ImmuneSigDB for PND35

Most of the top 25 significantly impacted pathways at PND 35 involve innate cellular immunity genes and included T- and B-cell-related genes (e.g., GSE15930 NAIVE VS 48H IN VITRO STIM CD8 TCELL DN and GSE22886 IGM MEMORY BCELL VS BLOOD PLASMA CELL DN, respectively), monocytes (e.g., GSE7509 UNSTIM VS FCGRIIB STIM MONOCYTE UP), viral, bacterial, and vaccine responsivity (e.g., GSE18791 CTRL VS NEWCASTLE VIRUS DC 18H UP, GSE2935 UV INACTIVATED VS LIVE SENDAI VIRUS INF MACROPHAGE UP, GSE36826 WT VS IL1R KO SKIN STAPH AUREUS INF UP, respectively) (Table 2). Several downregulated pathways [GSE37605 FOXP3 FUSION GFP VS IRES GFP TREG C57BL6 UP, GSE26727 WT VS KLF2 KO LPS STIM MACROPHAGE UP, GSE20715 0H VS 48H OZONE TLR4 KO LUNG UP] include important genes such as, *Mt1*, which encodes the neuroprotective protein metallothionein (log_2_FC: −2.9832), *Ccr9*, that encodes an immune-regulatory marker (CCR9) found on microglia (de Haas et al., 2008) (log_2_FC: −4.0328), and *Gch1* which encodes GTP cyclohydrolase 1 and is involved in monoamine neurotransmitter synthesis and microglial activation (log_2_FC: −3.6795) (Kapatos, 2013, Thöny et al., 2000). While these gene changes are not globally significant (*padj* > 0.1), these data suggest complex changes in microglial immune responsivity and protection against oxidative stress during acute EtOH exposure.

**Table 2.**
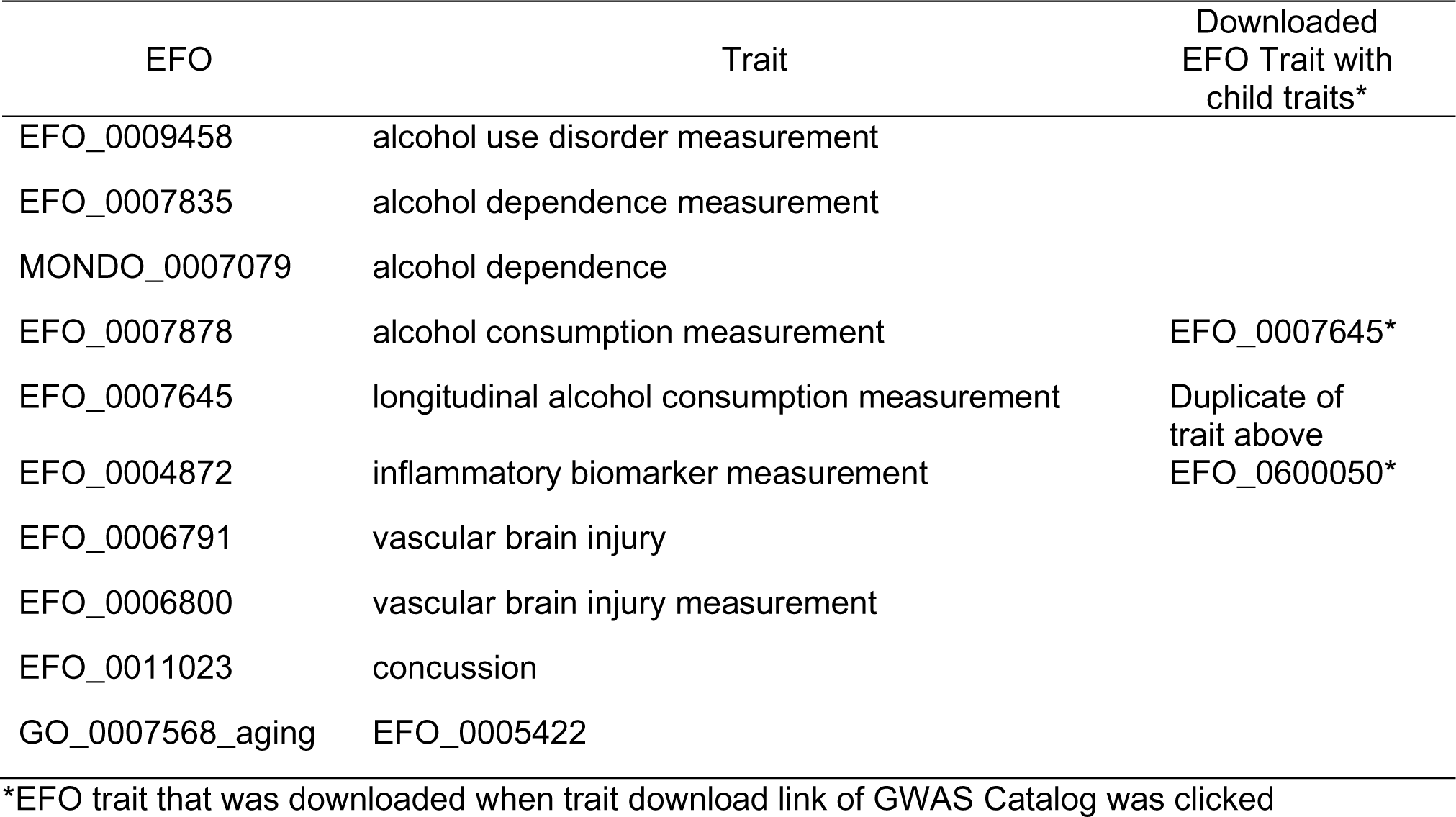
Queried EFO Traits.

| EFO | Trait | Downloaded EFO Trait with child traits* |
| --- | --- | --- |
| EFO_0009458 | alcohol use disorder measurement |  |
| EFO_0007835 | alcohol dependence measurement |  |
| MONDO_0007079 | alcohol dependence |  |
| EFO_0007878 | alcohol consumption measurement | EFO_0007645* |
| EFO_0007645 | longitudinal alcohol consumption measurement | Duplicate of trait above |
| EFO_0004872 | inflammatory biomarker measurement | EFO_0600050* |
| EFO_0006791 | vascular brain injury |  |
| EFO_0006800 | vascular brain injury measurement |  |
| EFO_0011023 | concussion |  |
| GO_0007568_aging | EFO_0005422 |  |

\*EFO trait that was downloaded when trait download link of GWAS Catalog was clicked

### GSEA using Gene Sets of MSigDB Hallmark Pathways for PND35

Upregulated pathways (e.g. HALLMARK_OXIDATIVE_PHOSPHORYLATION) at PND35 include genes that encode for proteins involved in oxidative phosphorylation (Table S2). Downregulated pathways included genes that are important in the regulation of neuroimmune TNFα, NFKβ, TGFβ, and β-catenin signaling (e.g. HALLMARK_HYPOXIA, HALLMARK_TFA_SIGNLAING_VIA_NFKB and HALLMARK_UV_RESPONSE_DN). Consistent with the ImmuneSigDB analysis, *Mt1* and *Gch1* were both downregulated. *Ccn1* gene expression was suppressed (log_2_FC: −1.6937) across 4 downregulated pathways (HALLMARK_HYPOXIA, HALLMARK_TNFA_SIGNALING_NFKB, HALLMARK_UV_RESPONSE_DN, and HALLMARK_EPITHELIAL_MESENCHYMAL_TRANSITION). This gene is involved in regulation of neuroinflammation and extracellular remodeling following injury (Lau, 2011), suggesting suppression of immune and remodeling signaling pathways during acute EtOH exposure.

### GSEA using Gene Sets of MSigDB Canonical Pathways for PND35

17 of the top 25 changed pathways at PND 35 were downregulated (Table S2). Interestingly, 5 out of the top 25 pathways were associated with sensory perception (e.g. PID_RHODOPSIN_PATHWAY and PID_CONE_PATHWAY) (Table S2). Upon closer analysis, many of the genes were not uniquely associated with sensory perception but rather were genes that overlapped robustly with non-motile cilium-related genes. 6 of the top 25 pathways impacted were associated with extracellular matrix and remodeling and were downregulated (e.g. NABA_CORE_MATRISOME and NABA_ECM_GLYCOPROTEINS). As observed in Hallmark pathway analysis, *Ccn1* was downregulated, as was *Esm1* (log_2_FC: −3.3555), which encodes the endothelial cell-specific molecule (ESM1) proteoglycan expressed and secreted by endothelial cells that is involved in the regulation of angiogenesis, inflammation, and permeability (Rocha et al., 2014). Three pathways involved in sleep/wake cycle regulation were significantly downregulated in response to binge EtOH exposure (for example, WP_CIRCADIAN_RHYTHM_RELATED_GENES) and involved robust downregulation of *Aanat* (log_2_FC: −4.0138), *Tph1* (log_2_FC: −6.8856), and *Drd4* (log_2_FC: −4.6455). These genes are critical in the regulation of night/day rhythm and particularly important in the synthesis of melatonin, serotonin, and the encoding of the dopamine receptor D4 (Coon et al., 1996, Kurhaluk and Tkachenko, 2020) (Hamdan and Ribeiro, 1999).

### GSEA using Gene Sets of MSigDB GO Annotations for PND35

As seen in MSigDB Canonical Pathway analysis, the majority of the top impacted pathways were related to the downregulation of sensory perception and overlap significantly with the non-motile cilium pathways (e.g. GOBP_SENSORY_PERCEPTION_OF_LIGHT_STIMULUS and GOBP_SENSORY_PERCEPTION) (Table S2). There was also a robust downregulation of pathways associated with cellular remodeling and reorganization (for example, GOMF_EXTRACELLULAR_MATRIX_STRUCTURAL_CONSTITUENT). The top affected gene was *Ccn1*, consistent with ImmuneSigDB and Canonical analysis. 3 gene pathways involved in sleep modulation and regulation were significantly downregulated in response to binge EtOH exposure and involved genes that emerged in the previous pathway analysis; *Aanat*, *Tph1*, and *Drd4*.

### GSEA using Gene Sets of MSigDB ImmuneSigDB for PND46

For the PND 46 time point, 8 of the top 25 pathways impacted by EtOH were CD8-related and were downregulated (e.g. GSE9650_NAIVE_VS_EFF_CD8_TCELL_DN and GOLDRATH_NAIVE_VS_EFF_CD8_TCELL_DN) (Table S2), suggesting increased vulnerability to viral infection. A pathway involved in the innate immune response (GSE22140_HEALTHY_VS_ARTHRITIC_GERMFREE_MOUSE_CD4_TCELL_DN) was downregulated and included an important neuro-immune-related cytokine *Il1β* (log_2_FC: −2.1624). Consistent with the earlier timepoint (PND 35), pathways involving *Gch1* were downregulated (e.g. GSE1432_CTRL_VS_IFNG_24H_MICROGLIA_DN), as was *Atf3* that encodes transcription factor 3 (ATF3) (log2FC: −2.0685) (e.g. GSE7509_DC_VS_MONOCYTE_UP). ATF3 is a member of the CREB family and plays numerous roles in immune regulation and is a marker for neuronal injury (Carlton et al., 2009, Kataoka et al., 2007). Overall, these data suggest that immune processes are negatively impacted by repeated adolescent EtOH exposure.

### GSEA using Gene Sets of MSigDB Hallmark Pathways for PND46

Focusing on the top 10 significantly modified pathways, we found that 5 of the 10 top enriched pathways at PND 46 involve genes related to neuroimmune signaling and cell death, all of which were downregulated (Table S3). Consistent with ImmuneSigDB, there was downregulation of *Il1β* and *Gch1*, with the additional downregulation of *Il2ra* (log2FC: −1.5139) (e.g. HALLMARK_APOPTOSIS and HALLMARK_IL6_JAK_STAT3_SIGNALING), emphasizing the presence of EtOH-induced immune suppression.

### GSEA using Gene Sets of MSigDB Canonical Pathways for PND46

When focusing on the top 25 most significantly changed pathways at PND 46, we found that the top 15 of 25 pathways showed a downregulation of gene expression pathways involved in extracellular matrix (ECM) protein regulation and neuronal and glial remodeling (Table S3) (NABA_CORE_MATRISOME and REACTOME_EXTRACELLULAR_MATRIX_REORGANIZATION). Three of these 15 pathways involved specific integrin-related genes sets involved in synaptic remodeling and synaptic transmission. *Il1β* was downregulated and was accompanied by the downregulation of *Gdf7* (log_2_FC: −2.0211) (NABA_MATRISOME_ASSOCIATED) that encodes TGFβ. TGFβ is a critical regulator of neuroinflammatory processes (Veldhoen and Stockinger, 2006), again demonstrating an overall loss of immune-related gene expression that may contribute to increased risk of infection at this timepoint.

### GSEA using Gene Sets of MSigDB GO Annotations for PND46

4 of the top 25 pathways were downregulated at PND 46 and involved ECM genes and genes involved in external cell structure (Table S3) (e.g. GOMF_EXTRACELLULAR_MATRIX_STRUCTURAL_CONSTITUTENT). While not high on the list (24^th^ pathway), the ‘GOBP CELL SUBSTRATE ADHESION’ pathway was also downregulated and involves the encoding of cell attachment proteins and underlying substrates such as ECM proteins. 14 of the top 25 pathways involve the dysregulation of genes important for synaptic organization and dendritic development (e.g. GOBP_SYNAPSE_ORGANIZATION and GOBP_DENDRITE_DEVELOPMENT). Similar to the gene ontology findings at PND35, one pathway associated with sensory perception remains robustly downregulated and driven by genes that encode the function and regulation of non-motile cilium within the hippocampus (GOBP_SENSORY_PERCEPTION_OF_LIGHT_STIMULUS).

### GSEA using Gene Sets of MSigDB ImmuneSigDB for PND70

At the oldest timepoint (PND 70), the top 25 pathways within ImmuneSigDB were all upregulated (Table S4). 9 of top 25 gene pathways impacted by EtOH are involved in T-cell regulation (e.g., GSE42021_TREG_PLN_VS_TREG_PRECUSRORS_THYMUS_DN and GSE42021_CD24HI_VS_CD24INT_TREG_THYMUS_DN). Several genes were particularly responsive to EtOH exposure: *Gbp6* encodes Guanylate Binding Protein Family Member 6 (GBP-6) was significantly upregulated (log_2_FC: 1.91) (e.g. GSE36527_CD69_NEG_VS_POS_TREG_CD62l_LOS_KLRG1_NEG_UP) suggesting increased immune responsivity (Kim et al., 2011); *Ccn1* was enriched (log_2_FC: 2.2094) (GSE4748_CTRL_VS_LPS_STIM_DC_3H_UP) and as mentioned previously is involved in remodeling and inflammatory events (Lau, 2011). These data suggest that immune response rebounds after a period of abstinence, resulting in the upregulation of immune response and ECM remodeling.

### GSEA using Gene Sets of MSigDB Hallmark Pathways for PND70

Focusing on the top 10 impacted pathways at PND 70, analysis demonstrated that two pathways were downregulated while the other 8 were upregulated (Table S4). The two pathways negatively impacted by EtOH (HALLMARK_OXIDATIVE_PHOSPHORYLATION and HALLMARK_MYC_TARGETS_V1) involve genes that encode proteins involved in oxidative phosphorylation and factors involved in the regulation of growth and cell cycle entry. The remaining 8 pathways were all upregulated in response to EtOH. The most effected genes include: *Ccn1*, *Gbp6*, *Atf3* (log_2_FC: 1.5869) (e.g., HALLMARK_TNFA_SIGNALING_VIA_NFKB and HALLMARK_INTERFERON_GAMMA_RESPONSE). These data again suggest that adolescent EtOH exposure results in upregulation of immune function during abstinence.

### GSEA using Gene Sets of MSigDB Canonical Pathways for PND70

8 of the top 25 pathways were upregulated at PND 70 and involve ECM remodeling (e.g. REACTOME_EXTRACELLULAR_MATRIX ORGANIZATION and NABA_CORE_MATRISOME) (Table S4). Four of the top 25 pathways include the downregulation of gene pathways involved in bioenergetics and mitochondrial dynamics (e.g. REACTOME_RESPIRATORY_ELECTRON_TRANSPORT_ATP_SYNTHESIS_BY_CHEMIOS MOTIC_COUPLING_AND_HEAT_PRODUCTION_BY_UNCOUPLING_PROTEINS and WP_MITOCHONDRIAL_COMPLEX_I_ASSEMBLY_MODEL_OXPHOS_SYSTEM), suggesting long-term impairment of cellular energetic processes. One gene that was robustly downregulated across multiple pathways was *Calcr* (log_2_FC: −2.4472) (e.g. REACTOME_G_ALPHA_S_SIGNALLING_EVENTS); this gene encodes the high affinity G-protein coupled receptor (GPCR) calcitonin and is involved in maintaining calcium homeostasis in a variety of tissue and species (Lafont et al., 2011, Copp et al., 1962).

### GSEA using Gene Sets of MSigDB GO Annotations for PND70

At PND 70, 11 of the top 25 pathways were downregulated and are related to bioenergetics (Table S4). The most relevant genes impacted within these pathways include *Rplp1* and *Rpl12* (e.g. GOMF_STRUCTURAL_CONSTITUENT_OF_RIBOSOME). The *Mcub* gene was not significantly downregulated, however was the most impacted gene within the 2.106, adjusted p-value (padj) = 8.759×10^-5^). This may be important since it encodes the mitochondrial calcium uniporter (MCUB) and is involved in cellular stress response (Lambert et al., 2019). 6 of the top 25 gene pathways were related to the encoding and regulation of ECM proteins and were upregulated. As observed in Hallmark pathway analysis, *Ccn1* was upregulated (e.g. GOMF_EXTRACELLULAR_MATRIX_STRUCTURAL_CONSTITUENT), suggesting an increase in remodeling events during abstinence.

### Gene pathway overlap across timepoints

We conducted further analysis to assess gene pathway overlap across all 3 timepoints (using adjusted p-value (padj) <0.05). Using Hallmark pathway analysis, only 2 gene pathways were consistently downregulated by EtOH at both PND 35 and PND 46: ‘EPITHELIAL MESENCHYMAL TRANSITION’ and ‘TNFA SIGNALING VIA NFKB’. However, after the prolonged period of abstinence, in adulthood (PND 70) both of these pathways were upregulated (NES: −1.54, padj = 3.78×10^-10^; NES: −1.6, padj = 1.57×10^-5^, respectively) (Figure 2). Epithelial-mesenchymal transition allows polarized epithelial cell conversion to a mesenchymal cell phenotype and is particularly important in cellular remodeling events (Kalluri and Weinberg, 2009). NFκβ signaling has been shown to be important for the epithelial-mesenchymal transition (Huber et al., 2004), suggesting a loss of regulation over these remodeling events following acute EtOH.

**Figure 2.**
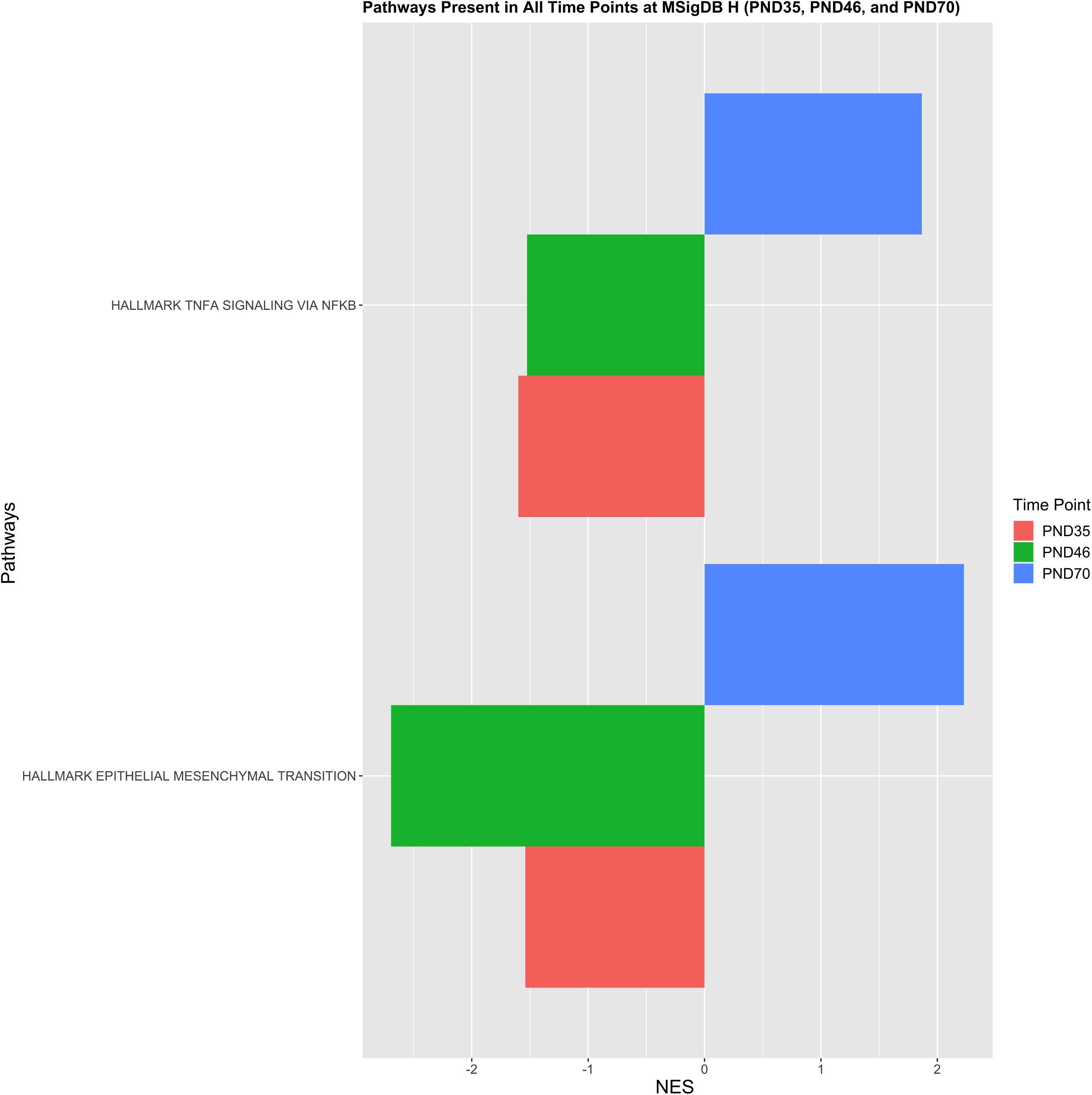
Pathways significantly dysregulated at all 3 time points based on the MSigDB Hallmark Pathways (H) gene sets. 2 pathways were downregulated at both acute time points (PND 35 and PND 46) and upregulated after abstinence (PND 70).

Canonical pathway analysis demonstrated that 4 gene pathways were consistently downregulated at both PND 35 and PND 46 in response to EtOH yet upregulated following the abstinence period (PND70). All 4 of these pathways are involved in cellular remodeling events: REACTOME ECM PROTEOGLYCANS, NES: 1.869, padj = 5.82×10^-5^; NABA CORE MATRISOME, NES: 1.975, padj = 5.82×10^-5^; NABA ECM GLYCOPROTEINS, NES: 1.729, padj = 2.11×10^-2^; REACTOME ELASTICE FIBRE FORMATION, NES: 2.064, padj = 8.24×10^-4^) (Figure 3).

**Figure 3.**
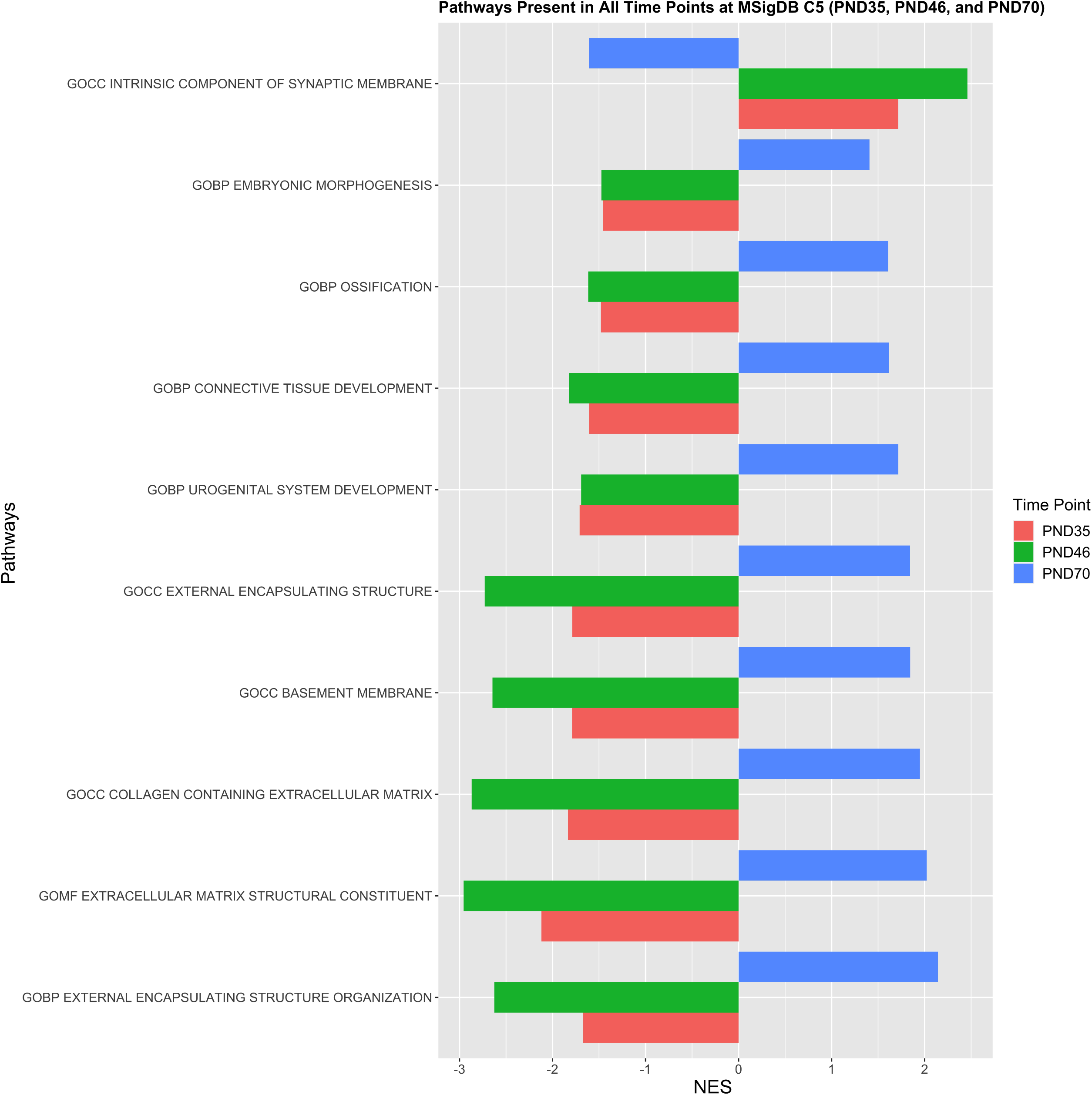
Pathways dysregulated at all 3 time points based on the MSigDB Canonical Pathways (C2) gene sets. 4 pathways were downregulated at both acute time points (PND 35 and PND 46) and upregulated after abstinence (PND 70).

Using the Gene Ontology pathway analysis, 10 pathways were significantly dysregulated by EtOH at all 3 time points (Figure 4). 9 of the 10 pathways showed downregulation of identical pathways at both acute time points (PND 35 and PND 46) and upregulation of these same pathways after abstinence (PND 70). The majority of these gene pathways are related to cellular and membrane organization, synaptic organization, receptor expression and organization, and neurotransmitter activity and synaptic vesicle recycling, suggesting that there is suppression of organizational components of cellular and synaptic remodeling events during EtOH exposure and subsequent upregulation during the abstinence period. The outlier is the ‘INTRINSIC COMPONENT OF SYNAPTIC MEMBRANE’ pathway in which we see EtOH-induced upregulation at both acute time points and downregulation following the abstinence period (NES: −1.609, padj = 4.66×10^-2^).

**Figure 4.**
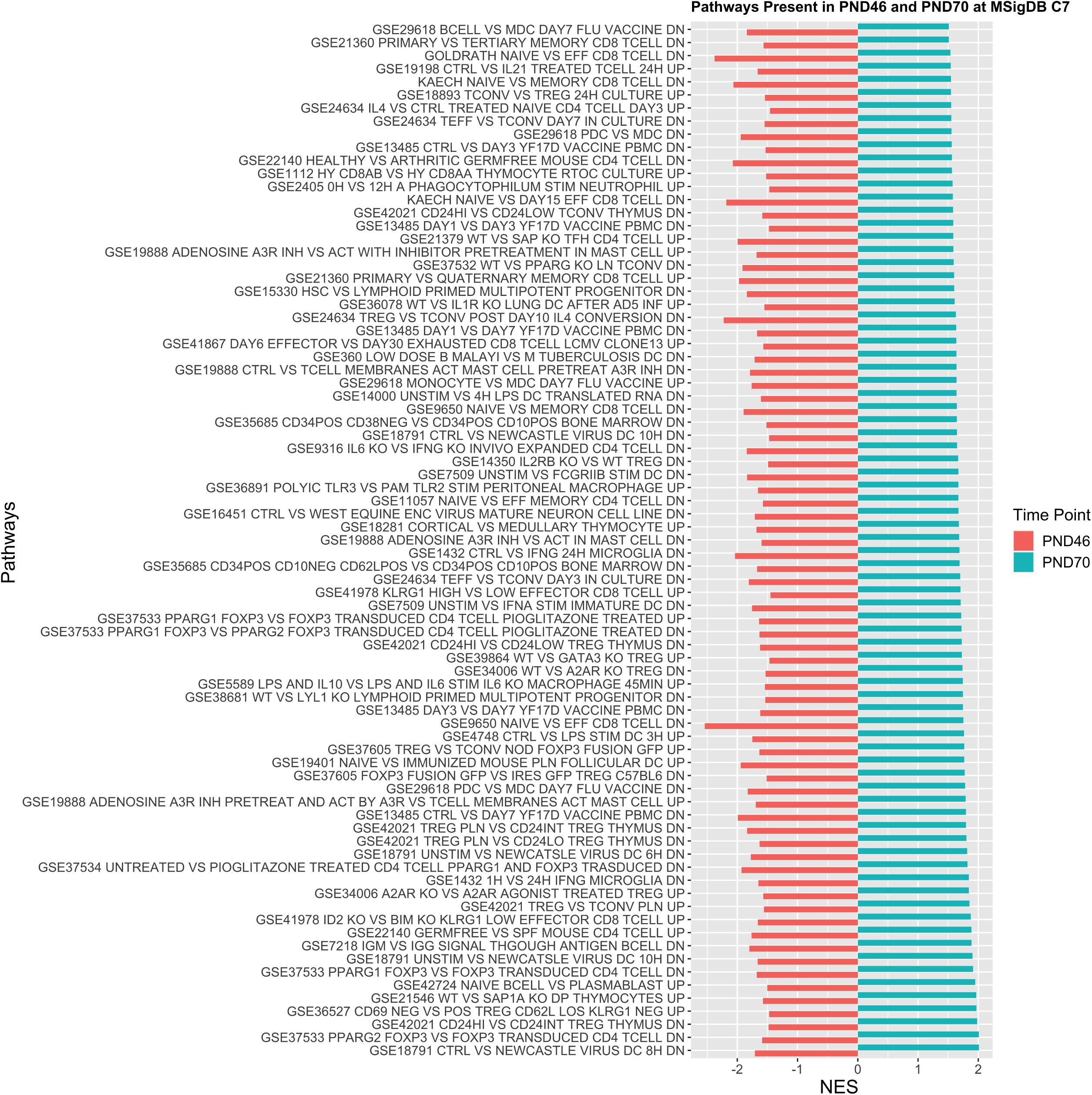
Pathways significantly dysregulated at all 3 time points based on the MSigDB GO Annotation (C5) gene sets. 9 of these 10 pathways were downregulated at both acute time points (PND 35 and PND 46) and upregulated after abstinence (PND 70).

The ImmuneSigDB analysis resulted in less robust patterns of pathway up- and downregulation relative to time point following EtOH exposure. There were no gene pathways consistently impacted by EtOH across all time points and there were no overlapping pathways across the two acute time points (PND 35 and PND 46) (Figure 5). However, 79 gene pathways that were downregulated at PND46 were upregulated following abstinence, suggesting that repeated exposure could be critical for the somewhat delayed suppression of these immune pathways.

**Figure 5.** Pathways dysregulated at PND46 and PND70 based on the MSigDB ImmuneSigDB (C7) gene sets. 79 pathways that were downregulated at PND46 were also upregulated following abstinence.

### Identification of human homologs of the candidate genes

To identify human homologs of candidate genes from the various dysregulated pathways, we parsed the variants associated with several disease traits from the Genome-Wide Association Studies Catalog (GWAS Catalog). Based on the gene sets in which we found the most pronounced changes, we focused on 14 EFO traits (Table 2). From the 14 EFO trait comparisons, we found 36 genes of interest (Table 3) and a total of 5 genes that are similarly impacted in humans that spanned the traits of immune function, alcohol consumption, and sleep. The gene IL1b is an important cytokine involved in the innate immune response and is downregulated across multiple pathways at the second acute time point (PND 46) but recovers to age-matched control levels following a period of abstinence. Based on a GWAS in humans, *IL1B* is associated with inflammation (EFO 0004872: inflammatory_biomarker_measurement) (Offenbacher et al., 2018), suggesting acute immune suppression during EtOH exposure. In support of this finding, *ATF3* is downregulated in our animal model of EtOH exposure across multiple pathways at PND 46 and upregulated after abstinence, in adulthood. The protein that *ATF3* encodes plays multiple roles in immune regulation and has been used as a marker for neuronal injury (Carlton et al., 2009, Kataoka et al., 2007). In humans, it has also been identified as a biomarker for inflammation (EFO 0004872: inflammatory_biomarker_measurement) (Fatumo et al., 2019). Overall, these data suggest that genes associated with these immune processes are depressed by repeated acute EtOH exposure and recover or become upregulated following abstinence.

**Table 3.**
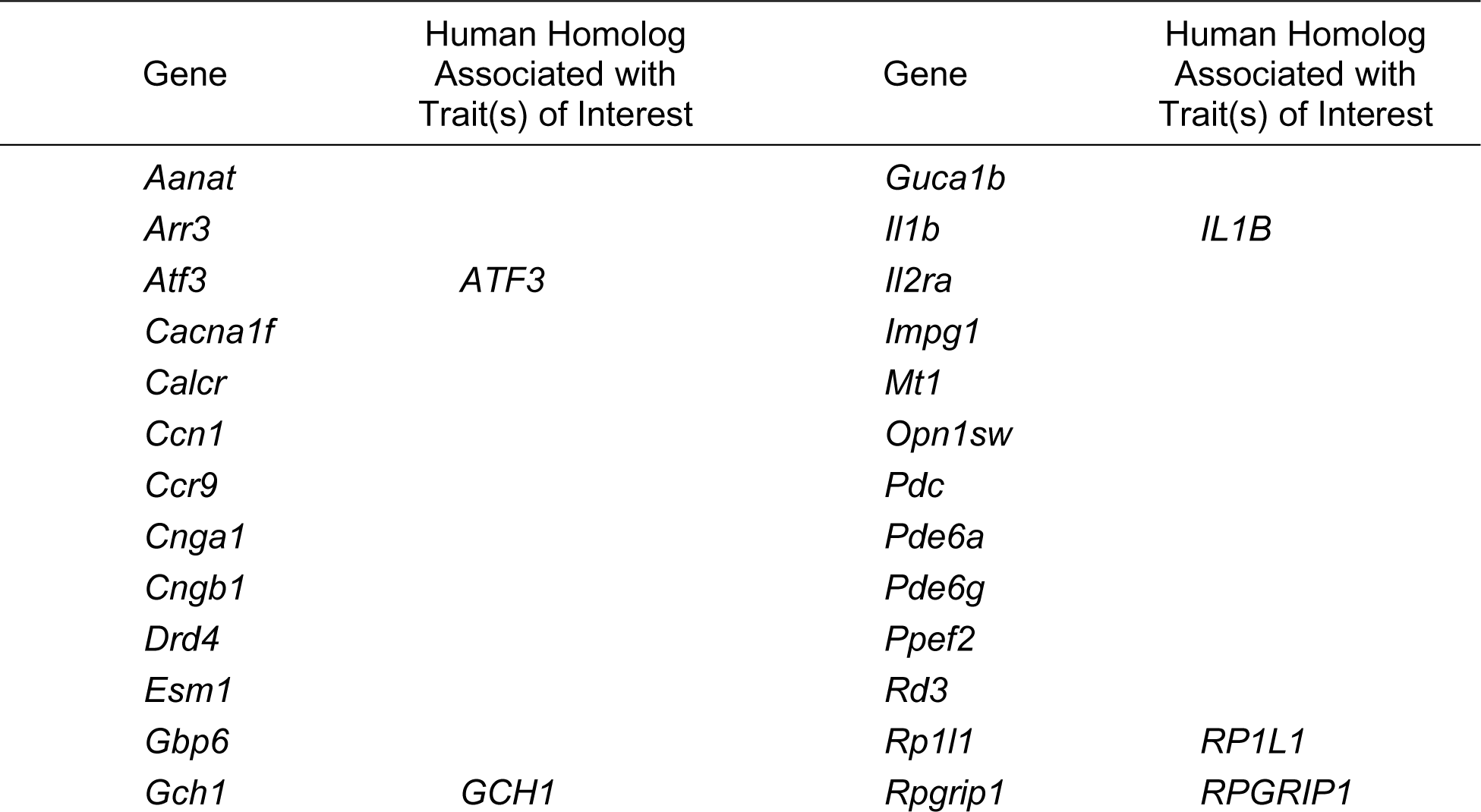
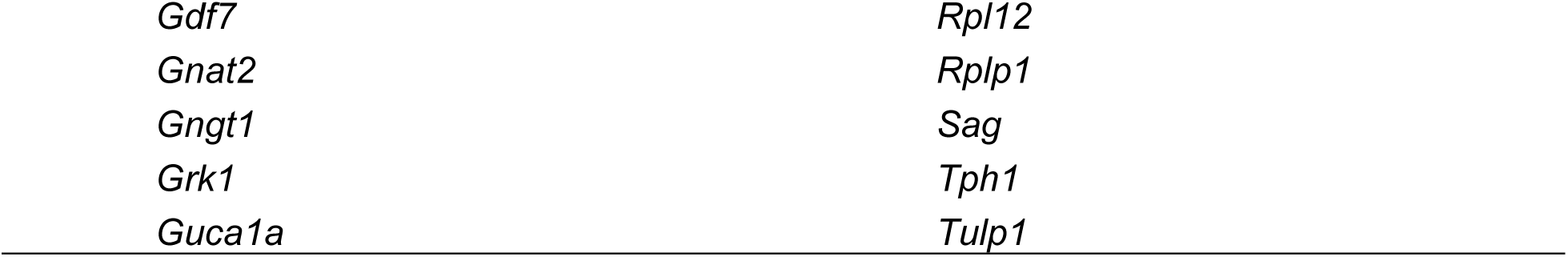
Genes Perturbed During and After Adolescent EtOH Exposure.

The third gene with a human homologue associated with EFO 0004872: inflammatory_biomarker_measurement was *Rpgrip1* (Han et al., 2020). Interestingly, *Rpgrip1* was only downregulated by EtOH at the early acute time point (PND 35) and recovered to age-matched control levels by PND 46. *Rpgrip1* encodes RPGR interacting protein 1 and is not well characterized outside of the eye. It is thought to function as a scaffolding protein, be involved with the function of non-motile cilium, and has been implicated the development of immune-related autosomal recessive congenital blindness (Roepman et al., 2005) due to the increased production of reactive oxygen species and inflammation (Li et al., 2021).

There was a pronounced change in one gene that matched the human trait assessing alcohol consumption (EFO_0007878: alcohol_consumption_measurement) (Feitosa et al., 2018). The gene *Rp1l1* was significantly downregulated but only at the first acute EtOH time point. Similar to *Rpgrip1,* this gene is involved in sensory perception and non-motile cilium-related function. However, a new role has been found for this gene in hypertension (Feitosa et al., 2018) that may be important in understanding the relationship between cardiovascular disease and excessive drinking.

*GCH1* is downregulated acutely during EtOH exposure ((PND35 and PND46) but recovers to age-matched control levels following a period of abstinence. This gene encodes GTP cyclohydrolase 1 and is involved in the upstream processes that generate a molecule called tetrahydrobiopterin (BH4) that is necessary for the synthesis of dopamine and serotonin. Given the role of serotonin in mood, emotion, sleep, and appetite it should be no surprise that the *Gch1* gene in rats has been identified as a gene associated with sleep quality in humans (Spada et al., 2016).

## DISCUSSION

We used RNA-seq analysis to understand the temporal gene changes that occur in response to repeated adolescent EtOH exposure. Using multiple timepoints (two during acute EtOH exposure (PND 35, PND 46) and one after abstinence, during adulthood (PND 70), we have identified vulnerable gene pathways and specific genes that are dysregulated in response to EtOH. We also compared gene changes to the GWAS catalogue to identify human homologs of candidate genes impacted by repeated adolescent EtOH exposure.

## SLEEP

Multiple pathways involved in sleep modulation and regulation were downregulated during acute EtOH exposure. *Aanat*, a downregulated gene identified within several of these pathways, that encodes the penultimate enzyme in the synthesis of melatonin and controls the night/day rhythm in melatonin production (Coon et al., 1996, Kurhaluk and Tkachenko, 2020). It is well understood that dysregulation of melatonin can disrupt the sleep/wake cycle. Moreover, there is convincing evidence that EtOH can cause significant sleep disruption in both human and animal models of adolescent binge drinking (Chakravorty et al., 2016, Ehlers et al., 2018).

Compounding these effects is the downregulation of *Tph1*, the gene that encode tryptophan hydroxylase 1 (TPH1). TPH1 is the enzyme that catalyzes the rate-limiting step in the synthesis of serotonin (5-HT) (Hamdan and Ribeiro, 1999) and since melatonin is derived from 5-HT it is no surprise that melatonin would also be affected by a reduction of *Tph1* encoding (Liu et al., 2019). There are two distinct genes that encode for tryptophan hydroxylase (TPH1 and TPH2) and variations in TPH gene expression have been associated with aggression, schizophrenia, AUD, substance use disorder (SUD), and depression in humans. TPH2 has been a major focus of the alcohol field due to its direct role in major depressive disorder and major affective disorder (Gao et al., 2012). However, there is evidence that TPH1 and TPH1 polymorphisms are associated with psychiatric conditions such as personality disorder (Wilson et al., 2012), major depressive disorder (Wang et al., 2011), and suicidal behavior (Liu et al., 2006). Recent work by Wang et al (2016) demonstrated a higher incidence of polymorphisms in the TPH1 gene in opiate-dependent patients but not in alcohol-dependent patients. This raises the question of whether adolescent binge EtOH exposure can selectively contribute to a higher propensity for substance use and dependence later in life through the alteration of TPH1 gene expression or whether TPH1 alterations can predict the level of consumption and escalation, or even contribute to later vulnerability to SUD.

The *Drd4* gene that encodes the dopamine receptor D4 (D4R) was acutely downregulated (at PND 35). Recent work has demonstrated that D4R signaling is critically involved in regulating sleep-wake sleep duration and that in the presence of a D4R antagonist, the time spent asleep was significantly reduced when compared to awake time (Nakazawa et al., 2015, Cavas and Navarro, 2006). Dysregulation and mutation of *Drd4* have been associated with multiple behavioral phenotypes linked to the increased propensity to consume illicit drugs, these behaviors include attention deficit disorder, novelty seeking, and disinhibition (Grady et al., 2003, Rogers et al., 2004, George et al., 1993, Wand et al., 2001, Wingo et al., 2016). A meta-analysis, which assessed Drd4 VNTR variation in patient data, revealed that polymorphisms within *Drd4* may be a risk factor for problematic alcohol use (Daurio et al., 2020). In genetic mouse models, studies have shown that a reduction in *Drd4* increases disinhibition, and impairs cognitive flexibility (Young et al., 2011), while increasing exploratory behavior (Thanos et al., 2015). Dysregulation in these types of behaviors have been implicated in the increased propensity to consume drugs and alcohol in both humans and animal models (George et al., 1993, Wand et al., 2001, Wingo et al., 2016, Thanos et al., 2015).

Even though the presence of these gene changes does not persist into adulthood, it is important to remember that adolescence and the emergence into young adulthood is a period in which many psychiatric disorders emerge (see Giedd for review (2008)). Moreover, there is a growing appreciation for the bi-directional relationship between psychiatric disorders, AUD, and sleep disorders. These data raise the question of whether repeated EtOH-induced sleep disruption during this developmental period contributes to an increased risk of the emergence of psychiatric disease and whether these EtOH-induced changes interfere with normal adolescent adaptations required for a successful transition to adulthood.

## REMODELING

At both acute timepoints, there was downregulation of gene pathways related to cellular remodeling and reorganization, which interestingly involve some genes that have overlapping roles in neuroimmune signaling and cellular remodeling. *Ccn1* is one such gene that was downregulated at both early timepoints and is involved in neuroimmune signaling (discussed previously, Chen et al., (2007)) and cellular remodeling (Jun and Lau, 2010, Jones and Bouvier, 2014b). *Ccn1* encodes a secreted ECM protein that interacts with multiple integrins and heparin sulfate proteoglycans and plays many important roles in inflammatory response, apoptosis, cell migration, cell proliferation, and extracellular remodeling following injury (Lau, 2011). *Esm1* is another example of a gene that was downregulated at the earliest timepoint and is involved in neuroimmune signaling and cytoskeletal reorganization. This proteoglycan is expressed and secreted by endothelial cells and interacts with numerous immune signals (Béchard et al., 2001). The EtOH-induced downregulation of four cellular remodeling pathways at PND 35 were more robustly downregulated at PND 46, likely due to the continued EtOH exposure (i.e., 4 binge doses versus 10 binge doses of EtOH). This was accompanied by the downregulation of multiple integrin-related gene sets. These findings have important implications due to the highly dynamic nature of ECMs that contribute greatly to neuronal and glial remodeling though protein degradation and deposition (Lu et al., 2011). Combined with the enrichment of multiple gene pathways involved in synaptic organization and dendritic development, these data suggest that degradation of ECM could open a window of opportunity for later deposition-related synaptic remodeling. Supporting this hypothesis was the finding that by adulthood (PND70), many pathways involved in the regulation of ECM-related remodeling were upregulated suggesting delayed deposition of ECM proteins and protracted remodeling that only occur within the abstinence period.

Many gene pathways related to non-motile cilium were impacted by EtOH. The complete functional role of non-motile cilium-related genes is yet to be determined; however, these genes have been positively identified in neurons and astrocytes (Berbari et al., 2007, Fuchs and Schwark, 2004). They have been shown to play a critical role in regulating hippocampal neurogenesis as demonstrated by Breunig et al., (2008). Using a mouse with genetic deletion of primary cilium, they demonstrated that loss of primary cilium resulted in abnormalities in glial and neuronal morphometrics, disruption of proliferative neurogenesis, and failed (ectopic) integration of newborn neurons into the granule cell layer of the dentate gyrus. Interestingly, dysregulation of neurogenesis within the granule cell layer (Morris et al., 2010, Vetreno et al., 2018, Reitz et al., 2021) and failed (ectopic) integration of newborn neurons (McClain et al., 2014) has also been reported in models of adolescent EtOH exposure, suggesting that dysregulation of cilium may play a critical role in the EtOH-induced disturbances in neurogenesis that occur within the dentate gyrus. However, the role of non-motile cilium is yet to be thoroughly investigated. Overall, these data suggest that adolescent EtOH acutely suppresses pathways important for non-motile cilia and ECM-related remodeling events followed by what appears to be a rebound/compensatory upregulation during the abstinence period. These events may be critically important since the brain is undergoing the final stages of maturation, making it increasingly vulnerable to late-stage and permanent EtOH-induced remodeling. Further work is necessary to determine the impact of this remodeling on hippocampal function and whether other brain regions undergo similar temporally-dependent events.

## NEUROINFLAMMATION

At the earliest timepoint (PND35), there was downregulation of multiple pathways involved in neuroprotection against oxidative stress and the innate immune system. The *Mt1* gene, which encodes the protein metallothionein, was downregulated across multiple MSigDB collections. This protein is a cellular redox sensor and plays an important role in protecting cells against hydroxyl free radicals that are generated during oxidative stress (Eibl et al., 2010). Two downregulated pathways involve cytokine and microglial immune responsivity. One of the genes most impacted in these pathways was *Ccr9*, which encodes an immunoregulatory marker (CCR9) found on microglia (de Haas et al., 2008) and is involved in feed forward neuroinflammatory action and neurodegeneration (Li et al., 2006), suggesting diminished neuroimmune responsivity during acute EtOH. Another microglial-related gene that is downregulated is *Gch1*, encodes GTP cyclohydrolase-1. While typically associated with the encoding of a rate limiting enzyme in the synthesis of tetrahydrobiopterin (BH4) and the synthesis of monoamine neurotransmitters (Kapatos, 2013, Thöny et al., 2000), recent work demonstrates that deletion of *Gch1* in zebrafish impairs microglial activation (Larbalestier et al., 2022). Further support of diminished immune regulation at this early timepoint emerges with the downregulation of the gene that encodes activating transcription factor 3 (ATF3). ATF3 is a member of the CREB family and plays numerous roles in immune regulation. Recent work demonstrates that ATF3 dysregulation occurs following acute EtOH (4.5g/kg, i.g.) in young adult mice and contributes to suppression of immune regulation through dysregulation of TNFα (Hu et al., 2017).

Downregulation of immune-related pathways did persist throughout the second acute EtOH timepoint (PND46), demonstrating continued loss of immune regulation. This was compounded by the downregulation of multiple pathways involved in CD8 function. This is consistent with previous studies that have reported lowered CD8 T cell counts in mice after repeated EtOH exposure and decreased efficiency in clearing viral infection (Loftis et al., 2016), and further corroborated in studies showing immune suppression in patients who have consumed EtOH prior to sustaining an injury (Messingham et al., 2002). Genes encoding for IL2rα, IL1β, and TGFβ were also downregulated. Given the role of these cytokines as mediators in innate immune response, loss of these factors has been associated with chronic infection and generalized immune dysfunction (Lehman and Ballow, 2008, Dinarello, 2018, Kelly et al., 2017). Moreover, dysregulation of IL1β has been implicated in several CNS inflammatory disease processes such as multiple sclerosis and experimental autoimmune encephalomyelitis (Paré et al., 2017) and neuroimmune driven neurodegenerative processes, when released from microglia following insult (Zhang et al., 2018). In the context of repeated EtOH exposure, immune responsivity with regard to microglial activation and TGFβ become more complex (see (Peng and Nixon, 2021) for further reading). In combination, these results demonstrate that oxidative stress pathways and immune processes are negatively impacted during acute EtOH exposure, leaving the brain more vulnerable to infection. Interestingly, after a period of abstinence, many of these pathways become upregulated, resulting in a switch from immune suppression to increased immune responsivity. This rebound upregulation was also seen in multiple pathways and particularly impacted genes that are involved in immune responsivity (Kim et al., 2011) and ECM remodeling (Jun and Lau, 2010, Jones and Bouvier, 2014a), once again demonstrating upregulation of neuroimmune function during abstinence that likely contributes to ongoing cellular remodeling.

## BIOENERGETICS

One interesting occurrence was the downregulation of multiple gene pathways involved in bioenergetics and mitochondrial dynamics that only occurred after the forced abstinence period, in adulthood. This is consistent with previous work by Tapia-Rojas et al (2018) in which they demonstrate persistent disruption in the expression of multiple protein components involved in the regulation of hippocampal mitochondrial function after adolescent binge EtOH exposure. The *Mcub* gene was the most impacted gene within one of these significantly downregulated pathways. This gene encodes the mitochondrial calcium uniporter dominant-negative subunit beta (MCUB), which is responsible for negatively regulating calcium uptake into mitochondria thereby limiting mitochondrial calcium overload during stress (Lambert et al., 2019). Loss of MCUB results in dysregulation of anti-inflammatory processes in injury models (Feno et al., 2021), while induction of MCUB has been shown to protect the heart from post-ischemic remodeling (Huo et al., 2020). These studies highlight the potential detrimental impact of reduced Mcub expression on immune regulation and for regulating mitochondrial Ca^2+^ uptake. Further work is required to understand the impact that EtOH-induced dysregulation of mitochondrial bioenergetics will have on overall cellular health and senescence.

## Pathways significantly dysregulated at all timepoints

There were multiple gene pathways that were suppressed acutely in response to EtOH and upregulated following abstinence, suggesting a rebound in gene activity during the abstinence period. Across Hallmark, Canonical, and GO analyses, there were consistent patterns of pathway downregulation at both acute time points (PND 35 and PND 46). All of the downregulated pathways are associated with cellular and membrane organization, with additional pathways that involve the regulation of synaptic organization, receptor expression and organization. However, following abstinence there was a robust upregulation of all these downregulated pathways, suggesting protracted cellular and synaptic remodeling events that persist well beyond the acute withdrawal period. It has previously been demonstrated that elevated cytokine expression can result in ECM degradation and further release of cytokines (Nanda et al., 2020). This led to the initial consideration of whether these remodeling events were triggered by immune upregulation, however, that appears not to be the case. No immune-related gene sets were impacted across all 3 time points. However, there were immune-related gene sets downregulated at the later acute time point (PND 46) that were followed by upregulation during abstinence (PND 70), suggesting that the immune related-gene changes did not precede the EtOH-induced cellular remodeling events. ECM and ECM-related genes are heavily involved in these remodeling events and typically undergo heavily regulated degradation and remodeling that are clearly impacted by repeated EtOH exposure. Interestingly, the ECM can recruit cytokines and serve as a depot for pro- and anti-inflammatory cytokines and the degradation process can result in the release of these factors (Zhang et al., 2014, O’Callaghan et al., 2018, Gray et al., 2022). A definitive understanding of the relationship between early suppression of remodeling processes and delayed suppression of immune-related gene pathways requires investigation, however this will be challenging due to the complex bidirectional signaling that occurs across these pathways.

It is hard to dispute the robust transition from acute pathway downregulation during EtOH exposure to identical pathway upregulation during abstinence. The majority of pathways impacted in this temporal manner are involved in cellular remodeling and immune regulation. Interestingly, there is also a very select group of genes that are impacted across multiple pathways that undergo identical shifts in expression. A more thorough analysis of the impact that these select genes changes have on brain function are necessary to uncover their importance in the enduring effects of EtOH. In addition, a more intricate temporal analysis during the abstinence period is necessary to determine how quickly and exactly when and why gene pathway suppression transitions to gene upregulation. Understanding these nuances will provide valuable information regarding an optimal ‘window’ of intervention during the abstinence period to target gene pathways that contribute to neuronal remodeling events and the enduring cognitive changes previously reported.

## Human Homologs of the Candidate Genes

Genes similarly impacted during or after repeated adolescent EtOH exposure and selected GWAS EFO traits were identified. The genes identified have been implicated in sleep quality in humans (sleep_quality EFO 0005272), as biomarkers of inflammation (inflammatory_biomarker_measurement EFO 0004872) and are known to be differentially regulated in light and heavy human drinkers (alcohol_consumption_measurement, EFO_0007878). Interestingly, many of the genes impacted by adolescent EtOH exposure in this rodent model were not in the GWAS data set, suggesting that at least some of the candidate genes in rats are yet to be identified in humans or have yet to be identified as participants in the EFO traits that were selected. These data may have an interesting impact on future studies that are beginning to look more carefully at the contributions of non-neuronal cells and the true complexity of cell variation and cell-to-cell communication necessary for immune modulation, neuronal remodeling, and drinking behavior.

## CONCLUSION

Here we highlight the robust effects of adolescent EtOH exposure on gene pathways involved in cellular remodeling, neuroimmune regulation, sleep, and bioenergetics. Accumulating evidence implicates a role for cellular and circuit remodeling and neuroimmune activation in the emergence of addiction. Here we provide detailed insight into the temporal effects of adolescent EtOH exposure on hippocampal gene expression during acute EtOH exposure and following a period of abstinence, in adulthood. These findings uncover novel pathways that will be an invaluable guide to understanding the mechanisms underlying the acute and long-term effects of adolescent EtOH exposure and provide insight into a potential ‘window’ of opportunity for therapeutic intervention during early abstinence.

## DATA AVAILABILITY

The data presented in this study is the property of the US Federal Government, however, are still available on request from the corresponding author.

## INSTITUTIONAL REVIEW BOARD STATEMENT

The animal study protocol was approved by the Durham Veterans Affairs Institutional Animal Care Committee and Duke University Institutional Animal Care Committee.

## AUTHOR CONTRIBUTIONS

Conceptualization: M-L.R and S.D.M; Data curation: M-L.R and H.G.S; Formal analysis: M-L.R. and A.Q.N; Funding acquisition: M-L.R.; Methodology: M-L.R, S.D.M, H.A.U.M, J.D, A.Q.N; Project administration: M-L.R, A.Q.N; Resources, M-L.R. D.A.P, A.Q.N; Supervision, M-L.R. and A.Q.N, J.D, D.A.P. S.D.M; Validation M-L.R and A.Q.N, D.A.P; Visualization, M-L.R and A.Q.N; Writing—original draft M-L.R. and A.Q.N; Writing—review and editing: A.Q.N, H.A.U.M, H.G.S, S.D.M, J.D, D.A.P, M-L.R. All authors have read and agreed to the published version of the manuscript

## ACKNOWLEDGMENTS

Many thanks to Dr.David Corcoran at the Duke University School of Medicine Center for Genomic and Computational Biology for generating the RNA sequencing data used in this manuscript. Many thanks to the Marshall University Genomics and Bioinformatics Core for help with the data analysis.

## Supplemental Tables

**Table S1. Differential Expression Analysis Results. *HeaderInfo*** sheet contains descriptions of the headers in the PND35, PND46, and PND70 sheets. ***SampleID*** sheet contains information regarding the time points and ethanol or water exposure of each of the samples indicated in the PND35, PND46, and PND70 sheets. ***PND35***, ***PND46***, and ***PND70*** sheets contain the differential expression analysis results for each of the time points. (.XLSX)

**Table S2. Top Pathways from GSEA using Gene Sets of MSigDB Collections for PND 35. *S2A*:** Top 25 Pathways from GSEA (based on p-value) using Gene Sets of MSigDB ImmuneSigDB for PND 35. ***S2B*:** Top 14 Pathways from GSEA (based on p-value) using Gene Sets of MSigDB Hallmark Pathways for PND 35. ***S2C*:** Top 25 Pathways from GSEA (based on p-value) using Gene Sets of MSigDB Canonical Pathways for PND 35. ***S2D*:** Top 25 Pathways from GSEA (based on p-value) using Gene Sets of MSigDB GO Annotations for PND 35. (.DOCX)

**Table S3. Top Pathways from GSEA using Gene Sets of MSigDB Collections for PND 46. *S3A***: Top 25 Pathways from GSEA (based on p-value) using Gene Sets of MSigDB ImmuneSigDB for PND 46. ***S3B*:** Top 15 Pathways from GSEA (based on p-value) using Gene Sets of MSigDB Hallmark Pathways for PND 46. ***S3C*:** Top 25 Pathways from GSEA (based on p-value) using Gene Sets of MSigDB Canonical Pathways for PND 46. ***S3D*:** Top 25 Pathways from GSEA (based on p-value) using Gene Sets of MSigDB GO Annotations for PND 46. (.DOCX)

**Table S4. Top Pathways from GSEA using Gene Sets of MSigDB Collections for PND 70. *S4A***: Top 25 Pathways from GSEA (based on p-value) using Gene Sets of MSigDB ImmuneSigDB for PND 70. ***S4B*:** Top 14 Pathways from GSEA (based on p-value) using Gene Sets of MSigDB Hallmark Pathways for PND 70. ***S4C*:** Top 25 Pathways from GSEA (based on p-value) using Gene Sets of MSigDB Canonical Pathways for PND 70. ***S4D*:** Top 25 Pathways from GSEA (based on p-value) using Gene Sets of MSigDB GO Annotations for PND 70. (.DOCX)

