## Supplemental S1-S4 for "Comparative gene pathway analysis during adolescent binge-EtOH exposure, withdrawal, and following abstinence": Table_S2.docx

**Table S2A. Top 25 Pathways from GSEA (based on p-value) using Gene Sets of MSigDB ImmuneSigDB for PND 35**

| Pathway | pval | padj | log2err | ES | NES | size | Genes | Rank |
| --- | --- | --- | --- | --- | --- | --- | --- | --- |
| GSE15930 NAIVE VS 48H IN VITRO STIM CD8 TCELL DN | 3.87E-06 | 1.86E-02 | 6.11E-01 | 0.431 | 1.811 | 197 | Pdia6 Ssr2 Exosc7 Manf Nans Ube2s Sec13 mrpl11 Ruvbl2 Htatip2 Ndufa8 Thumpd3 Reep5 Abcf1 Ttk Mrpl13 Nutf2 Cops7a Clpp Cox17 Ubfd1 Cdc123 Tfdp1 Fam98a Nop16 Banf1 Pold1 Xpo1 Tardbp Mrps33 Ncaph2 Smc2 Hspd1 Psmd7 Phf5a Elof1 Ubac1 Arl1 Psmb7 Pgk1 Psma1 Hmgb2 Pebp1 Gpn2 Cdc45 Pdap1 Drg1 Psat1 Psmg1 Zranb2 Pdcl3 Farsa Itpa Dpp3 Ssbp1 Prpf31 Ube2m Fasn Mtch2 Cdc34 Psma5 Pfkl Acot9 Pkm Prelid1 | 1 |
| GSE7509 UNSTIM VS FCGRIIB STIM MONOCYTE UP | 1.28E-05 | 1.86E-02 | 5.93E-01 | 0.46 | 1.868 | 152 | Cct3 Sdf2l1 Serpinb9 Lonrf1 Hsd17b10 Eif2a Msrb1 Hsp90ab1 Nmt1 Lman2 Doc2b Cdk2ap1 Cops4 Nagpa Commd6 Sec11a Cript Ogfr Nop16 Tcp1 Cct7 Gabarap Acot13 Cib1 Mrps33 Mrpl15 Ncaph2 Hspd1 RGD1562987 Calm2 Bag1 Mapk8ip1 Cdc42ep5 Pgk1 Ndrg2 Gabarapl2 Psma2 Psat1 Cldnd1 Ssrp1 Lamtor3 Rab1b Sarnp Ppp1r11 Atp6v1e1 Phgdh Ube2a Hdgf Aip Cops5 Ppp6c Tpmt Gde1 Nipsnap3b Hibadh Rhbdd1 Tomm22 Gdi2 Dhcr7 Rab5c Agpat3 Skp1 | 2 |
| GSE18791 CTRL VS NEWCASTLE VIRUS DC 18H UP | 1.32E-05 | 1.86E-02 | 5.93E-01 | 0.411 | 1.722 | 192 | Etv5 Ptk2 Cct3 Nf2 Cct6a Hsd17b10 Mrps6 Ndufa8 Ppil1 Pnp Phax Mtmr12 Bap1 Cops4 Spred1 Rnf14 Jkamp Polr2g Atpsckmt Tfdp1 Zfp185 Timm21 Ndufs2 Impdh2 Cct4 Cdc40 Asb13 Gtf3c2 Ubac1 Pet100 Trmt1 Nelfcd Cetn2 Stam Hvcn1 Snapc5 Daam1 Tchp Serac1 Tfpt Mppe1 Arrb1 Mrpl16 Wdr36 Itpa Nacc2 Dpp3 Elovl1 Endog Tti1 Stradb Suox Ints8 Phf14 Usp38 mrpl24 Aifm1 Rxra Dok2 Tprg1l Acot7 Acss2 Klhdc3 Dcaf13 | 3 |
| GSE24634 NAIVE CD4 TCELL VS DAY5 IL4 CONV TREG DN | 1.91E-05 | 1.86E-02 | 5.76E-01 | 0.41 | 1.72 | 194 | Mcat Cct3 Ciao1 Manf Ahsa1 Ube2s Nf2 Hsd17b10 Ndufa8 Creld2 Ormdl2 Med20 Rab8b Emg1 Ube2k Hspe1 Clpp Wdr18 Dnaja1 Rnf14 Rangap1 Cyb5b Tnfrsf8 Dkc1 Glo1 Fam98a Nop16 Cct7 Acot13 Plpp1 Mars1 Actb Psmc4 Gtdc1 Hspd1 Ube2l3 Tefm Fam136a Gsto1 Coasy Acsl5 Rrp15 Aven Uggt1 Syt11 Phb Kntc1 Cisd1 Psmg1 Cd74 Capza1 Mrpl16 Dpp3 Aimp2 Rbm28 Dohh Aasdhppt Atp6v1c1 C1qbp Hdgf RGD1311164 Arl3 Lars2 Blvra Ppia Rbpj Slc25a5 Rwdd2b Immt Rad1 Gtpbp4 Csnk2a1 Myb | 4 |
| GSE37605 FOXP3 FUSION GFP VS IRES GFP TREG C57BL6 UP | 2.24E-05 | 1.86E-02 | 5.76E-01 | -0.46 | -1.743 | 171 | Sik1 Id1 Arl4d Scarb1 Apold1 Fos Ppp1r15a Gem Dusp1 Adamts1 Tns2 Dll1 Mt1 Ackr3 Padi2 Btg2 Slc16a7 Ptprm Id3 Nr4a1 Cebpb Per1 Ccn2 Atf3 Ier3 H6pd Ipmk Bloc1s3 Eya2 Snx7 Junb Angpt2 Csrnp1 Ch25h Frrs1 Mrm3 Flvcr2 Klhdc10 Pmf1 Tns1 Cnksr3 Trim25 Esam Egr1 Rdh10 Slc25a25 Crem Selenon Akap12 Zfp36 Cmklr1 | 5 |
| GSE29617 CTRL VS DAY7 TIV FLU VACCINE PBMC 2008 DN | 2.29E-05 | 1.86E-02 | 5.76E-01 | 0.431 | 1.768 | 163 | Pdia6 Mettl3 Hyou1 Sdf2l1 Ppib Pdia4 Manf Ca9 Hsp90b1 Spop Homer1 H4f3 Creld2 Borcs8 Ostc Lman2 Sel1l3 Ddost Atxn2 Uqcc1 Gmppb Hax1 Cct7 Siglec8 Ggh Mars1 Aars1 Zwint Cct4 Nudt13 | 6 |
| GSE17721 LPS VS POLYIC 8H BMDC UP | 2.72E-05 | 1.89E-02 | 5.76E-01 | 0.414 | 1.729 | 188 | Pdia6 Lmo4 Hsp90b1 Cct6a Btbd1 Kyat3 ST7 Thumpd3 Rffl Fcgr3a Nsmf Fam162a Gsr Mrpl13 Yipf5 Atxn7l3 Actr1b Lsm4 Cd2bp2 Mllt11 Stk4 Snrpd1 Fcer1g Ehbp1l1 Rnf5 Ncbp1 Zwint Hras Dnajc3 Idnk Tefm Kpna1 Acsl5 Arpc3 Emd Nol11 Chst11 Epha4 Gpn2 Sephs2 Cul1 Nckap1 Fkbp4 Nmnat1 Ltn1 Ppp1r11 Tmem222 Ubl5 Mak16 Cep19 Ncoa5 Grk2 Bnip1 | 7 |
| GSE29617 CTRL VS TIV FLU VACCINE PBMC 2008 DN | 3.28E-05 | 2.00E-02 | 5.57E-01 | 0.42 | 1.745 | 181 | Plac9 Sdf2l1 Tubb5 Pdia4 Hsp90b1 Spop Mrps6 Ing4 Suclg1 Arv1 Ndufa8 Homer1 Borcs8 Ostc Ormdl2 Lman2 Tnnt1 Hdhd2 Tsnax Cdh10 Hax1 Dynll1 Cyb5b Cct7 Siglec8 Ggh Trappc8 Aars1 Rnf5 Zwint Psmd14 Mtif3 Nudt13 Eif2s1 Arl2 Ppp3cb Drg1 Zbtb8os Glt8d1 Oma1 Slc44a1 Sdhaf3 Mtch2 Mea1 Apobec3 Ssb mrpl24 Samm50 Uqcr10 Gnpat Gatd3a Avil Zfp623 Timm8b Tmem260 Psme3ip1 Cct2 Naa20 Armc1 Uqcrq Gnpda1 Scoc Lrif1 Dpy30 C1qb Tmco1 Paqr8 Prdx1 | 8 |
| GSE20715 0H VS 48H OZONE TLR4 KO LUNG DN | 1.15E-04 | 5.99E-02 | 5.38E-01 | 0.398 | 1.66 | 185 | Khdrbs3 Hyou1 Sdf2l1 Spp1 Tubb5 Manf Calr Cyp7b1 Hsph1 Hsp90b1 F3 Bhlhe22 Myc Foxk2 Cop1 Mrps18b Nfil3 Emg1 Hint1 Ywhag Ttk Dnaja1 Rab2a Cct7 Ppa1 Cdk2ap2 Hspd1 Syvn1 Yrdc Gja1 Akr1b8 Cox5a Strap Pgk1 Eif2s1 Cep55 Psmd2 Rras2 Esd Homer2 Ptpn1 Stam Mvd Ckap4 Psma2 Traf3 Eif4e Cacybp Mcfd2 | 9 |
| GSE22886 IGM MEMORY BCELL VS BLOOD PLASMA CELL DN | 1.32E-04 | 5.99E-02 | 5.19E-01 | 0.391 | 1.636 | 193 | Stip1 Ciao1 Tbc1d1 Tm9sf4 Ssr2 Yipf3 Emc9 Cct5 Lap3 Sgk1 Tst Mrps18b Calm3 Actr1a Gmppb Tmx2 Maip1 Ubfd1 Cdc123 Pdhx Vcp Map2k6 Prcp Psmc4 Ufsp2 Aggf1 Sae1 Cars Rpap3 Calm2 Trappc2l Ss18 Aven H2bc12 Cadm1 Cebpd Gpn2 Inpp1 Pds5b Il6st Ywhae Clic4 Rnf34 Tfb2m Snapc5 Rad51ap1 Serinc3 Mpc2 Glt8d1 Rhoq Tfpt Dpp3 Aimp2 Ube2m Acsl4 Gpaa1 Fut8 Uck2 | 10 |
| GSE15930 NAIVE VS 72H IN VITRO STIM TRICHOSTATINA CD8 TCELL UP | 1.75E-04 | 5.99E-02 | 5.19E-01 | -0.435 | -1.652 | 177 | Nr1d2 Ccr9 Eef2k Ncoa3 Usp2 Herpud1 St3gal1 Slc39a4 Klf2 Klhl21 Rbms1 Ackr3 Zfp292 Foxk1 Nr2c1 Reck Ercc2 Scaf8 Tmem268 Cd1d1 Rab33b Spink4 Sh2b3 Grem2 Polg2 Alkbh5 Pomc B4galt1 Ccdc117 Plekha5 Traf2 Atad2b Tcn2 | 11 |
| GSE26727 WT VS KLF2 KO LPS STIM MACROPHAGE UP | 1.81E-04 | 5.99E-02 | 5.19E-01 | -0.434 | -1.65 | 177 | Ppef2 Sik1 Id1 Tob1 St3gal1 Rdh5 Col14a1 Pik3ip1 Fyb1 Gucy2d Tyrp1 Trim66 Tanc2 Bdh1 Per1 Xdh Hsf2bp Scn7a Tbc1d9 Map3k3 Flrt1 Mrc1 Mt2A Casq2 Rab31 Mfng Klf5 Blm Zfp507 Eya2 Acaa1a C2cd2 Srd5a1 Tpm2 Myoc Mid2 Fam3a | 12 |
| GSE27786 BCELL VS ERYTHROBLAST UP | 1.81E-04 | 5.99E-02 | 5.19E-01 | 0.396 | 1.646 | 182 | Nudt18 Nudt3 Manf Lmo4 Entpd5 Lonrf1 Bcl2a1 Ccr5 Ruvbl2 Cyp51 Nabp1 Mllt1 Jagn1 LOC691113 Ebp Osm Mrpl45 Rtca Rhot1 Cox17 Klc1 Bax Timm21 Cct7 Prr3 Traf3ip3 Magi2 Coro1c Pik3cg Mbd1 Trappc3 Pja1 LOC498122 RGD1565059 Fam172a Strap Cnpy3 Cited2 Ywhab Bst2 Tapbp Zadh2 Cluap1 Ddx39b Mfsd3 Dusp16 Fkbp4 Mcfd2 Ltn1 Zfp637 Mlx Tm6sf1 Polr1c Atp5f1a Rbm28 Dcaf15 Nop56 Ap3b1 | 13 |
| GSE13547 CTRL VS ANTI IGM STIM BCELL 12H DN | 1.83E-04 | 5.99E-02 | 5.19E-01 | 0.403 | 1.655 | 163 | Mrpl21 Spg7 Vps4a Prxl2a Irf2bp2 Cct6a Mtap Suclg1 Kbtbd11 Lamp1 Thumpd3 Mrps18b Fam162a Ptpn6 Slco3a1 Prss12 Gpr183 Hax1 Rnaseh2c Cd2bp2 Wdr74 Vps26a Rplp1 Bax Mrpl23 Klhl6 Ei24 Chordc1 Bccip Chchd5 Camsap2 Hspd1 RGD1562987 Acyp1 Mrps18a Erp29 Cpm Elp2 Rrp15 Sephs2 RT1-CE10 Gtf2h5 Fgf13 Psmg1 Dph5 Wdr83os Aup1 Fkbp4 Jakmip1 | 14 |
| GSE20715 0H VS 48H OZONE TLR4 KO LUNG UP | 1.84E-04 | 5.99E-02 | 5.19E-01 | -0.434 | -1.65 | 177 | Nr1d2 Per3 Atp2a3 Nr1d1 Dbp Postn Ier2 B3gnt8 Sun1 Epb41 Cavin2 Dll4 Npnt Ephx1 Hlx Fndc1 Kcnk3 Btg2 Cldn5 Fabp3 Mov10 Rbmx Adra1b Stk17b Mettl27 Hdac11 Pex16 Ucp2 Grem2 Lair1 Stc1 Adcy8 Mylip Lama2 Ntn4 Mpp3 Tap1 Dnah8 Ushbp1 Ddx42 RT1-A2 Gpha2 | 15 |
| GSE16385 UNTREATED VS 12H ROSIGLITAZONE TREATED MACROPHAGE UP | 2.71E-04 | 7.47E-02 | 3.45E-01 | -0.473 | -1.715 | 121 | Ppef2 Krt19 Scrn1 Fbxo30 Slc24a1 Dmrtb1 Trpm4 Arid1a Crb1 Rax Clic2 Foxq1 P2rx1 Ttll4 Foxf1 Smpx Tshz3 Isl2 Cmbl F2rl2 Klhl11 Aldh1l2 Sh2d4b Usp43 Zc2hc1c Rai14 Arhgap20 | 16 |
| GSE14415 INDUCED TREG VS FAILED INDUCED TREG UP | 2.84E-04 | 7.47E-02 | 4.99E-01 | 0.383 | 1.582 | 171 | Arhgap21 Ahsa1 Nans Ube2s Cenpn Sec13 Myc Lap3 Kyat3 Creld2 Psmc3ip Ppil1 Calm3 Morf4l2 Psmd6 Mrps22 Cdk2ap1 Cops4 Mdc1 Rbbp7 Banf1 Spc24 Pold1 Psmc4 Ppa1 Snrpd1 Cars Ube2l3 Calm2 Rfc5 Fam136a Gsto1 Hsd17b12 Pgk1 Snrpa1 Cops3 Cep72 Sgo1 Cisd1 Rad18 Gemin5 | 17 |
| GSE11884 WT VS FURIN KO NAIVE CD4 TCELL DN | 2.90E-04 | 7.47E-02 | 3.35E-01 | -0.487 | -1.738 | 107 | Gpr151 Usp2 Tafa3 Ranbp10 Polr2a Zic3 Tob2 Foxo3 Zxdc Casp8ap2 Slc43a2 Slc16a7 Ube3d Ubn1 Fblim1 Fbxo10 Pitpnm2 Dgkd Wee1 Rprd2 Wnk1 Mex3d Ccdc60 Irx3 Mlxip | 18 |
| GSE15930 NAIVE VS 48H IN VITRO STIM IL12 CD8 TCELL UP | 3.03E-04 | 7.47E-02 | 3.19E-01 | -0.429 | -1.628 | 175 | Nr1d2 Nrep Usp2 St3gal1 Slc39a4 Klf2 Klhl21 Hr Rere Gucy2d Ackr3 Zfp292 Grik1 Ikzf2 Qsox1 Kcnq1 Reck Ercc2 Scaf8 Ltbp1 Pak4 Ephb2 Cd1d1 Rab33b Dpysl3 Mov10 Spink4 Dtx1 Alas2 Spata6 Pip5k1a Mfge8 Tcf7 B4galt1 Plekha5 Tbr1 Casq1 Tcn2 | 19 |
| GSE2935 UV INACTIVATED VS LIVE SENDAI VIRUS INF MACROPHAGE UP | 3.07E-04 | 7.47E-02 | 3.19E-01 | -0.444 | -1.665 | 156 | Sik1 Ccr9 Etv3 St3gal1 Col4a4 Pik3c2a Ephx1 Tet1 Tmem214 Ikzf2 Kdm5a Sh3pxd2a Id3 Lrat St6gal1 Pmepa1 Dnmt3a Pitpnm2 Ism2 Tgfbr3 Gnas Hif1a Dact1 Ppic Igfbp4 Rasgrp1 Gdf10 Plekhg2 C2cd4b Tcf7 Lipe Zic1 Gpr146 N4bp2 Klf13 Tcf20 Ptpn14 Calcrl Foxo1 Birc5 Fam78a Tmem131 Cyth3 Nab2 Eng Tox Rbm38 Abcg1 Ext1 Pdlim4 Tbl1x Satb1 Gtf2ird1 | 20 |
| GSE26669 CTRL VS COSTIM BLOCK MLR CD8 TCELL UP | 4.40E-04 | 9.12E-02 | 4.99E-01 | 0.378 | 1.575 | 183 | Srp14 Serpinb9 Calr Ocel1 Mcts1 Cct6a Kbtbd11 Enoph1 Epsti1 Nabp1 Nol7 Ddn Atxn7l1 Fam98c Dynll1 Lsm4 Tfdp1 Bid Pfdn1 Tcp1 Elmo2 Bcap29 Cpt1c Lynx1 Mrps33 Treml2 Hacd3 Ube2l3 Rfc5 Gsto1 Krt10 Kif18b Eif2s1 Atic Zfp524 Mrpl48 Tapbp Psma2 Hmgcr Rad51ap1 Tiam1 Ccdc157 LOC681282 Farsa Hars1 Fntb Phgdh Zscan22 Psma5 Chmp5 Zfp207 S100a8 Cops5 Ppp6c Ccna2 Eif5a Mtx2 Alg14 Hsd17b7 Nsa2 Chchd1 Pam16 | 21 |
| GSE36826 WT VS IL1R KO SKIN STAPH AUREUS INF UP | 4.68E-04 | 9.12E-02 | 3.93E-01 | 0.376 | 1.57 | 187 | Etv5 Spp1 Tubb5 Dhcr24 Tubb2a Sec13 Ube2c Pstpip1 Nmt1 Actr1a Psmd6 Mipep Arf3 Ptpn6 Cops7a Cox17 Arpc5l Csrp1 Xpo1 Psmc4 Zwint Canx Hspd1 Dusp6 Cct4 Cox5a Ncapg Mcub Psmb7 Emc8 Eif2s1 Cep55 Fanci Hjurp Cox5b Snap29 Cdc45 Atic Lamtor5 Rrp7a P2ry6 Aarsd1 Dpp3 Fasn Got1 Ipo4 Acot9 Blvra Apobec3 Sh3bgrl3 Clta Ranbp1 Ran Arf1 Tpmt Cemip2 Procr Acot7 Gpi Ifi35 Tpx2 Racgap1 Nisch Psmd8 Ndc1 Uqcrq Otulinl Atp2a2 Wdr12 | 22 |
| GSE2770 IL12 AND TGFB ACT VS ACT CD4 TCELL 6H DN | 4.95E-04 | 9.12E-02 | 3.79E-01 | 0.379 | 1.574 | 180 | Pdia6 Stip1 Cct3 Arl15 Ptges3 Manf Ahsa1 Nans Eif1 Rnls Xbp1 Nudt15 Hspa5 Mtmr1 Snrpg Nabp1 Rab29 Gmppb Grpel1 Fbxo4 Cox17 Arpc5l Nrp1 Nop16 Cct7 Ppa1 Faslg Pa2g4 Fes Ube2l3 Cd53 Yrdc Snca Gne Tmem30b Egfl6 Nucb1 Strap Dnajb6 Ak4 Rras2 Tfg Pim1 Ube3c Ptpn1 Rnf34 Prnp Ddx39b Mmd Fkbp4 Cflar Dctpp1 Nacc2 Pcsk5 | 23 |
| GSE40277 EOS AND LEF1 TRANSDUCED VS GATA1 AND SATB1 TRANSDUCED CD4 TCELL UP | 4.99E-04 | 9.12E-02 | 2.46E-01 | -0.415 | -1.59 | 187 | Igsf9 Dbp Lnx2 Fbxo30 Eef2k Gch1 Ncoa3 Ca8 Greb1 Ddc Nsmaf Pard6b Myo1c Mpzl2 Adgre5 Unc119 Foxo3 Klf9 Wnt3 Pmepa1 Csrp2 Gsap Ssh1 Bhlhe40 Purg Gm2a Ccn4 Abcc1 Rims3 Lipe Cish Arhgap20 Lamc1 Ttc39b Ilvbl Zc3h12a Phtf2 Nfkb1 Bnip5 Atxn1 RT1-A2 | 24 |
| GSE37605 TREG VS TCONV NOD FOXP3 FUSION GFP UP | 5.39E-04 | 9.12E-02 | 2.42E-01 | -0.457 | -1.663 | 125 | Arl4d Ier2 Fzd4 Apold1 Fos Smad7 Ppp1r15a Dusp1 Adamts1 Polr2a Optc Mt1 Btg2 Tbx3 Id3 Nr4a1 Cebpb Gabrd Foxs1 Birc3 Atf3 Ier3 Oasl Trib1 Grm8 Itga5 Gsto2 Junb Flvcr2 Cryga Egr1 Ust Slc25a25 Dusp8 Akap12 Zfp36 | 25 |

**Table S2B. Top 14 Pathways from GSEA (based on p-value) using Gene Sets of MSigDB Hallmark Pathways for PND 35**

| Pathway | pval | padj | log2err | ES | NES | size | Genes | Rank |
| --- | --- | --- | --- | --- | --- | --- | --- | --- |
| HALLMARK MYC TARGETS V1 | 1.01E-06 | 5.04E-05 | 6.44E-01 | 0.443 | 1.857 | 196 | Cct3 Ptges3 Exosc7 Cct5 Myc Ruvbl2 Hsp90ab1 Snrpg Mrps18b Hnrnpa2b1 Hspe1 Hnrnpr Ctps1 Stard7 Tfdp1 Glo1 Nop16 Tcp1 Mrpl23 Cct7 Xpo1 Tardbp Psmc4 Snrpa Snrpd1 Ncbp1 Pa2g4 Psmd14 Impdh2 Canx Hspd1 Psmd7 Cct4 Ube2l3 Ube2e1 Eif2s2 Cox5a Pgk1 Psma1 Eif2s1 Rad23b Snrpa1 Txnl4a Cdc45 Hnrnpc Cul1 Ywhae Phb Psma2 Vdac3 Rps2 Eif4e Aimp2 Ssbp1 Prpf31 Nap1l1 C1qbp Pcbp1 Nop56 Hdgf Eif1a Snrpd3 Ppia Nolc1 Ssb Srsf2 Cops5 Ranbp1 Ldha Ran Ccna2 Pabpc1 Psmc6 Eif1ax Lsm2 Cct2 H2az1 Psmb3 Ddx21 Eprs U2af1 Smarcc1 Psmd8 Rpl14 | 1 |
| HALLMARK HYPOXIA | 9.47E-05 | 2.37E-03 | 5.38E-01 | -0.442 | -1.686 | 182 | Scarb1 Lxn Pgam2 Pdgfb Ccn1 Fos Ppp1r15a Dusp1 Srpx Tgfbi Mt1 Ackr3 Foxo3 Zfp292 Efna1 Cavin3 Ccn2 Csrp2 Stc2 Siah2 Atf3 Stbd1 Ier3 Ccng2 Ets1 Mxi1 Bhlhe40 Mt2A Dcn Kdm3a Cdkn1c Pam Pgf Dtna Stc1 Noct Atp7a Ppargc1a Slc2a1 Gpc4 Ilvbl Kif5a Akap12 Zfp36 Pgm1 Ndst1 Cp Col5a1 Nedd4l | 2 |
| HALLMARK ESTROGEN RESPONSE EARLY | 1.43E-04 | 2.39E-03 | 5.19E-01 | -0.438 | -1.669 | 180 | Krt15 Krt19 Scarb1 Klf10 Mlph Hes1 Fos Calcr Tob1 Slc7a5 Greb1 Tmem164 Hr Unc119 Aff1 Fkbp5 Clic3 Pdlim3 Flnb Klf4 Slc7a2 Stc2 Siah2 Slc22a5 Igf1r Slc19a2 Rps6ka2 Frk Tsku Bhlhe40 Rhobtb3 Igfbp4 Fhl2 Rab31 Rasgrp1 Olfml3 Cish B4galt1 Slc27a2 Gab2 Slc2a1 Elf1 Mreg Calb2 Podxl Inhbb | 3 |
| HALLMARK TNFA SIGNALING VIA NFKB | 4.17E-04 | 5.21E-03 | 2.70E-01 | -0.423 | -1.608 | 178 | Sik1 Ier2 Jag1 Klf10 Gch1 Hes1 Ccn1 Fos Ppp1r15a Gfpt2 Gem Dusp1 Klf2 Traf1 Ackr3 Btg2 Nfe2l2 Klf9 Birc2 Efna1 Nr4a1 Klf4 Cebpb Per1 Pmepa1 Birc3 Atf3 Ier3 Bhlhe40 Trib1 Sphk1 Cxcl6 | 4 |
| HALLMARK UV RESPONSE DN | 7.85E-04 | 7.85E-03 | 1.99E-01 | -0.436 | -1.614 | 143 | Nr1d2 Id1 Dbp Ccn1 Smad7 Dusp1 Irs1 Mt1 Ptpn21 Ptprm Scaf8 Ltbp1 Igf1r Tgfbr3 Bhlhe40 Fbln5 Slc7a1 Fhl2 Has2 Pik3r3 Inpp4b Rgs4 Atp2b4 Abcc1 Prdm2 Lamc1 Slc22a18 Col1a1 Ddah1 Nfkb1 Cdk13 Atxn1 Dlg1 Plcb4 Col3a1 Akt3 Cdon Kcnma1 Sfmbt1 Anxa4 Ica1 Met | 5 |
| HALLMARK EPITHELIAL MESENCHYMAL TRANSITION | 1.08E-03 | 8.96E-03 | 1.67E-01 | -0.407 | -1.552 | 184 | Postn Ccn1 Gem Tgfbi Lum Eln Anpep Mgp Timp3 Qsox1 Tnfrsf11b Lgals1 Cald1 Lama1 Ecm1 Pmepa1 Ccn2 Tgfbr3 Dpysl3 Fbn2 Fbln5 Mylk Itga5 Cxcl6 Fstl1 Dcn Col4a1 Igfbp4 Fn1 Efemp2 Vcan Rgs4 Prrx1 Abi3bp Tpm2 Fmod Sdc1 Nt5e Lama2 Lamc1 Copa P3h1 Col6a3 Col1a1 Flna Myl9 Mfap5 Tagln Col5a1 Igfbp2 Col3a1 Dst Itga2 Sntb1 Pvr Cd44 Fap Acta2 Mest Sgcb Pcolce Gadd45a Fuca1 Pdlim4 Foxc2 Itgb5 Col7a1 Itgav Colgalt1 Bgn Msx1 | 6 |
| HALLMARK MTORC1 SIGNALING | 3.60E-03 | 2.57E-02 | 1.40E-01 | 0.343 | 1.436 | 193 | Stip1 Sdf2l1 Calr Xbp1 Elovl5 Hsp90b1 Dhcr24 P4ha1 Hspa5 Slc1a4 Uchl5 Cct6a Cyp51 Nfil3 Pnp Nmt1 Ebp Hspe1 Gsr Arpc5l Sec11a Cyb5b Ube2d3 Scd2 Mllt11 Psmc4 Ppa1 Psmd14 Canx Hspd1 Actr3 Sla Eif2s2 Skap2 Lta4h Pgk1 Slc37a4 Ak4 Fads2 Ddit3 Atp5mc1 Pdap1 Psat1 Hmgcr Psmg1 Qdpr Cacybp Rit1 Phgdh Rpn1 Got1 Ccng1 Dapp1 Pfkl Acsl3 Ppia Tmem97 Cops5 Asns Ssr1 Ldha Immt Psmc6 Slc9a3r1 Gpi Sc5d Uso1 Tfrc Gtf2h1 Dhcr7 Serpinh1 Egln3 Idh1 Eprs Ifi30 Rab1a Ykt6 Tuba4a Atp2a2 | 7 |
| HALLMARK TGF BETA SIGNALING | 5.40E-03 | 3.37E-02 | 7.95E-02 | -0.516 | -1.636 | 52 | Id1 Klf10 Smad7 Ppp1r15a Id3 Pmepa1 Smurf1 Cdkn1c Rab31 Junb Skil | 8 |
| HALLMARK WNT BETA CATENIN SIGNALING | 8.18E-03 | 4.55E-02 | 6.53E-02 | -0.556 | -1.642 | 36 | Jag1 Fzd1 Maml1 Dll1 Ppard Dvl2 Hdac5 Hdac11 Tcf7 Ptch1 Notch1 | 9 |
| HALLMARK OXIDATIVE PHOSPHORYLATION | 1.17E-02 | 5.68E-02 | 7.74E-02 | 0.323 | 1.352 | 193 | Ndufa5 Hsd17b10 mrpl11 Suclg1 Ndufa8 Mrps11 Glud1 Mrps22 Pdhb Idh3b Atp6v1g1 Grpel1 Rhot1 Cox17 Retsat Mrps30 Pdhx Bax Mpc1 Mrpl15 Ndufs2 Atp1b1 Cox5a Dld Cox5b Idh3a Atp5mc1 Etfb Ndufs8 Vdac3 Iscu Atp6v1e1 Atp5f1a Ndufa3 Dlst Atp6v1c1 Ndufa2 Acaa2 Aifm1 Slc25a5 Uqcr10 Atp5me Ldha Atp6v1f Ndufv2 Immt Dlat Tomm22 Mtx2 Gpi Timm8b Ndufv1 Cyb5a Cs Atp6v0c Idh1 Uqcrq Ndufb7 Slc25a20 | 10 |
| HALLMARK UNFOLDED PROTEIN RESPONSE | 1.25E-02 | 5.68E-02 | 6.93E-02 | 0.374 | 1.446 | 108 | Pdia6 Hyou1 Calr Xbp1 Hsp90b1 Tubb2a Hspa5 Slc1a4 Lsm1 Dnaja4 Cnot2 Nabp1 Sec11a Lsm4 Nfyb Dkc1 Banf1 Atp6v0d1 Zbtb17 Dnajc3 Eif2s1 Imp3 Atf4 Dnajb9 Psat1 Eif4e Eif4g1 Ttc37 Gemin4 Nop56 Nolc1 Vegfa Asns Ssr1 | 11 |
| HALLMARK COMPLEMENT | 1.84E-02 | 7.65E-02 | 6.01E-02 | 0.327 | 1.345 | 166 | Dgkg Kcnip2 F3 Gmfb Hspa5 Itgam Msrb1 Dgkh Lap3 Calm3 Olr1 Gng2 Scg3 Rnf4 Csrp1 Prcp Fcer1g Xpnpep1 Pik3cg Dusp6 S100a9 Calm2 Dusp5 Cpm Lta4h Gca Pim1 Was Casp1 Src Lyn Ehd1 Actn2 Ctss Hspa1a C1s Pik3r5 Car2 Mt3 Apobec3 Lck Plaur | 12 |
| HALLMARK ESTROGEN RESPONSE LATE | 2.51E-02 | 9.67E-02 | 3.42E-02 | -0.354 | -1.346 | 177 | Krt19 Gper1 Scarb1 Fos Calcr Tob1 Slc7a5 Slc29a1 Hr Aff1 Fkbp5 Clic3 Dnajc12 Pdlim3 Emp2 Flnb Klf4 Ass1 Siah2 Slc22a5 Rps6ka2 Frk Igfbp4 Cdc6 Rab31 Atp2b4 Cish Impa2 Slc27a2 Pdcd4 Llgl2 Gins2 Chst8 Gale Zfp36 Kcnk5 Rbbp8 Stil Sord Rnaseh2a Myof Gfus Hmgcs2 Tfap2c St14 Nab2 Ugdh St6galnac2 Cd44 Ret Aldh3a2 Mest Foxc1 Cacna2d2 Mdk Cd9 | 13 |
| HALLMARK KRAS SIGNALING DN | 4.04E-02 | 1.44E-01 | 2.73E-02 | -0.359 | -1.326 | 141 | Krt15 Oxt Cacna1f Pax4 Nphs1 Ngb Cpeb3 Btg2 Itih3 Tfcp2l1 Col2a1 Slc16a7 Slc38a3 Lfng Shox2 Pax3 Fgf22 Cyp11b2 Smpx Gprc5c Tfap2b Lypd3 Dtnb Thnsl2 Zc2hc1c Skil | 14 |

**Table S2C. Top 25 Pathways from GSEA (based on p-value) using Gene Sets of MSigDB Canonical Pathways for PND 35**

| Pathway | pval | padj | log2err | ES | NES | size | Genes | Rank |
| --- | --- | --- | --- | --- | --- | --- | --- | --- |
| PID RHODOPSIN PATHWAY | 2.56E-11 | 4.74E-08 | 8.63E-01 | -0.926 | -2.384 | 19 | Grk1 Pde6g Slc24a1 Rdh5 Guca1b Sag Guca1a Pde6a Rgs9 Gucy2d Rpe65 Cnga1 Lrat Gngt1 | 1 |
| NABA CORE MATRISOME | 8.67E-11 | 8.02E-08 | 8.39E-01 | -0.528 | -2.055 | 220 | Tinagl1 Postn Col26a1 Impg2 Esm1 Bmper Ccn1 Col20a1 Col4a4 Dpt Sspo Nyx Srpx Col14a1 Tgfbi Lum Npnt Tnn Optc Eln Fndc1 Mgp Lamb1 Col4a3 Gldn Col2a1 Crispld2 Impg1 Lama5 Lama1 Cilp Prelp Ecm1 Thbs4 Fndc7 Ccn2 Ush2a Ltbp1 Igfbp7 Hmcn1 Tsku Col6a6 Fbn2 Fbln5 Ccn4 Egflam Dcn Col4a1 Igfbp4 Sbspon Fn1 Efemp2 Vcan Vwa5b1 Lgi2 Abi3bp Mfge8 Fmod Fras1 Dmp1 Col4a6 Lama2 Lamc1 Ntn4 Ntng2 Ibsp Col6a3 Col1a1 Matn4 Kcp Mfap5 Col4a5 Srgn Col5a1 Gas6 Igfbp2 Vtn Ltbp3 Nid1 Col3a1 Sned1 | 2 |
| PID CONE PATHWAY | 6.29E-09 | 3.88E-06 | 7.62E-01 | -0.9 | -2.285 | 18 | Grk1 Gnat2 Arr3 Gngt2 Rdh5 Gnb3 Guca1b Guca1a Rgs9 Gucy2d Rpe65 Pde6c Lrat | 3 |
| WP CIRCADIAN RHYTHM RELATED GENES | 5.14E-08 | 2.38E-05 | 7.20E-01 | -0.52 | -1.978 | 178 | Nr1d2 Sik1 Per3 Ciart Nr1d1 Dbp Pax4 Klf10 Tnfrsf11a Aanat Usp2 Bhlhe41 Tph2 Drd4 Ddc Nr2f6 Kcnh7 Tph1 Klf9 Kdm5a Id4 Drd1 Id3 Adora2a Cavin3 Per1 Kmt2a Nr1h3 Hdac3 Ube3a Zfhx3 Bhlhe40 Sirt1 Sin3a Rorb Tyms Crx Avp Noct Per2 Hs3st2 Ppargc1a Arnt Prokr2 Egr1 Prkg2 Pspc1 Setx | 4 |
| REACTOME VISUAL PHOTOTRANSDUCTION | 1.19E-06 | 4.40E-04 | 6.44E-01 | -0.612 | -2.07 | 76 | Grk1 Cngb1 Bco2 Pde6g Slc24a1 Rdh5 Guca1b Sag Guca1a Pde6a Rgs9 Rbp3 Gucy2d Lrp12 Rpe65 Opn1sw Cnga1 Lrat Gngt1 | 5 |
| REACTOME THE PHOTOTRANSDUCTION CASCADE | 3.51E-06 | 1.08E-03 | 6.27E-01 | -0.751 | -2.107 | 28 | Grk1 Cngb1 Pde6g Slc24a1 Guca1b Sag Guca1a Pde6a Rgs9 Gucy2d Cnga1 Gngt1 | 6 |
| NABA ECM GLYCOPROTEINS | 4.42E-06 | 1.17E-03 | 6.11E-01 | -0.496 | -1.848 | 151 | Tinagl1 Postn Bmper Ccn1 Dpt Sspo Srpx Tgfbi Npnt Tnn Eln Fndc1 Mgp Lamb1 Gldn Crispld2 Lama5 Lama1 Cilp Ecm1 Thbs4 Fndc7 Ccn2 Ush2a Ltbp1 Igfbp7 Hmcn1 Tsku Fbn2 Fbln5 Ccn4 Egflam Igfbp4 Sbspon Fn1 Efemp2 Vwa5b1 Lgi2 Abi3bp Mfge8 Fras1 Dmp1 Lama2 Lamc1 Ntn4 Ntng2 Ibsp Matn4 Kcp Mfap5 Gas6 Igfbp2 Vtn Ltbp3 Nid1 Sned1 | 7 |
| REACTOME MITOCHONDRIAL TRANSLATION | 8.63E-06 | 2.00E-03 | 5.93E-01 | 0.52 | 1.949 | 89 | Mrpl21 Mrps10 Mrps6 mrpl11 Mrps18c Mrps11 Mrps18b Mrps35 Mrps22 Mrpl13 Mrpl49 Mrps30 Mrps27 Gfm1 Mrpl23 Mrps33 Mrpl15 Mtif3 Mrps18a Gadd45gip1 Mrps26 Mrps16 Mrps17 Mrpl48 Mtfmt Mrpl17 Mrpl40 Mrpl16 Mrps7 Mrps36 mrpl24 Mrpl20 | 8 |
| WP CANONICAL AND NONCANONICAL NOTCH SIGNALING | 1.76E-05 | 3.21E-03 | 5.76E-01 | -0.739 | -2.042 | 26 | Jag1 Hes1 Dll4 Maml1 Dll1 Dlk1 Dll3 Rras Dner Cntn6 Notch3 Notch1 | 9 |
| REACTOME HSP90 CHAPERONE CYCLE FOR STEROID HORMONE RECEPTORS SHR | 1.78E-05 | 3.21E-03 | 5.76E-01 | 0.642 | 2.12 | 45 | Stip1 Ptges3 Tubb2a Ar Tuba8 Dctn5 Dnaja4 Tubb4b Hsp90ab1 Actr1a Capza2 Tubb4a Dnaja1 Dynll1 Dctn4 Capzb Dync1i2 Dnaja2 Tubb3 Fkbp4 Capza1 Dctn2 Hspa1a | 10 |
| REACTOME COOPERATION OF PREFOLDIN AND TRIC CCT IN ACTIN AND TUBULIN FOLDING | 1.91E-05 | 3.21E-03 | 5.76E-01 | 0.725 | 2.122 | 26 | Cct3 Tubb2a Cct5 Tuba8 Cct6a Tubb4b Tubb4a Pfdn1 Tcp1 Cct7 Pfdn4 Actb Cct4 Tubb3 Pfdn6 Cct8 Tuba1b Cct2 Tuba4a Cct6b Tubb2b | 11 |
| PID ERBB1 INTERNALIZATION PATHWAY | 2.61E-05 | 4.03E-03 | 5.76E-01 | 0.64 | 2.06 | 40 | Ptk2 Chmp3 Pik3r2 Sh3gl2 Arhgef7 Spry2 Pik3cd Ube2d1 Ube2d3 Epn1 Cdc42 Hras Ube2d2 Eps15 Dnm1 Src Pik3r1 | 12 |
| REACTOME SENSORY PERCEPTION | 5.73E-05 | 8.16E-03 | 5.57E-01 | -0.467 | -1.741 | 149 | Grk1 Kcnq4 Cngb1 Bco2 Pde6g Slc24a1 Rdh5 Guca1b Myo1c Sag Guca1a Pde6a Rgs9 Rbp3 Gucy2d Lrp12 Rpe65 Clic5 Opn1sw Reep4 Twf2 Cnga1 Lrat Reep6 Gngt1 Ctbp2 Actg1 Ripor2 Fscn2 Myo7a | 13 |
| NABA BASEMENT MEMBRANES | 6.44E-05 | 8.51E-03 | 5.38E-01 | -0.657 | -1.964 | 38 | Col4a4 Npnt Lamb1 Col4a3 Lama5 Lama1 Ush2a Hmcn1 Col6a6 Col4a1 Col4a6 Lama2 Lamc1 Ntn4 Ntng2 Col6a3 Col4a5 Nid1 | 14 |
| REACTOME COPI INDEPENDENT GOLGI TO ER RETROGRADE TRAFFIC | 1.01E-04 | 1.25E-02 | 5.38E-01 | 0.608 | 1.987 | 43 | Pla2g6 Tubb2a Tuba8 Dctn5 Tubb4b Actr1a Galnt1 Capza2 Tubb4a Dynll1 Dctn4 Capzb Dync1i2 Bicd2 Tubb3 Capza1 Dctn2 Rab3gap2 Dync1i1 Rab6a Tuba1b Agpat3 Tuba4a Tubb2b | 15 |
| REACTOME ECM PROTEOGLYCANS | 1.25E-04 | 1.44E-02 | 5.19E-01 | -0.563 | -1.879 | 70 | Col4a4 Lum Tnn Lamb1 Col4a3 Col2a1 Lama5 Lama1 Col6a6 Dcn Col4a1 Fn1 Vcan Fmod Dmp1 Col4a6 Lama2 Lamc1 Ibsp Col6a3 Col1a1 Matn4 Col4a5 Col5a1 Vtn Col3a1 Itga2 Col9a1 Itgb5 Itga8 Itgav Bgn Col9a3 Lama4 | 16 |
| REACTOME FORMATION OF TUBULIN FOLDING INTERMEDIATES BY CCT TRIC | 2.27E-04 | 2.32E-02 | 5.19E-01 | 0.738 | 2.002 | 19 | Cct3 Tubb2a Cct5 Tuba8 Cct6a Tubb4b Tubb4a Tcp1 Cct7 Cct4 Tubb3 Cct8 Tuba1b Cct2 Tuba4a Cct6b Tubb2b | 17 |
| REACTOME MHC CLASS II ANTIGEN PRESENTATION | 2.29E-04 | 2.32E-02 | 5.19E-01 | 0.457 | 1.747 | 101 | Ap1m1 Tubb2a Sh3gl2 Tuba8 Dctn5 Sec13 Ap2a2 Tubb4b Actr1a Capza2 Sptbn2 Tubb4a Ap2m1 Dynll1 Actr1b Klc1 Dctn4 Ap1s3 Capzb Dync1i2 Rilp Canx Osbpl1a Ctsf Tubb3 Dnm1 Cd74 Capza1 Ctss Dctn2 Kif3b Clta Sec23a Arf1 Dync1i1 Kifap3 Ap2b1 RT1-DOb Racgap1 Tuba1b Sec24c Ifi30 Kif2c Tuba4a RT1-Ba Ap1s2 Tubb2b | 18 |
| NABA COLLAGENS | 2.51E-04 | 2.32E-02 | 3.79E-01 | -0.631 | -1.902 | 40 | Col26a1 Col20a1 Col4a4 Col14a1 Col4a3 Col2a1 Col6a6 Col4a1 Col4a6 Col6a3 Col1a1 Col4a5 Col5a1 Col3a1 Col24a1 Col9a1 Col7a1 Col22a1 Col9a3 Col12a1 Col17a1 Col13a1 Col6a2 Col18a1 Col8a1 Col9a2 | 19 |
| REACTOME COLLAGEN CHAIN TRIMERIZATION | 2.51E-04 | 2.32E-02 | 3.79E-01 | -0.631 | -1.902 | 40 | Col26a1 Col20a1 Col4a4 Col14a1 Col4a3 Col2a1 Col6a6 Col4a1 Col4a6 Col6a3 Col1a1 Col4a5 Col5a1 Col3a1 Col24a1 Col9a1 Col7a1 Col22a1 Col9a3 Col12a1 Col17a1 Col13a1 Col6a2 Col18a1 Col8a1 Col9a2 | 20 |
| WP NEOVASCULARISATION PROCESSES | 3.03E-04 | 2.67E-02 | 3.45E-01 | -0.638 | -1.885 | 36 | Jag1 Pdgfb Dll4 Ephb4 Ephb2 Hif1a Acvrl1 Smad2 Mmp9 Notch3 Notch1 Nfkb1 Kitlg | 21 |
| REACTOME UNFOLDED PROTEIN RESPONSE UPR | 3.30E-04 | 2.78E-02 | 4.99E-01 | 0.481 | 1.789 | 85 | Pdia6 Add1 Hyou1 Dnajb11 Exosc7 Calr Xbp1 Ddx11 Hsp90b1 Hspa5 Extl1 Mbtps2 Nfyb Atp6v0d1 Zbtb17 Syvn1 Dnajc3 Eif2s2 Eif2s1 Ddit3 Gsk3a Atf4 Dnajb9 | 22 |
| KEGG ECM RECEPTOR INTERACTION | 3.48E-04 | 2.80E-02 | 3.11E-01 | -0.525 | -1.768 | 74 | Col4a4 Tnn Lamb1 Col2a1 Lama5 Lama1 Thbs4 Col6a6 Itga10 Itga5 Col4a1 Fn1 Col4a6 Sdc1 Lama2 Lamc1 Ibsp Col6a3 Col1a1 Itga11 Hmmr Col5a1 Vtn Col3a1 Itga2 Itga1 Cd44 Itgb5 Itga8 Itgav Vwf Lama4 Itga3 | 23 |
| REACTOME SLC MEDIATED TRANSMEMBRANE TRANSPORT | 3.67E-04 | 2.83E-02 | 2.86E-01 | -0.414 | -1.599 | 202 | Slc15a1 Ctns Slc30a5 Slc24a1 Slc7a5 Ddi2 Slc8b1 Slc29a1 Slc39a4 Slc29a4 Slc8a3 Slc30a7 Slco4a1 Slc43a2 Slc16a7 Slc38a3 Slc11a2 Slc4a1 Slc35b2 Slc17a6 Slc12a6 Slc5a8 Slc7a2 Slc22a5 Slc7a1 Slc38a1 Slc44a3 Slc43a1 Avp Slc38a5 Slc22a8 Slc22a3 Slc1a5 Slco2a1 Slc2a1 Slc5a7 Slc30a1 Slc3a1 Slc22a18 Slc16a10 Slc26a1 Slc39a7 Emb Slc7a3 Slc26a4 Slc2a4 Slc13a5 Slc39a6 Cp Slc5a6 Slc22a6 Slc25a10 Slc22a2 Slc4a10 Slc12a2 Slc6a6 Slc45a3 Slc41a2 Slc16a3 Slc39a14 Slc36a1 Slc13a4 | 24 |
| NABA PROTEOGLYCANS | 4.46E-04 | 3.14E-02 | 2.86E-01 | -0.661 | -1.868 | 29 | Impg2 Esm1 Nyx Lum Optc Impg1 Prelp Dcn Vcan Fmod | 25 |

**Table S2D. Top 25 Pathways from GSEA (based on p-value) using Gene Sets of MSigDB GO Annotations for PND 35**

| Pathway | pval | padj | log2err | ES | NES | size | Genes | Rank |
| --- | --- | --- | --- | --- | --- | --- | --- | --- |
| GOBP SENSORY PERCEPTION OF LIGHT STIMULUS | 6.47E-23 | 3.17E-19 | 1.24E+00 | -0.687 | -2.603 | 171 | Ppef2 Grk1 Cacna2d4 Cacna1f Cngb1 Rpgrip1 Gnat2 Cabp4 Arr3 Impg2 Rd3 Cplx4 Pde6g Slc24a1 Cplx3 Rdh5 Trpm1 Rp1l1 Rom1 Nyx Dll4 Tgfbi Lum Guca1b Cnnm4 Tulp1 Crb1 Lrit3 Guca1a Pde6a Rgs9 Rbp3 Gucy2d Kifc3 Unc119 Rpe65 Rax Eya3 Clic5 Timp3 Opn1sw Pde6c Col2a1 Aipl1 Pdc Impg1 Cnga1 Lrat Reep6 Prcd Ush2a Hmcn1 Crybg3 Rgs16 Zic2 Eml2 Adgrv1 Grm8 Rs1 Rorb Atxn7 Crx Fscn2 Myo7a Cryga | 1 |
| GOBP SENSORY PERCEPTION | 1.91E-16 | 4.67E-13 | 1.05E+00 | -0.508 | -2.09 | 404 | Ppef2 Grk1 Cacna2d4 Oxt Lxn Cacna1f Kcnq4 Fzd4 Cngb1 Rpgrip1 Gnat2 Cabp4 Myo7b Arr3 Impg2 Rd3 Cplx4 Pde6g Slc24a1 Cplx3 Rdh5 Trpm1 Rp1l1 Rom1 Nyx Ttc8 Dll4 Tgfbi Lum Hoxd1 Guca1b Nr2f6 Cnnm4 Tulp1 Crb1 Lrit3 Guca1a Pde6a Rgs9 Rbp3 Gucy2d Kifc3 Unc119 Pkhd1l1 Rpe65 Rax Eya3 Itpr3 Clic5 Timp3 Tbx1 Opn1sw Col4a3 Pde6c Ntsr1 Col2a1 Aipl1 Kcnq1 Pdc Adora2a Impg1 Cnga1 Lrat Reep6 Prcd Ush2a Pax3 Pgap1 Hmcn1 Crybg3 Rgs16 Zic2 Gnas Eml2 Pou4f1 Adgrv1 Grm8 Arrb2 Srrm4 Asic2 Rs1 Rorb Cdc14a Syt10 Atxn7 Ano1 Ripor2 Crx Scn9a Fscn2 Myo7a Cryga Six1 B3gnt2 Grik2 Opa1 Chrna4 Cln6 Myh14 Lrig2 P2rx4 Cngb3 Trpv1 Rpgr Nxnl2 Dcdc2 Col1a1 Rdh10 Casp3 Lrrc51 Slc26a4 P2rx7 Ror1 Cep250 Bbs7 Nlgn2 Otof Usp53 Dram2 F2r Calhm1 Oprk1 Birc5 | 2 |
| GOBP DETECTION OF LIGHT STIMULUS | 7.44E-13 | 1.22E-09 | 9.21E-01 | -0.765 | -2.487 | 60 | Grk1 Cacna2d4 Cacna1f Cngb1 Gnat2 Cabp4 Pde6g Gngt2 Rom1 Guca1b Tulp1 Sag Crb1 Guca1a Pde6a Gucy2d Unc119 Rpe65 Opn1sw Pde6c Aipl1 Pdc Cnga1 Reep6 Gngt1 Asic2 Rs1 Cds1 Gpr88 Pitpnm1 | 3 |
| GOBP RESPONSE TO LIGHT STIMULUS | 4.49E-12 | 5.50E-09 | 8.87E-01 | -0.517 | -2.061 | 279 | Sik1 Per3 Grk1 Cacna2d4 Cacna1f Cngb1 Gnat2 Cabp4 Aanat Usp2 Ctns Fbxl17 Pde6g Slc24a1 Fos Gngt2 Trpm1 Dusp1 Rom1 Guca1b Nr2f6 Tulp1 Sag Crb1 Guca1a Pde6a Gucy2d Unc119 Rpe65 Cpt1b Opn1sw Pde6c Map3k4 Aipl1 Drd1 Pdc Cdc25a Aurkb Cnga1 Per1 Reep6 Ercc2 Gngt1 Nfatc4 Nedd4 Hif1a Bhlhe40 Asic2 Rs1 Sirt1 Hyal1 Atr Cds1 Eif2ak4 Gpr88 Xpc Primpol Pias1 Bak1 Pitpnm1 Crip1 Mmp9 Polk Per2 | 4 |
| GOCC PHOTORECEPTOR OUTER SEGMENT | 1.71E-11 | 1.68E-08 | 8.63E-01 | -0.72 | -2.416 | 72 | Ppef2 Grk1 Cacna1f Cngb1 Gnat2 Arr3 Rd3 Pde6g Rp1l1 Rom1 Guca1b Mertk Tulp1 Sag Crb1 Guca1a Pde6a Gucy2d Cdhr1 Opn1sw Myrip Pdc Impg1 Cnga1 Prcd Gngt1 Iqcb1 Mak Vcan Myo7a Ocrl Nxnl1 Cngb3 Rpgr | 5 |
| GOCC 9PLUS0 NON MOTILE CILIUM | 6.64E-11 | 5.42E-08 | 8.39E-01 | -0.639 | -2.271 | 106 | Ppef2 Grk1 Cacna1f Cngb1 Rpgrip1 Gnat2 Arr3 Rd3 Pde6g Rp1l1 Rom1 Guca1b Mertk Tulp1 Sag Crb1 Guca1a Pde6a Gucy2d Cdhr1 Opn1sw Myrip Pdc Impg1 Cnga1 Prcd Gngt1 Ush2a Ttll4 Ift122 Iqcb1 Mak | 6 |
| GOBP PHOTOTRANSDUCTION | 4.79E-10 | 3.35E-07 | 8.01E-01 | -0.766 | -2.363 | 45 | Grk1 Cngb1 Gnat2 Cabp4 Pde6g Gngt2 Guca1b Sag Guca1a Pde6a Gucy2d Unc119 Opn1sw Pde6c Aipl1 Pdc Cnga1 Gngt1 Asic2 Rs1 Cds1 Gpr88 Pitpnm1 | 7 |
| GOBP CIRCADIAN RHYTHM | 8.43E-10 | 5.16E-07 | 8.01E-01 | -0.53 | -2.03 | 188 | Nr1d2 Sik1 Id1 Per3 Ciart Nr1d1 Dbp Klf10 Tnfrsf11a Aanat Usp2 Fbxl17 Bhlhe41 Tph2 Drd4 Ddc Nr2f6 Tph1 Rpe65 Klf9 Kdm5a Id4 Id3 Adora2a Cavin3 Per1 Kmt2a Nr1h3 Ass1 Siah2 Hdac3 Kdm2a Ube3a Adrb1 Zfhx3 Bhlhe40 Sirt1 Sin3a Mycbp2 Rorb Gpr157 Tyms Srd5a1 Noct Per2 Hs3st2 Ppargc1a Prokr2 Egr1 Kdm8 Pspc1 Setx Cdk1 Crem | 8 |
| GOCC NON MOTILE CILIUM | 1.67E-09 | 9.09E-07 | 7.88E-01 | -0.584 | -2.145 | 133 | Ppef2 Grk1 Cacna1f Cngb1 Rpgrip1 Gnat2 Arr3 Rd3 Pde6g Rp1l1 Rom1 Ttc8 Guca1b Mertk Tulp1 Sag Crb1 Guca1a Pde6a Gucy2d Cdhr1 Ahi1 Opn1sw Myrip Drd1 Pdc Impg1 Arl13b Cnga1 Prcd Gngt1 Ush2a Ttll4 Ift122 Iqcb1 Glis2 Mak Cdc14a Vcan | 9 |
| GOBP DETECTION OF ABIOTIC STIMULUS | 2.45E-09 | 1.20E-06 | 7.75E-01 | -0.601 | -2.16 | 114 | Grk1 Cacna2d4 Lxn Cacna1f Cngb1 Gnat2 Cabp4 Pde6g Gngt2 Rom1 Guca1b Nr2f6 Tulp1 Sag Crb1 Guca1a Pde6a Gucy2d Unc119 Rpe65 Opn1sw Pde6c Ntsr1 Aipl1 Pdc Cnga1 Reep6 Gngt1 Adgrv1 Arrb2 Asic2 Rs1 Cds1 Ano1 Gpr88 | 10 |
| GOCC EXTERNAL ENCAPSULATING STRUCTURE | 5.79E-09 | 2.53E-06 | 7.62E-01 | -0.431 | -1.782 | 424 | Tinagl1 Postn Col26a1 Impg2 Bmper Pdgfb Mxra7 Ccn1 Col20a1 Col4a4 Dpt Adamts1 Sspo Nyx Srpx Angptl2 Col14a1 Tgfbi Lum Npnt Tnn Optc Rbp3 Eln Pkhd1l1 Lgals3 Mgp Timp3 Lamb1 Col4a3 Col2a1 Crispld2 Tnfrsf11b Ccbe1 Lgals1 Impg1 Lama5 Wnt3 Lama1 Cilp Prelp Ecm1 Thbs4 Cbln1 Ccn2 Ush2a Ltbp1 Igfbp7 Anxa11 Hmcn1 Tgfbr3 Col6a6 Fbn2 Flrt1 Fbln5 Serping1 Ccn4 Egflam Dcn Col4a1 Fgf1 Vasn Kazald1 Sbspon Fn1 Lrrc17 Efemp2 Gdf10 Sod3 Vcan Pcsk6 Angpt2 Abi3bp Mfge8 Mmp9 Fmod Pf4 Myoc Egfl7 Wnt2b Fras1 Dmp1 Col4a6 Lingo1 Rarres2 Lama2 Lamc1 Lrig2 Ntn4 Gpc4 Ntng2 P3h1 Col6a3 Olfml2a Col1a1 Matn4 Clec3b Lrrc32 Adamts17 Chi3l1 Dlg1 Mfap5 Serpinf1 Adamts12 Col4a5 Ctsc Cstb Frem2 Ccdc80 Col5a1 Oc90 Vtn Ltbp3 Adamts18 Clu Nid1 Col3a1 Cdon Dst Anxa4 Icam1 RGD1565033 Plscr1 Lingo3 Ssc5d Col24a1 Tgfb1i1 Lgals3bp Prg2 Glg1 Vwa2 Fgl2 Pcolce Mdk Bmp7 Col9a1 Wnt11 Adamts7 Spock2 Col7a1 Mmp19 Vwf Col22a1 Anxa1 Bgn Mmp23 Col9a3 Smoc1 Lama4 Fbn1 Col12a1 Sulf1 Ptn Timp4 Aebp1 Rtbdn Adamtsl3 Adipoq Sfrp1 Shh Col17a1 | 11 |
| GOMF EXTRACELLULAR MATRIX STRUCTURAL CONSTITUENT | 6.20E-09 | 2.53E-06 | 7.62E-01 | -0.583 | -2.132 | 129 | Tinagl1 Postn Impg2 Ccn1 Col4a4 Dpt Srpx Col14a1 Tgfbi Lum Npnt Optc Eln Umodl1 Mgp Lamb1 Col4a3 Col2a1 Impg1 Lama5 Lama1 Cilp Prelp Ecm1 Ltbp1 Igfbp7 Hmcn1 Col6a6 Fbn2 Fbln5 Dcn Col4a1 Sbspon Fn1 Efemp2 Vcan Abi3bp Mfge8 Fmod Fras1 Col4a6 Lama2 Lamc1 Col6a3 Col1a1 Chi3l1 Mfap5 Col4a5 Col5a1 Vtn Nid1 Col3a1 Col24a1 Prg2 Fgl2 Pcolce Col9a1 Col7a1 Vwf Col22a1 Bgn Col9a3 Lama4 Fbn1 Col12a1 Aebp1 Adipoq Col17a1 | 12 |
| GOBP RESPONSE TO RADIATION | 6.76E-09 | 2.55E-06 | 7.62E-01 | -0.438 | -1.797 | 395 | Sik1 Per3 Grk1 Cacna2d4 Cacna1f Cngb1 Gnat2 Cabp4 Tnfrsf11a Aanat Usp2 Ctns Fbxl17 Pde6g Slc24a1 Fos Gngt2 Trpm1 Dusp1 Rom1 Atm Guca1b Nr2f6 Tulp1 Sag Crb1 Guca1a Pde6a Gucy2d Unc119 Rpe65 Cpt1b Eya3 Opn1sw Pde6c Map3k4 Aipl1 Drd1 Pdc Cdc25a Aurkb Cnga1 Per1 Reep6 Ercc2 Gngt1 Babam1 Nfatc4 Dnmt3a Nedd4 Hif1a Bhlhe40 Mrnip Asic2 Rs1 Sirt1 Hyal1 Atr Cds1 Blm Eif2ak4 Gpr88 Xpc Angpt2 Primpol Pias1 Bak1 Pitpnm1 Crip1 Mmp9 Polk Per2 Tspyl5 Net1 Egr1 Rev1 Rad54l | 13 |
| GOMF UNFOLDED PROTEIN BINDING | 9.30E-09 | 3.14E-06 | 7.48E-01 | 0.583 | 2.227 | 101 | Cct3 Hyou1 Ppib Ptges3 Spg7 Dnajb11 Calr Hsp90b1 Hspa5 Cct5 Cct6a Ruvbl2 Cryab Dnaja4 Tubb4b Hsp90ab1 Hspe1 Grpel1 Clgn Dnaja1 Srsf10 Pfdn1 Tcp1 Cct7 Pfdn4 Nudc Canx Hspd1 Syvn1 Cct4 Dnajb13 Dnajb2 Mkks Dnaja2 Dnajb6 Pet100 Uggt1 Tapbp Cdc37 | 14 |
| GOBP PROTEIN FOLDING | 9.62E-09 | 3.14E-06 | 7.48E-01 | 0.479 | 2.012 | 200 | Pdia6 Cct3 Sdf2l1 Ppib Ptges3 Pdia4 Dnajb11 Ahsa1 Calr Entpd5 Hsph1 Hsp90b1 Hspa5 Cct5 Cct6a Grn Ruvbl2 Cryab Dnaja4 Sdf2 Hsp90ab1 Pdia3 Ppil1 Gng2 Gnao1 Fkbp1a Hspe1 Grpel1 Clgn Dnaja1 Amfr Pfdn1 Tcp1 Vcp Cct7 Pfdn4 Dnajc5 Chordc1 Pex19 Erp44 Nudc Canx Hspd1 Cct4 Dnajb13 Bag1 Dnajb2 Mkks Dnajc3 Erp29 Dnaja2 Fut10 Dnajb6 Rad23b Hspbp1 | 15 |
| GOCC COLLAGEN CONTAINING EXTRACELLULAR MATRIX | 2.19E-08 | 6.71E-06 | 7.34E-01 | -0.456 | -1.838 | 319 | Tinagl1 Postn Col26a1 Impg2 Pdgfb Mxra7 Ccn1 Col20a1 Col4a4 Dpt Adamts1 Srpx Angptl2 Col14a1 Tgfbi Lum Npnt Tnn Rbp3 Eln Lgals3 Mgp Timp3 Lamb1 Col4a3 Col2a1 Lgals1 Impg1 Lama5 Lama1 Cilp Prelp Ecm1 Thbs4 Cbln1 Ccn2 Ush2a Ltbp1 Igfbp7 Anxa11 Hmcn1 Col6a6 Fbn2 Fbln5 Serping1 Egflam Dcn Col4a1 Kazald1 Sbspon Fn1 Efemp2 Gdf10 Sod3 Vcan Pcsk6 Angpt2 Abi3bp Mfge8 Mmp9 Fmod Pf4 Myoc Egfl7 Wnt2b Fras1 Col4a6 Rarres2 Lama2 Lamc1 Ntn4 Gpc4 Ntng2 P3h1 Col6a3 Col1a1 Matn4 Clec3b Dlg1 Mfap5 Serpinf1 Col4a5 Ctsc Cstb Frem2 Ccdc80 Col5a1 Vtn Ltbp3 Clu Nid1 Col3a1 Cdon Dst Anxa4 Icam1 RGD1565033 Plscr1 Ssc5d Col24a1 Tgfb1i1 Lgals3bp Prg2 Vwa2 Fgl2 Pcolce Mdk Bmp7 Col9a1 Col7a1 Vwf Anxa1 Bgn Mmp23 Col9a3 Smoc1 Lama4 Fbn1 Col12a1 Sulf1 Ptn Aebp1 Rtbdn Adipoq Sfrp1 Shh Col17a1 | 16 |
| GOBP RHYTHMIC PROCESS | 2.43E-08 | 7.01E-06 | 7.34E-01 | -0.479 | -1.898 | 261 | Nr1d2 Sik1 Id1 Per3 Ciart Nr1d1 Dbp Fzd4 Klf10 Tnfrsf11a Aanat Usp2 Fbxl17 Bhlhe41 Tph2 Drd4 Ddc Adamts1 Nr2f6 Tph1 Rpe65 Klf9 Kdm5a Id4 Id3 Adora2a Cavin3 Sgpl1 Per1 Kmt2a Nr1h3 Hlf Ass1 Siah2 Hdac3 Kdm2a Ube3a Adrb1 Ets1 Zfhx3 Bhlhe40 Arrb2 Sirt1 Sin3a Mycbp2 Pam Rorb Has2 Gpr157 Gpr149 Tyms Srd5a1 Noct Per2 Hs3st2 Ppargc1a Prokr2 Egr1 Kdm8 Pspc1 Setx Cdk1 Dtl Casp3 Crem | 17 |
| GOCC ENDOPLASMIC RETICULUM PROTEIN CONTAINING COMPLEX | 9.94E-08 | 2.71E-05 | 7.05E-01 | 0.532 | 2.08 | 118 | Pdia6 Hyou1 Sdf2l1 Ormdl3 Calr Emc9 Emc3 Srebf2 Sel1l Hspa5 Srp9 Derl3 Pdia3 Ostc Ormdl2 Ddost Fkbp1a Sec11a Amfr Pigs Vcp Stt3b Sec61g Syvn1 Hsd17b12 Pyurf Emc8 Dad1 Rnf125 RT1-CE10 Sec63 Tapbp Selenos Zw10 Aup1 Arl6ip1 Prkcsh Gpaa1 Rpn1 | 18 |
| GOBP CIRCADIAN REGULATION OF GENE EXPRESSION | 6.87E-07 | 1.77E-04 | 6.59E-01 | -0.642 | -2.1 | 62 | Id1 Per3 Ciart Nr1d1 Usp2 Bhlhe41 Kdm5a Id4 Id3 Cavin3 Per1 Kmt2a Hdac3 Kdm2a Zfhx3 Bhlhe40 Sirt1 Mycbp2 Noct Per2 Ppargc1a Egr1 Kdm8 | 19 |
| GOCC CHAPERONE COMPLEX | 1.26E-06 | 3.04E-04 | 6.44E-01 | 0.77 | 2.254 | 26 | Stip1 Cct3 Sdf2l1 Ptges3 Dnajb11 Cct5 Cct6a Sdf2 Hsp90ab1 Tcp1 Cct7 Cct4 Cdc37 Psmg1 | 20 |
| GOBP DETECTION OF LIGHT STIMULUS INVOLVED IN SENSORY PERCEPTION | 1.30E-06 | 3.04E-04 | 6.44E-01 | -0.856 | -2.143 | 17 | Cacna2d4 Cacna1f Cngb1 Gnat2 Rom1 Tulp1 Crb1 Rpe65 Reep6 | 21 |
| GOBP RHODOPSIN MEDIATED SIGNALING PATHWAY | 3.61E-06 | 8.04E-04 | 6.27E-01 | -0.78 | -2.115 | 24 | Grk1 Cngb1 Pde6g Guca1b Sag Guca1a Pde6a Gucy2d Aipl1 Cnga1 Gngt1 Rs1 | 22 |
| GOBP MITOCHONDRIAL TRANSLATION | 4.21E-06 | 8.97E-04 | 6.11E-01 | 0.484 | 1.919 | 129 | Mrpl21 Mrps10 Mrps6 Taco1 mrpl11 Rars2 Mrps18c Mrps11 Mrps18b Mrps35 Trub2 Mrps22 Uqcc1 Mrpl45 Mrpl13 Mrpl49 Mrps30 Mrps27 Gatb Gfm1 Mrpl23 Mrps33 Mrpl15 Mtif3 Mrps18a Aars2 Gadd45gip1 Mrps26 Mrps16 Malsu1 Mrps17 Mrpl48 Mrpl17 Mrpl40 Hars1 Mrpl16 Wars2 Mpv17l2 C1qbp Rpusd3 Mrps7 Lars2 Mrps36 mrpl24 Mrpl20 Mrpl4 Iars2 Chchd1 Qrsl1 Ears2 Mrpl42 | 23 |
| GOBP PHOTOTRANSDUCTION VISIBLE LIGHT | 4.57E-06 | 9.06E-04 | 6.11E-01 | -0.749 | -2.116 | 29 | Grk1 Cngb1 Pde6g Guca1b Sag Guca1a Pde6a Gucy2d Pde6c Aipl1 Cnga1 Gngt1 Rs1 | 24 |
| GOBP EXTERNAL ENCAPSULATING STRUCTURE ORGANIZATION | 4.62E-06 | 9.06E-04 | 6.11E-01 | -0.413 | -1.668 | 329 | Postn Flot1 Pdgfb Ccn1 Nphs1 Col4a4 Tie1 Dpt Adamts1 Col14a1 Tgfbi Lum Npnt Optc Eln Lamb1 Col4a3 Col2a1 Qsox1 Crispld2 Lcp1 Tnfrsf11b Sh3pxd2a Cav2 Lama5 Lama1 Tgm3 Reck Ercc2 Ccn2 Foxf1 Atxn1l Fbn2 Itga10 Ets1 Fbln5 Itga5 Egflam Loxl4 Dcn Col4a1 Kazald1 Has2 Fn1 Efemp2 Vcan P3h4 Dpp4 Sh3pxd2b B4galt1 Mmp9 Fmod Atp7a Dmp1 Col4a6 Lama2 Lamc1 Klk4 Spint2 Ntn4 Ntng2 Ibsp Col6a3 Olfml2a Bcl3 Col1a1 Matn4 Notch1 Ctsk Adamts17 Pdpn Itga11 Mfap5 Adamts12 Col4a5 Itgb2 Ccdc80 Col5a1 Gas6 Vtn Myh11 Ltbp3 Adamts18 Nid1 Col3a1 Scube3 Itga2 Icam1 Eng Itga1 Cd44 Fap Col24a1 Foxc1 Prdx4 Ext1 Foxc2 Col9a1 Adamts7 Itgb5 Nf1 Spock2 Col7a1 Itga8 Mmp19 Itgav Vwf Colgalt1 Col22a1 Fgfr4 Bgn Tll2 Mmp23 Col9a3 Smoc1 Lama4 Fbn1 Abl1 Itga3 Col12a1 Sulf1 Pdgfa Aebp1 Adamtsl3 Tmem38b Col17a1 | 25 |
