## Supplemental S1-S4 for "Comparative gene pathway analysis during adolescent binge-EtOH exposure, withdrawal, and following abstinence": Table_S3.docx

**Table S3A. Top 25 Pathways from GSEA (based on p-value) using Gene Sets of MSigDB ImmuneSigDB for PND 46**

| Pathway | pval | padj | log2err | ES | NES | size | Genes | Rank |
| --- | --- | --- | --- | --- | --- | --- | --- | --- |
| GSE9650 NAIVE VS EFF CD8 TCELL DN | 1.35E-20 | 6.59E-17 | 1.18E+00 | -0.643 | -2.552 | 177 | Lgals1 Plscr1 Fgl2 Lgals3 Anxa2 Cdk1 Anxa6 Txndc5 Gem Atp6v0e1 Ppib Itgal S100a10 Elovl1 Stab1 Anxa1 Litaf S100a4 Lamc1 Kcnj8 Tmem37 Ahnak Kif22 Cd48 Dkk2 Casp7 Ctla4 Fcgrt Asl Casp1 Myo1f Arhgdib Chd7 Padi2 Bcl2a1 Tspan31 Ctsa Hopx Ptpn13 Dock5 Ptgr1 St3gal4 Unc119 RT1-A2 Dap Itgax Mt2A Carhsp1 Casp4 Lrp10 Cd68 Il18rap Prelid1 S100a6 Adam19 S100a11 P3h4 Itgam Dbi Prr13 Slc66a3 Itgb1 Cstb Lamtor5 Entpd1 Ccr5 Spdl1 Mapk3 Xdh Wnt3 Tspo S100a13 Sirt2 | 1 |
| GOLDRATH NAIVE VS EFF CD8 TCELL DN | 6.10E-15 | 1.49E-11 | 9.97E-01 | -0.603 | -2.39 | 175 | Lgals1 Plscr1 Fgl2 Lgals3 Anxa2 Kif11 Cdk1 Ect2 Vcl Gem Rom1 S100a10 Plp2 Anxa1 F2rl3 Litaf Ifi30 S100a4 Emp1 Tmem37 Ube2t Mki67 Kif22 Cd48 Ifitm3 Dstnl1 Cd99 Prc1 Serpinb9 Cdca8 Emp3 Myo1f Scpep1 Tnfrsf9 Nudt4 Nusap1 Bcl2a1 Serpinb6b Hopx Prdm1 Dock5 Ezh2 Tk1 Unc119 Plin2 Tmem14c Itgax Odc1 Mt2A Carhsp1 S100a6 Rfc5 Cks2 H1f2 Txn1 Cd244 Ccnf Itgb1 Kif4a Cmc2 Top2a Ccr5 Kifc1 Pcna C3 Mcm5 Bmp2k Brca1 Ncaph Pclaf Myadm | 2 |
| KAECH NAIVE VS DAY8 EFF CD8 TCELL DN | 5.27E-14 | 8.56E-11 | 9.65E-01 | -0.578 | -2.294 | 177 | Lgals1 Plscr1 Lgals3 Anxa2 Kif11 Cdk1 Anxa6 Bak1 Txndc5 Gem Atp6v0e1 Rom1 S100a10 Plp2 Anxa1 F2rl3 Litaf S100a4 Lamc1 Emp1 Ahnak Mki67 Kif22 Cd48 Dstnl1 Casp7 Prc1 Ctla4 Srp14 Serpinb9 Casp1 Myo1f Nusap1 Chd7 Bcl2a1 Tspan31 Hopx Ptpn13 Prdm1 Rrbp1 Dock5 Ptgr1 RT1-A2 Itgax Rrm2 Carhsp1 Casp4 Il18rap Prelid1 S100a6 Adam19 S100a11 Arl6 Dbi Prr13 Slc66a3 Txn1 Cd44 Itgb1 Prdx1 Top2a Lamtor5 Ccr5 Clic4 Septin10 S100a13 Ndufb9 Ncaph Ap2s1 Cdc34 Tnfrsf1b Fam89b | 3 |
| GSE24634 TREG VS TCONV POST DAY10 IL4 CONVERSION DN | 2.94E-12 | 3.58E-09 | 8.99E-01 | -0.563 | -2.236 | 178 | Anpep Fgl2 Plekhb1 Nr1h3 Grn Pmp22 Cyp1b1 Fkbp9 Mrc1 Gsn Tubb6 Clec10a Lmna Plbd1 Ifi30 Abcb1b Lamc1 Myof RT1-Da Slc29a3 Stom Hmox1 Gla Cd63 Eng Rasgrp3 Vat1 Fcgrt Itgb2 St14 Scpep1 Cd14 Cd163 Sparc Lpar6 Ctsa Tcn2 Ahcy Ldlrap1 Ctsc Ftl1 Qprt Gpx1 Slc31a2 Npc2 Ctsl Itgax Lipa Slc38a6 S100a9 Cyp27a1 Chn2 C1qa Tfpi Itgam Ccr1 Echdc3 Acsl1 Smarcd3 Idh1 Slc2a6 Man2b1 Taf4 Steap3 Galnt12 C2 Clec7a Naga Ctsz Pygl Lhfpl2 Cd86 | 4 |
| KAECH NAIVE VS DAY15 EFF CD8 TCELL DN | 4.37E-11 | 3.82E-08 | 8.51E-01 | -0.554 | -2.195 | 175 | Lgals1 Plscr1 Fgl2 Lgals3 Anxa2 Txndc5 Itgal S100a10 Anxa1 Litaf S100a4 Lamc1 Raly Kcnj8 Emp1 Srgn Ahnak Cd4 Dstnl1 Il13ra2 Gmfg Casp7 Ctla4 Fcgrt Serpinb9 Emp3 Casp1 Myo1f Ccnd3 Bcl2a1 Tspan31 Hopx Ptpn13 Prdm1 Rrbp1 Dock5 RT1-A2 Cr1l Itgax Reck Man2a1 Carhsp1 Casp4 Lrp10 Il18rap S100a6 Adora2a Nbeal2 S100a11 Prr13 Sspn H1f2 Txn1 Cd44 Enpp1 Itgb1 Prdx1 Cmc2 Entpd1 Ccr5 St3gal6 Xdh Laptm5 S100a13 Alcam | 5 |
| GSE24634 TEFF VS TCONV DAY10 IN CULTURE DN | 4.71E-11 | 3.82E-08 | 8.51E-01 | -0.544 | -2.165 | 181 | Fgl2 Lgals3 Plekhb1 Stard8 Nr1h3 Vamp3 Tlr8 Ctsh Grn Vcl Lyz2 Mrc1 Itga5 Clec10a Lmna Vtn Gabarap Plbd1 Ifi30 Sgpl1 RT1-Da Cd63 Dab2 Itgb2 Hexa Tcf7 Scpep1 Cavin1 Ldlrap1 Hmg20b Ftl1 Milr1 RT1-A2 Slc2a9 Npc2 Itgax Ncf2 Lipa S100a9 Slc8b1 Il6r Sav1 Tlr2 C1qa P2ry6 Itgam Ehd4 F11r Cnnm4 Ccr1 Echdc3 Clic4 Idh1 Slc2a6 C3 Dysf Man2b1 Tspo S100a13 | 6 |
| KAECH DAY8 EFF VS DAY15 EFF CD8 TCELL UP | 1.55E-10 | 1.08E-07 | 8.27E-01 | -0.531 | -2.125 | 187 | Tagln2 Lgals1 Lgals3 Anxa2 Kif11 Cdk1 Anxa6 Gyg1 Ppib Rom1 Plp2 F2rl3 Selenoh Cpt1a Mki67 Kif22 Cd48 Casp7 Ssna1 Prc1 Srp14 Ldha Cib1 Arhgdib Nusap1 Psmb6 Txndc17 Ptgr1 Dap Ctnna1 Nfe2 Rrm2 Sem1 Usp18 Prelid1 Arl6 Dbi Slc66a3 Txn1 Tubb5 Kif4a Prdx1 Cmc2 Rpsa Lmnb1 Top2a Lamtor5 Spdl1 Hm13 Mcm5 Xdh Ndufb9 Ncaph Rpn1 Cox6c Pclaf Cdc34 Iqgap3 Ccna1 Nelfcd Fam89b Ints9 Lgals9 MGC95208 | 7 |
| GSE2405 HEAT KILLED LYSATE VS LIVE A PHAGOCYTOPHILUM STIM NEUTROPHIL 24H UP | 4.31E-10 | 2.62E-07 | 8.14E-01 | -0.525 | -2.091 | 182 | Angptl2 Lgals3 Anxa2 P2rx4 Hsd3b7 Grn Vcl Pmp22 Pard6g Mgp Tbata Hspa1a Anxa1 Litaf Ifi30 S100a4 Srgn Actg2 Ahnak Ifitm3 Fn1 Gnb3 Ly86 Tagln Myh11 Pkp1 Pltp Rgs5 Tpm2 Serpinb9 Col4a2 St14 Slco2a1 Decr1 Bcl2a1 Tec RT1-Ba Gpx1 Plaat3 Sfrp1 Hmgn3 Plin2 Csrp2 Ncf2 Ctnna1 Copz2 Casp4 Rnh1 Cd68 Usp18 S100a6 Cxcl9 Wfdc2 Prr13 Ehd4 Cstb Axl Arhgef16 Hexb Tmem176a Mfge8 C3 Lrrk1 Slc6a8 Wfikkn2 Alcam Mmp23 Serpinb1a Dbndd2 Plek C2 Inf2 Clec7a Syngr2 Lpcat2 Swap70 | 8 |
| GSE24142 DN2 VS DN3 THYMOCYTE UP | 6.62E-10 | 3.58E-07 | 8.01E-01 | -0.516 | -2.054 | 180 | Tagln2 Lgals1 Plscr1 P2rx4 Egfl7 Mrc2 Niban1 Clec10a Anxa1 Litaf Ldaf1 Hhex Vim S100a4 Srgn Cd48 Ripk1 Flna Tal1 Gm2a Atp1b3 Itga2b Deptor Esm1 Tas1r1 Optc Eci1 Cavin1 Mgst1 Slc25a24 Spi1 Ak2 Cnn2 Cables1 Ppic Csrp2 Ca13 Itgax Mif4gd Nfe2 Otulinl Rnh1 S100a6 Rgs18 H1f6 Nudt7 Ctnnal1 Cd44 Cd244 Fads3 Gcnt1 Uap1l1 Dhrs11 Lrrk1 Serpinb1a Steap3 Fanca Myadm Kctd14 Tspan2 Sh2d3c Pygl Ccdc34 Parp16 | 9 |
| GSE22140 HEALTHY VS ARTHRITIC GERMFREE MOUSE CD4 TCELL DN | 1.49E-09 | 6.48E-07 | 7.88E-01 | -0.522 | -2.077 | 179 | Lgals1 Nid1 Tgfbi P2rx4 Nqo1 Pmp22 Igf1 Gem Folr2 Mrc1 Tex30 Ctsk Lmna Hspa1a Stab1 P2rx7 Oasl Sgta S100a4 Emp1 Timp1 Lgmn Hmox1 Il1b Cd99 Slc3a2 Tnfrsf11a Phldb1 Clic1 Dnajc8 Pltp Ifi44 F13a1 Btg2 Emp3 Slc7a11 Tnfrsf9 Cd163 Cpq Ifit2 Ctsl Ifit3 Tlr2 Rgs16 Pmaip1 Adam19 Zfhx3 Crem Ccr1 Gpr107 Furin Tgfa Stx4 Serpine1 Ppp1r15a Rpn1 Irf1 Gna15 Atp6v0d1 Tnfrsf1b Selenop Ctsz Npc1 Nfkb1 Eif4a3 Pdgfb | 10 |
| KAECH NAIVE VS MEMORY CD8 TCELL DN | 1.58E-09 | 6.48E-07 | 7.88E-01 | -0.523 | -2.07 | 170 | Lgals1 Klf4 Plscr1 Fgl2 Anxa2 Txndc5 S100a10 Anxa1 S100a4 Kcnj8 Emp1 Tmem37 Ahnak Dpp7 Slco3a1 Ctla4 Fcgrt Itgb2 Adgre5 Casp1 Myo1f Ccnd3 Cyb5r3 Bcl2a1 Tspan31 Hopx Ptpn13 Dock5 St3gal4 Unc119 RT1-A2 Bche Itgax Odc1 Xrcc5 Ctnna1 Reck Casp4 Tob1 Cdk4 Il18rap S100a6 Adam19 Hcfc1r1 S100a11 Arl6 Prr13 Txn1 Tnfsf10 Cd44 Itgb1 Jund Ccr5 Pcna St3gal6 Xdh Tent5c S100a13 Cd7 Septin1 Pclaf Cdc34 Tnfrsf1b | 11 |
| GSE7509 DC VS MONOCYTE UP | 1.60E-09 | 6.48E-07 | 7.88E-01 | -0.517 | -2.068 | 189 | Angptl2 Atf3 Lgals1 Lgals3 Col4a1 Ctsh Kdelr1 Cyp1b1 Igf1 Folr2 Gyg1 Crtapl1 Gsn Anxa1 Plbd1 Litaf Hsd17b11 Abcb1b Polr2a Actg2 Cd63 Gm2a Csf2rb Ly86 Timp3 Dab2 Csf1 Hexa Pltp Nes Tec Mgst1 Gstz1 Cnn2 Antxr2 Dcn Dap Anxa5 Plin2 Itgax Phldb2 Rab34 C1qc Tmem109 Cd68 C1qa Glb1 H1f2 Thbd Entpd1 Ccr5 Mfge8 Stx4 Ndufa3 Tm2d2 Pithd1 Cnih1 Wbp1l Zdhhc14 | 12 |
| GOLDRATH NAIVE VS MEMORY CD8 TCELL DN | 2.09E-09 | 7.83E-07 | 7.75E-01 | -0.524 | -2.075 | 174 | Lgals1 Plscr1 Fgl2 Lgals3 Anxa2 Txndc5 S100a10 Plp2 Anxa1 Plbd1 Litaf Ldaf1 Hsd17b11 Abcb1b S100a4 Kcnj8 Emp1 Ostf1 Plat Tmem37 Ahnak Notch4 Dstnl1 Fcgrt Serpinb9 Casp1 Myo1f Eci1 Ccnd3 Bcl2a1 Serpinb6b Ptpn13 Prdm1 Cybb Dock5 Antxr2 St3gal4 Tk1 Abhd5 Itgax Odc1 Ctnna1 Reck Il10rb Golim4 Casp4 Il18rap S100a6 Hcfc1r1 Rasa4 Nbeal2 Arl6 Slc66a3 Cd44 Acy1 Enpp1 Itgb1 Ccr5 Cpne3 Cd22 St3gal6 Pbx3 S100a13 Mapk12 Dbndd2 Il15 Myadm | 13 |
| GSE25123 IL4 VS IL4 AND ROSIGLITAZONE STIM MACROPHAGE DAY10 UP | 2.74E-09 | 9.54E-07 | 7.75E-01 | -0.512 | -2.033 | 177 | Angptl2 Lgals1 Plscr1 Lgals3 Id3 Il1r1 Ppfibp1 Slc39a4 Rom1 Asb2 Litaf Crybg3 Hhex Srgn Mki67 Plekhh3 Lgmn PCOLCE2 Nid2 Cd74 Il1rl1 Penk St14 Casp1 Myo1f Ehbp1l1 Tnfrsf9 Rilpl2 Tmbim1 Bcl2a1 Snx33 Vps54 Slc25a24 Ptpn13 Tjp2 Tnfrsf4 Unc119 Poglut3 Plekhg3 Npr1 Il10rb S100a6 Vdr S100a11 Cass4 Snai2 Ehd4 Kif4a Gcnt1 Top2a Spdl1 Gpr20 Rhoc Ifitm2 Brca1 Gdpd5 Serpinb1a Mapk12 Kdelr2 Cd38 Gna15 | 14 |
| GSE1432 CTRL VS IFNG 24H MICROGLIA DN | 3.12E-09 | 1.01E-06 | 7.75E-01 | -0.518 | -2.046 | 169 | Klf4 Fgl2 Gch1 Serping1 RT1-Db1 Sp110 Apobec3 Myof Rhbdf2 Ifitm3 Stom Arid5a Adgre5 Apol3 Ifi44 Tgm2 Gbp2 Cxcl10 Ces1d Tapbpl RT1-Bb Ak2 Ezh2 Fcgr1a Phf11 Vamp5 RT1-A2 Psmb8 Psmb9 Myd88 Isoc1 Lap3 Rhobtb3 Mvp Mt2A Casp4 Ifit3 Tmem109 Rnf114 Cxcl9 Ciita Sspn H1f6 Tnfsf10 Cstf3 Gimap6 Scarf1 Psme4 Rsad2 Lmnb1 Nhej1 Tap1 Psme1 Dynlt1 Ifitm2 C1rl Top1 Il15 Cd38 Irf1 Plek Mob1a Ascl2 P2ry14 Vrk2 | 15 |
| GSE6269 FLU VS STAPH AUREUS INF PBMC DN | 6.98E-09 | 2.13E-06 | 7.62E-01 | -0.521 | -2.033 | 153 | Anpep Lgals3 Vamp3 P2rx4 Grn Cybrd1 Lama4 Lyz2 Rhoa Fhl3 S100a10 Plp2 Stab1 Hs3st3b1 Plbd1 Ifi30 Vim Srgn Timp1 Cd63 Slco3a1 Zyx Wls Cd93 Laptm4a Spi1 Adipor1 Fut4 Phc2 Fcgr1a Npc2 S100a9 Tlr2 Flvcr2 Dse Mxd1 Socs3 Acsl1 Smarcd3 Dysf Serpinb1a Impa2 P2ry2 Ppp1r15a Steap3 Gna15 Plek C2 Plin3 Clec7a Arf4 Asgr2 Vcan | 16 |
| GSE19888 ADENOSINE A3R ACT VS TCELL MEMBRANES ACT IN MAST CELL UP | 1.55E-08 | 4.20E-06 | 7.34E-01 | -0.502 | -1.997 | 179 | Nid1 Pcolce Serping1 Bak1 Rarres2 Lama4 Mrc2 Hip1 Lum Casp12 F2rl3 Sned1 Col3a1 Fermt3 Timp1 Prtg Galns Pkhd1l1 Slc38a10 Tgfb1 C4b Adgre5 Agtrap Gbp2 Casp1 Myo1f Arhgdib Oaf Ctsc Tln1 Phc2 Ppic Dcn Plin2 Gpx7 B2m Psmb9 Itga1 C1qc Ssc5d Pid1 Arhgap30 Cxcl9 P2ry6 Eif1b Ppp1r18 Itgam Eif4e3 Slc16a12 Sdhaf2 Col6a1 Dynlt1 Cpxm1 Irf1 Fanca Gnai2 Kiss1r Slc25a45 Gpsm3 Gpx3 Slc13a5 Ecscr Sqor Ctsz Slc12a7 Mgat1 Ptpn18 Ifnar2 C1r Adamts2 Col5a2 Tbxa2r Smpdl3b Crip1 | 17 |
| GSE21379 WT VS SAP KO TFH CD4 TCELL UP | 1.55E-08 | 4.20E-06 | 7.34E-01 | -0.502 | -1.997 | 179 | Aplnr Rnd3 Tie1 Col4a1 Fah Frk Abcc3 Gpr4 Fkbp9 Mrc1 Bgn Guca1a Vamp8 Sh3rf2 Arhgap29 Adgra2 Dab2 Shroom4 Cdh5 Col4a2 Tmem204 Cd14 Rhoj Lamb2 Dock5 Tle6 Esam Mtus1 Lrg1 Tlcd3a Dock6 Exoc3l1 Rab34 Serpinh1 Cyp27a1 Sem1 Trip6 Tmem170a Kdr Tfpi Tead2 Scarf1 Selenom Card10 Axl Sox18 Bok Myom1 C3 Xdh Rnd2 Bmp2k Atp5f1e Maf Ctnnd1 Cd38 Fhl1 Zfp385a Ap2s1 Spa17 Evc2 Ecscr Dlk1 Nek5 Gpx8 Ajuba C1r Rab3il1 Dmkn Smpdl3b | 18 |
| GSE13485 CTRL VS DAY7 YF17D VACCINE PBMC DN | 1.74E-08 | 4.46E-06 | 7.34E-01 | -0.505 | -1.996 | 170 | Plscr1 Nid1 Gch1 Gngt2 Serping1 Heg1 Sp110 Tpx2 Rnf213 Oasl Myof Mki67 Ifitm3 Prpsap1 Atp10a Cnp RGD1310951 Sp100 Ifi44 Odf3b Cxcl10 Parp14 Tcn2 Lamp3 Parp9 Fcgr1a Timeless Slc31a2 Phf11 Ifit2 Psmb8 Adap2 Psmb9 Myd88 Lap3 Mt2A Casp4 Ifit3 Trip6 Mvb12a Usp18 Tnfsf10 Utrn Rsad2 Nexn Ccr1 Trim69 Ccr5 Axl Tap1 Ankfy1 Ifitm2 Mthfd2 Fbxo6 Trim5 Il15 Cd38 Shisa5 Kctd14 C2 | 19 |
| GSE22886 NAIVE CD4 TCELL VS DC DN | 1.89E-08 | 4.60E-06 | 7.34E-01 | -0.49 | -1.97 | 195 | Fgl2 Anxa2 Vamp3 Tgfbi RT1-Db1 Ctsh Grn Atp6v0e1 Gsn Gstp1 Hspa1a Gabarap Ifi30 Vamp8 Vim RT1-Da Gla Cd63 Plod1 Cd74 Fcgrt Bckdk Cox8a Slc1a5 Bet1 St14 Tuba1b Ctsc Ftl1 Snap29 Gpx1 Dera Slc31a2 Anxa5 Itgax Src Ncf2 Lipa Isoc1 Ctnna1 Lap3 Polr3k Nfe2l2 Rnh1 Serf2 Flvcr2 S100a11 Glb1 Prdx1 Cstb Ccr1 Akip1 Clic4 Idh1 Gng5 Tax1bp3 Man2b1 Mthfd2 Mreg Tm9sf1 Plcg2 Ap2s1 Clec7a Syngr2 Prkcd | 20 |
| GSE22140 GERMFREE VS SPF MOUSE CD4 TCELL DN | 2.47E-08 | 5.73E-06 | 7.34E-01 | -0.498 | -1.985 | 182 | Anpep Nid1 Lgals3 Anxa2 Id3 Tgfbi P2rx4 Nqo1 Heg1 Pmp22 Mrc1 Apobec3 Ctsk Plp2 Lmna Hspa1a Stab1 Oasl Sgta S100a4 Emp1 Timp1 Ahnak Lgmn Il1b Eng Cd99 Dab2 Slc3a2 Tnfrsf11a Phldb1 Clic1 Pltp Ifi44 Emp3 Slc7a11 Fcmr Cd163 Cpq Med20 Ifit2 Plin2 Ctsl | 21 |
| GOLDRATH EFF VS MEMORY CD8 TCELL UP | 3.35E-08 | 7.41E-06 | 7.20E-01 | -0.485 | -1.93 | 180 | Tagln2 Lgals1 Fgl2 Lgals3 Anxa2 Kif11 Cdk1 Ect2 Gem Fkbp5 Rom1 S100a10 Anxa1 F2rl3 Litaf S100a4 Emp1 Ube2t Mki67 Kif22 Ifitm3 Cd99 Prc1 Degs1 Serpinb9 Cdca8 Emp3 Nudt4 Nusap1 Tuba1b Dock5 Ezh2 Tk1 Tmem14c Itgax Rrm2 Mt2A Rfc5 Cks2 Dbi Txn1 Tuba1a Cd244 Ccnf Kif4a Cmc2 Lmnb1 Top2a Spdl1 C3 Brca1 Ncaph Pclaf Myadm Syce2 Iqgap3 Anln Dtl Idi1 Bub1 Ybx3 Cdc6 | 22 |
| GSE32901 NAIVE VS TH17 NEG CD4 TCELL UP | 4.65E-08 | 9.79E-06 | 7.20E-01 | -0.523 | -2.03 | 146 | Kif11 Ctsh Ect2 Grn Txndc5 Tpx2 Niban1 Ifi30 Myo1c Trip13 Cenph Mki67 Rasgrp3 Ly86 Cd74 Prc1 Tgm2 Ecm1 Tnfrsf9 Fcmr Cr2 Pros1 Spi1 Cybb Tnfrsf4 Tk1 Hmgn3 Plin2 Odc1 Cenpe Ctnna1 Sdc4 Mvp Pik3c2b S100a6 Nuf2 Ccnf Ehd4 Kif4a Cstb Entpd1 Cd22 Wdfy4 Casp8 Impa2 Cd38 Anln Lpcat3 Ctsz Dtl | 23 |
| GSE21360 PRIMARY VS QUATERNARY MEMORY CD8 TCELL UP | 4.82E-08 | 9.79E-06 | 7.20E-01 | -0.508 | -1.983 | 153 | Anpep Atf3 Lgals3 P2rx4 Nqo1 Pmp22 Igf1 S100a10 Lmna Rnf213 Myof Vwf Emp1 Ahnak Ttyh2 Flna Ifitm3 Cd74 Vat1 Tnfsf4 Penk Adgre5 Ifi44 Myo1f Scpep1 Il2rg St3gal1 Rilpl2 Elf2 Laptm4a Cybb Ptgr1 Ifit2 Anxa5 Odc1 Ly75 Ifit3 Slc7a1 Usp18 Bri3 Rasa4 Mlkl Rsad2 Cstb Entpd1 Clic4 C3 Mthfd2 Slc6a8 Dot1l Serpine1 | 24 |
| GSE19401 NAIVE VS IMMUNIZED MOUSE PLN FOLLICULAR DC UP | 7.01E-08 | 1.37E-05 | 7.05E-01 | -0.488 | -1.945 | 181 | Anpep Atf3 Lgals1 Tgfbi P2rx4 Heg1 Gem Sp110 Mrc1 Tex30 Lmna Hspa1a Stab1 Oasl S100a4 Emp1 Cd4 Hmox1 Arid5a Il1b Eng Cd99 Phldb1 Clic1 Sp100 Pltp Ifi44 Serpinb9 F13a1 Btg2 Emp3 Bcl3 Zc3hav1 Cd163 Prdm1 Rplp0 S100a9 Ctnna1 Itm2a Chn2 Ifit3 P2ry6 Cks2 Crem Zfp263 Plod3 Cemip Clk1 Srsf3 Lamtor5 Serpine1 Ppp1r15a Gna15 Slc7a5 Ssr4 Atp13a3 Asgr2 Macf1 Il3ra Cd86 Npc1 Nfkb1 Eif4a3 | 25 |

**Table S3B. Top 15 Pathways from GSEA (based on p-value) using Gene Sets of MSigDB Hallmark Pathways for PND 46**

| Pathway | pval | padj | log2err | ES | NES | size | Genes | Rank |
| --- | --- | --- | --- | --- | --- | --- | --- | --- |
| HALLMARK EPITHELIAL MESENCHYMAL TRANSITION | 1.17E-24 | 5.85E-23 | 1.29E+00 | -0.674 | -2.691 | 184 | Postn Anpep Tnfrsf11b Lgals1 Cald1 Lama1 Col4a1 Pcolce Tgfbr3 Mfap5 Tgfbi Mylk Pmp22 Fstl1 Col1a1 Col6a3 Gem Itga5 Fbn2 Bgn Ppib Mgp Lum Col12a1 Lox Fbln5 P3h1 Col7a1 Loxl2 Fbn1 Col3a1 Vim Myl9 Lamc1 Tfpi2 Abi3bp Jun Tpm4 Timp1 Calu PCOLCE2 Flna Loxl1 Fn1 Ccn1 Plod1 Nid2 Timp3 Dab2 Tagln Fmod Tgfb1 Eln Acta2 Igfbp2 Pmepa1 Tpm2 Col4a2 Col6a2 Tgm2 Emp3 Ecm1 Sparc Pdgfrb Col5a3 Sfrp1 Ccn2 Dcn Gadd45a Gpx7 Fstl3 Nnmt Lama2 Wnt5a Tnc Gas1 Serpinh1 Sdc4 Col5a1 Colgalt1 Thbs2 Snai2 Cd44 Fbln1 Pvr Itgb1 Plod3 | 1 |
| HALLMARK COAGULATION | 1.68E-07 | 2.98E-06 | 6.90E-01 | -0.561 | -2.063 | 101 | Serping1 Ctsh Gsn Ctsk Anxa1 Cfd Fbn1 Tfpi2 Vwf Plat Timp1 Lgmn Fn1 Rgn Timp3 Maff Pros1 Dct Sparc Cpq Ctsl C1qa Cfh Thbd Csrp1 Furin C3 S100a13 Sirt2 Serpine1 Msrb2 Dpp4 Plek C2 Arf4 Cpn1 Sh2b2 Pdgfb Cd9 C1r Tf Bmp1 | 2 |
| HALLMARK APOPTOSIS | 2.10E-07 | 2.98E-06 | 6.90E-01 | -0.505 | -1.963 | 150 | Atf3 Gch1 Lgals3 Tgfbr3 Txnip Rela Bgn Gsn Lum Lmna Anxa1 Hspb1 Gucy2d Jun Gpx4 Emp1 Plat Timp1 Ifitm3 Hmox1 Il1b Bcl2l11 Casp7 Timp3 Mgmt Btg2 Lef1 Casp1 Cd14 Pdgfrb Ddit3 Gpx1 Dcn Dap Gadd45a Casp4 Pea15 Pmaip1 Tnfsf10 Cd44 Top2a Cdkn1b Tap1 Rhot2 Ccnd2 Cth Tspo Brca1 Casp8 Bax Igfbp6 Birc3 Cd38 Irf1 Rock1 Gna15 Ccna1 Gpx3 Bcl10 Erbb2 | 3 |
| HALLMARK INTERFERON GAMMA RESPONSE | 2.39E-07 | 2.98E-06 | 6.75E-01 | -0.483 | -1.921 | 178 | Plscr1 Fgl2 Gch1 Serping1 RT1-Db1 Txnip Sp110 Rnf213 Ifi30 Vamp8 Oasl Ripk1 Ifitm3 Csf2rb Casp7 Cd74 Ifi44 Casp1 Cxcl10 Parp14 Stat3 RT1-Ba Fcgr1a Ifit2 Vamp5 RT1-A2 Psmb8 B2m Psmb9 Myd88 Isoc1 Lap3 Mvp Mt2A Casp4 Ifit3 Usp18 Cxcl9 Cfh Ciita Sspn Eif4e3 Socs3 Tnfsf10 Rsad2 Tap1 Nod1 Psme1 Ifitm2 Mthfd2 Casp8 RT1-CE4 Il15 Cd38 Irf1 | 4 |
| HALLMARK IL6 JAK STAT3 SIGNALING | 1.95E-06 | 1.95E-05 | 6.27E-01 | -0.591 | -2.043 | 69 | Il2ra Il1r1 Acvrl1 Bak1 Crlf2 Jun Ifnar1 Hmox1 Il1b Csf2rb Lepr Csf1 Tgfb1 Il17ra Il2rg Cxcl10 Cd14 Stat3 Hax1 Myd88 Il10rb Tlr2 Cxcl9 Ltbr Socs3 Tnfrsf1a Cd44 Cxcl13 Ccr1 | 5 |
| HALLMARK P53 PATHWAY | 6.59E-06 | 5.50E-05 | 6.11E-01 | -0.446 | -1.776 | 182 | Atf3 Klf4 Itgb4 Txnip Bak1 Osgin1 Fam162a S100a10 Ptpn14 Ifi30 Vamp8 Zfp36l1 S100a4 Jun Rhbdf2 Stom Hmox1 Gm2a Slc19a2 Slc3a2 Tgfb1 Btg2 Slc7a11 St14 Casp1 Ccnd3 Rrad Tcn2 Ddit3 Epha2 Gadd45a Sphk1 Notch1 Sp1 Tob1 Rgs16 Ephx1 Vdr Cd82 Mxd1 Dcxr H1f2 Phlda3 Kif13b Tap1 Pcna Ccnd2 Tax1bp3 Tgfa Bax Hexim1 Tnfsf9 Hdac3 Ppp1r15a | 6 |
| HALLMARK MYOGENESIS | 1.30E-05 | 9.29E-05 | 5.93E-01 | -0.451 | -1.79 | 176 | Apod Aplnr Itgb4 Mylk Nqo1 Col1a1 Igf1 Col6a3 Casq1 Ldb3 Svil Des Gsn Cfd Myo1c Col3a1 Aebp1 Tnnt3 Tagln Igfbp7 Myh11 Tgfb1 Cox6a2 Tead4 Tpm2 Col4a2 Col6a2 Hrc Smtn Sparc Gja5 Casq2 Sphk1 Lama2 Sh3bgr Notch1 Cryab Pgam2 Sspn Itgb1 Sorbs3 Acsl1 Myom1 Nav2 Slc6a8 Sirt2 Mapk12 Myl4 | 7 |
| HALLMARK INTERFERON ALPHA RESPONSE | 1.72E-05 | 1.08E-04 | 5.76E-01 | -0.529 | -1.918 | 92 | Plscr1 Txnip Sp110 Ifi30 Oasl Ifitm3 Cd74 Cnp Csf1 Ifi44 Gbp2 Casp1 Cxcl10 Parp14 Lpar6 Lamp3 Parp9 Ifit2 RT1-A2 Psmb8 B2m Elf1 Psmb9 Lap3 Ifit3 Mvb12a Usp18 Rsad2 Tap1 Psme1 Ifitm2 Casp8 Trim5 Il15 Irf1 | 8 |
| HALLMARK IL2 STAT5 SIGNALING | 2.96E-05 | 1.65E-04 | 5.76E-01 | -0.429 | -1.705 | 177 | Plscr1 Fgl2 Il2ra Fah P2rx4 Myo1c Abcb1b Gpx4 Emp1 Ahnak Cd48 Ifitm3 Slc39a8 Pth1r Csf1 Ctla4 Il1rl1 Penk Maff Slc1a5 Tgm2 Ecm1 Wls Tnfrsf9 Ccnd3 Cxcl10 Rhoh Serpinb6b Hopx Ahcy Tnfrsf4 St3gal4 Plin2 Odc1 Umps Rnh1 Rgs16 Adam19 Mxd1 Capn3 Tnfsf10 Cd44 Enpp1 Col6a1 Furin Ccnd2 Alcam Muc1 Phlda1 Snx9 Tnfrsf1b Syngr2 Swap70 Ctsz Scn9a Galm Il3ra Cd86 Cdc6 Coch Ager Twsg1 Hk2 Bmp2 Klf6 Socs2 Ikzf4 Cdkn1c Myc Itgae | 9 |
| HALLMARK ANGIOGENESIS | 7.39E-05 | 3.70E-04 | 5.38E-01 | -0.659 | -1.988 | 34 | Postn Jag1 Fstl1 Lum Vtn Col3a1 S100a4 Kcnj8 Timp1 Slco2a1 | 10 |
| HALLMARK ALLOGRAFT REJECTION | 3.60E-04 | 1.64E-03 | 3.27E-01 | -0.426 | -1.654 | 148 | Il2ra Itgal Cd1d1 Fyb1 Krt1 Stab1 Srgn Timp1 RT1-Da Cd4 Flna Ccl19 Was Il1b Ly86 Cd74 Csf1 Tgfb1 Degs1 Itgb2 Ets1 Gbp2 Bcl3 Il2rg Ccnd3 Spi1 RT1-Ba RT1-A2 Nme1 B2m Ly75 Tlr2 Tlr3 Il18rap Cxcl9 Tpd52 Cxcl13 Gcnt1 Ccr1 Ccr5 Tap1 Ccnd2 Brca1 Itk Cd7 Ikbkb Dars1 Il15 C2 Traf2 Stat1 Bcl10 Nck1 Cd86 Map4k1 Ptprc Ifnar2 | 11 |
| HALLMARK CHOLESTEROL HOMEOSTASIS | 4.24E-04 | 1.77E-03 | 3.04E-01 | -0.507 | -1.772 | 73 | Jag1 Atf3 Plscr1 Lgals3 Niban1 Abca2 Lgmn Sema3b Antxr2 Anxa5 Pmvk Cxcl16 S100a11 Tm7sf2 Fbxo6 Alcam Acat2 Idi1 Gpx8 Cyp51 Cd9 | 12 |
| HALLMARK COMPLEMENT | 9.82E-04 | 3.58E-03 | 1.95E-01 | -0.399 | -1.573 | 166 | Plscr1 Lgals3 Gngt2 Serping1 Ctsh Cp Apobec3 Hspa1a Tfpi2 Rhog Plat Timp1 Notch4 Lgmn Fn1 Was Casp7 Maff Col4a2 Casp1 Cr2 Cpq Ctsc Cr1l Anxa5 Ctsl Src Psmb9 Lipa S100a9 Lap3 C1qc Casp4 C1qa Cfh Atox1 Itgam Phex Csrp1 C3 S100a13 Stx4 Serpine1 Dpp4 Irf1 Pla2g7 Plek C2 Gnai2 Cpm C4bpb Prkcd Xpnpep1 Adra2b Usp16 Pdgfb | 13 |
| HALLMARK WNT BETA CATENIN SIGNALING | 1.00E-03 | 3.58E-03 | 1.99E-01 | -0.599 | -1.827 | 36 | Jag1 Axin2 Notch4 Tcf7 Lef1 Nkd1 Ppard Notch1 Wnt6 Wnt5b Ccnd2 Jag2 Fzd1 Myc Skp2 Ncor2 Psen2 Hey2 Ptch1 | 14 |
| HALLMARK INFLAMMATORY RESPONSE | 1.11E-03 | 3.69E-03 | 1.84E-01 | -0.393 | -1.556 | 172 | Aplnr Gch1 Il1r1 P2rx4 Itga5 Rela Stab1 P2rx7 Ccl24 Rhog Ifnar1 Timp1 Sema4d Pdpn Cd48 Ifitm3 Il1b Csf1 Btg2 Emp3 Adora2b Tnfrsf9 Cxcl10 Cd14 Sgms2 Cybb Lamp3 Slc31a2 Sphk1 Tlr2 Slc7a1 Rgs16 Tlr3 Il18rap Cxcl9 Cd82 Mxd1 Tnfsf10 Pvr Scarf1 Axl Ifitm2 Tnfsf9 Serpine1 P2ry2 Il15 Irf1 Gna15 Edn1 Tnfrsf1b | 15 |

**Table S3C. Top 25 Pathways from GSEA (based on p-value) using Gene Sets of MSigDB Canonical Pathways for PND 46**

| Pathway | pval | padj | log2err | ES | NES | size | Genes | Rank |
| --- | --- | --- | --- | --- | --- | --- | --- | --- |
| NABA CORE MATRISOME | 1.08E-30 | 2.00E-27 | 1.44E+00 | -0.683 | -2.78 | 220 | Postn Nyx Fndc1 Col4a5 Fgl2 Nid1 Srpx Lama1 Epyc Col4a1 Pcolce Mfap5 Tgfbi Impg2 Ccn4 Col14a1 Mmrn2 Col1a1 Dpt Col6a3 Lama4 Lgi3 Fbn2 Tinagl1 Col4a6 Bgn Col26a1 Mgp Lum Col12a1 Fbln5 Vtn Lamb1 Col7a1 Col18a1 Sned1 Fbn1 Col3a1 Egflam Aebp1 Lamc1 Abi3bp Emilin1 Col13a1 Colq Vwf Srgn PCOLCE2 Col4a4 Fn1 Ccn1 Nid2 Lama5 Prelp Igfbp7 Fmod Impg1 Eln Igfbp2 Mfap2 Esm1 Optc Col4a2 Col9a2 Col6a2 Cilp Ecm1 Igsf10 Ltbp1 Hapln2 Spon2 Crispld2 Sparc Hmcn1 Col24a1 Lamb2 Dmp1 Npnt Col5a3 Ccn2 Dcn Vwa1 Lrg1 Lama2 Ltbp4 Lamc3 Tnc Col9a1 Col5a1 Fndc7 Papln Vwa5b1 Thbs2 Fbln1 Svep1 Col6a1 Tecta Ccn5 Col2a1 Col4a3 Mfge8 | 1 |
| REACTOME EXTRACELLULAR MATRIX ORGANIZATION | 8.01E-25 | 7.41E-22 | 1.30E+00 | -0.624 | -2.573 | 247 | Col4a5 Nid1 P4hb Lama1 Col4a1 Pcolce Mfap5 Itgb4 Col14a1 Col1a1 Col6a3 Lama4 Crtapl1 Itga5 Fbn2 Adamts1 Col4a6 Bgn Ppib Col26a1 Itgal Lum Ctsk Col12a1 Lox Fbln5 P3h1 Vtn Lamb1 Col7a1 Col18a1 Loxl2 Fbn1 Col3a1 Lamc1 Emilin1 Col13a1 Vwf Timp1 PCOLCE2 Loxl1 Col4a4 Fn1 P4ha2 Plod1 Capn1 Nid2 Lama5 Fmod Tgfb1 Eln Itgb2 Mfap2 Itga2b Adamts14 Optc Col4a2 Col9a2 Col6a2 Ltbp1 Bsg Sparc Scube3 Col24a1 Lamb2 Dmp1 Col5a3 Dcn Ctsl Lama2 Itgax Ltbp4 Lamc3 Tnc Col9a1 Serpinh1 Itga1 Cast Sdc4 Col5a1 Colgalt1 Ddr1 Adam19 Kdr Itgam Capn3 Cd44 Fbln1 Capn6 Itgb1 F11r Plod3 Col6a1 Scube1 Furin Col2a1 Col4a3 Capn10 | 2 |
| REACTOME NEURONAL SYSTEM | 6.62E-22 | 4.08E-19 | 1.21E+00 | 0.534 | 2.353 | 366 | Slitrk2 Adcy1 Prkcb Camk2g Homer1 Kcnh7 Prkca Gabrg2 Kcnab2 Camk2a Kcnb1 Kcnc4 Kcns3 Chrna5 Gad1 Kcnq2 Pdpk1 Adcy2 Grin2a Nrxn1 Kcnq3 Gng7 Gabrb2 Grm5 Cacnb4 Dlgap2 Gabrb1 Gabra1 Slc1a2 Shank1 Gabrb3 Aldh5a1 Lrrc7 Adcy8 Cacng3 Grin2b Cacnb2 Nlgn1 Slitrk1 Dlgap1 Gria2 Lrfn4 Apba2 Gabra5 Gabra2 Kcnj3 Lrrtm3 Kcnc3 Grik3 Gabrg3 Mapk1 Ppm1e Erbb4 Grik2 Gabra3 Kcnh2 Git1 Dlg3 Slitrk5 Il1rapl1 Gad2 Gria1 Hcn1 Slc38a1 Slc1a1 Kcnj6 Gnb5 Nlgn2 Ncald Ap2b1 Sipa1l1 Glrb Prkar1a Plcb1 Gnai1 Prkacb Prkaa2 Dnajc5 Camkk2 Gria3 Gabbr1 Arhgef9 Rasgrf1 Epb41l3 Adcy3 Dlg2 Kcnab1 Lrfn3 Dlg4 Cacna2d3 Homer2 Kcnn1 Nlgn3 Cacng8 Kcna3 Kcnj10 Akap5 Prkx Kcna1 Lin7c Prkar1b Nptn Gria4 Ppfia2 Grip1 Kcng1 Ntrk3 Kcna4 Kcnh4 Kcna2 Kcnn2 Kcnj11 Grik1 Gabrq Nsf Dlgap4 Camk2b Chrna2 Abat Ap2a1 Ppm1f Dlgap3 Rtn3 Shank2 Flot1 Kcnmb2 Glra2 Gnb1 Kcns2 Tspan7 Syt2 Apba1 Cacnb1 | 3 |
| NABA ECM GLYCOPROTEINS | 2.14E-18 | 9.88E-16 | 1.11E+00 | -0.657 | -2.557 | 151 | Postn Fndc1 Fgl2 Nid1 Srpx Lama1 Pcolce Mfap5 Tgfbi Ccn4 Mmrn2 Dpt Lama4 Lgi3 Fbn2 Tinagl1 Mgp Fbln5 Vtn Lamb1 Sned1 Fbn1 Egflam Aebp1 Lamc1 Abi3bp Emilin1 Colq Vwf PCOLCE2 Fn1 Ccn1 Nid2 Lama5 Igfbp7 Eln Igfbp2 Mfap2 Cilp Ecm1 Igsf10 Ltbp1 Spon2 Crispld2 Sparc Hmcn1 Lamb2 Dmp1 Npnt Ccn2 Vwa1 Lrg1 Lama2 Ltbp4 Lamc3 Tnc Fndc7 Papln Vwa5b1 Thbs2 Fbln1 Svep1 Tecta Ccn5 | 4 |
| REACTOME COLLAGEN FORMATION | 3.54E-15 | 1.31E-12 | 1.01E+00 | -0.728 | -2.583 | 80 | Col4a5 P4hb Col4a1 Pcolce Itgb4 Col14a1 Col1a1 Col6a3 Crtapl1 Col4a6 Ppib Col26a1 Col12a1 Lox P3h1 Col7a1 Col18a1 Loxl2 Col3a1 Col13a1 PCOLCE2 Loxl1 Col4a4 P4ha2 Plod1 Adamts14 Col4a2 Col9a2 Col6a2 Col24a1 Col5a3 Ctsl Col9a1 Serpinh1 Col5a1 Colgalt1 Plod3 Col6a1 Col2a1 Col4a3 Dst | 5 |
| PID INTEGRIN1 PATHWAY | 4.64E-15 | 1.43E-12 | 9.97E-01 | -0.776 | -2.633 | 62 | Col4a5 Nid1 Lama1 Col4a1 Tgfbi Col1a1 Col6a3 Lama4 Itga5 Col4a6 Vtn Lamb1 Col7a1 Col18a1 Fbn1 Col3a1 Lamc1 Col4a4 Fn1 Lama5 F13a1 Col6a2 Tgm2 Cd14 Lamb2 Npnt Lama2 Tnc Itga1 Col5a1 Thbs2 Itgb1 Col6a1 Col2a1 Col4a3 Cspg4 | 6 |
| REACTOME DEGRADATION OF THE EXTRACELLULAR MATRIX | 3.72E-14 | 9.52E-12 | 9.65E-01 | -0.672 | -2.495 | 108 | Col4a5 Nid1 Col4a1 Col14a1 Col1a1 Col6a3 Fbn2 Adamts1 Col4a6 Col26a1 Ctsk Col12a1 Lamb1 Col7a1 Col18a1 Fbn1 Col3a1 Lamc1 Col13a1 Timp1 Col4a4 Fn1 Capn1 Lama5 Eln Optc Col4a2 Col9a2 Col6a2 Bsg Scube3 Col5a3 Dcn Ctsl Col9a1 Cast Col5a1 Capn3 Cd44 Capn6 Col6a1 Scube1 Furin Col2a1 Col4a3 Capn10 | 7 |
| NABA MATRISOME ASSOCIATED | 4.12E-14 | 9.52E-12 | 9.65E-01 | -0.467 | -2.028 | 429 | Angptl2 Ptn Clec3b Pdgfd Lgals1 Lgals3 Megf6 Anxa2 Il34 Pappa2 Serping1 Egfl7 Megf8 Anxa6 Ctsh Bmp6 Fstl1 Igf1 C1qtnf7 Cela3b Adamts1 Ctsk Lox Adam2 Clec10a S100a10 Gdf7 Bmp15 P3h1 Angpt2 Nrtn Anxa1 Tgm3 Loxl2 Ccl24 Frzb S100a4 Adamts12 Tnfsf12 Plat Pamr1 Timp1 Sema4d Igf2 Adamts15 Ccl19 Loxl1 Il1b P4ha2 Plod1 Htra3 Wnt2b Timp3 Clec14a Csf1 Gdf10 Sema3b Tnfsf4 Tgfb1 Adamts14 Serpinb9 F13a1 Tgm2 St14 Cd109 Cxcl10 Scube3 Ctsa Serpinb6b Kazald1 Ctsc Serpinf1 Adamtsl2 Pgf Fgf1 Sfrp1 Clec12a Anxa5 Adamts7 Adamts10 Ctsl Bmp5 Serpinf2 Fstl3 Anxa3 Wnt5a S100a9 Serpinh1 Sdc4 C1qc Anxa8 Dhh S100a6 Cxcl9 C1qa Adam19 Sema3a S100a11 Wnt6 C1qtnf6 Tnfsf10 Hyal2 Colec12 Cxcl13 Sema3g Cstb Plod3 Serpind1 Cela1 Scube1 Hcfc2 Wnt5b Vegfc Fgf22 Cspg4 Tgfa Wnt3 Wfikkn2 Adamtsl4 S100a13 Egfl8 Mmp23 Tnfsf9 Serpinb1a Serpine1 Adam32 Muc1 Il15 Lman1 Mmp19 Agt Cst6 Wnt9b Clec7a | 8 |
| REACTOME INTEGRIN CELL SURFACE INTERACTIONS | 1.15E-13 | 2.36E-11 | 9.44E-01 | -0.734 | -2.564 | 73 | Col4a5 Col4a1 Col1a1 Col6a3 Itga5 Col4a6 Itgal Lum Vtn Col7a1 Col18a1 Fbn1 Col3a1 Col13a1 Vwf Col4a4 Fn1 Itgb2 Itga2b Col4a2 Col9a2 Col6a2 Bsg Col5a3 Itgax Tnc Col9a1 Itga1 Col5a1 Kdr Itgam Cd44 Itgb1 F11r Col6a1 Col2a1 Col4a3 | 9 |
| REACTOME TRANSMISSION ACROSS CHEMICAL SYNAPSES | 1.19E-12 | 2.21E-10 | 9.10E-01 | 0.516 | 2.176 | 240 | Adcy1 Prkcb Camk2g Prkca Gabrg2 Camk2a Chrna5 Gad1 Pdpk1 Adcy2 Grin2a Gng7 Gabrb2 Cacnb4 Gabrb1 Gabra1 Slc1a2 Gabrb3 Aldh5a1 Lrrc7 Adcy8 Cacng3 Grin2b Cacnb2 Gria2 Gabra5 Gabra2 Kcnj3 Grik3 Gabrg3 Mapk1 Ppm1e Erbb4 Grik2 Gabra3 Git1 Dlg3 Gad2 Gria1 Slc38a1 Slc1a1 Kcnj6 Gnb5 Ncald Ap2b1 Glrb Prkar1a Plcb1 Gnai1 Prkacb Prkaa2 Dnajc5 Camkk2 Gria3 Gabbr1 Arhgef9 Rasgrf1 Adcy3 Dlg2 Dlg4 Cacna2d3 Cacng8 Kcnj10 Akap5 Prkx Lin7c Prkar1b Nptn Gria4 Ppfia2 Grip1 Grik1 Gabrq Nsf Camk2b Chrna2 Abat Ap2a1 Ppm1f | 10 |
| REACTOME COLLAGEN BIOSYNTHESIS AND MODIFYING ENZYMES | 3.45E-12 | 5.81E-10 | 8.99E-01 | -0.741 | -2.52 | 63 | Col4a5 P4hb Col4a1 Pcolce Col14a1 Col1a1 Col6a3 Crtapl1 Col4a6 Ppib Col26a1 Col12a1 P3h1 Col7a1 Col18a1 Col3a1 Col13a1 PCOLCE2 Col4a4 P4ha2 Plod1 Adamts14 Col4a2 Col9a2 Col6a2 Col24a1 Col5a3 Col9a1 Serpinh1 Col5a1 Colgalt1 Plod3 Col6a1 Col2a1 Col4a3 | 11 |
| REACTOME NEUROTRANSMITTER RECEPTORS AND POSTSYNAPTIC SIGNAL TRANSMISSION | 5.39E-12 | 8.30E-10 | 8.87E-01 | 0.549 | 2.238 | 180 | Adcy1 Prkcb Camk2g Prkca Gabrg2 Camk2a Chrna5 Pdpk1 Adcy2 Grin2a Gng7 Gabrb2 Gabrb1 Gabra1 Gabrb3 Lrrc7 Adcy8 Cacng3 Grin2b Gria2 Gabra5 Gabra2 Kcnj3 Grik3 Gabrg3 Mapk1 Ppm1e Erbb4 Grik2 Gabra3 Git1 Dlg3 Gria1 Kcnj6 Gnb5 Ncald Ap2b1 Glrb Prkar1a Plcb1 Gnai1 Prkacb Prkaa2 Camkk2 Gria3 Gabbr1 Arhgef9 Rasgrf1 Adcy3 Dlg2 Dlg4 Cacng8 Kcnj10 Akap5 Prkx Lin7c Prkar1b Nptn Gria4 Grip1 Grik1 Gabrq Nsf Camk2b Chrna2 Ap2a1 Ppm1f Glra2 Gnb1 Tspan7 Apba1 | 12 |
| REACTOME ASSEMBLY OF COLLAGEN FIBRILS AND OTHER MULTIMERIC STRUCTURES | 9.08E-12 | 1.29E-09 | 8.75E-01 | -0.748 | -2.467 | 53 | Col4a5 Col4a1 Pcolce Itgb4 Col14a1 Col1a1 Col6a3 Col4a6 Col12a1 Lox Col7a1 Col18a1 Loxl2 Col3a1 Loxl1 Col4a4 Col4a2 Col9a2 Col6a2 Col24a1 Col5a3 Ctsl Col9a1 Col5a1 Col6a1 Col2a1 Col4a3 Dst | 13 |
| WP FRAGILE X SYNDROME | 2.22E-11 | 2.93E-09 | 8.63E-01 | 0.607 | 2.337 | 117 | Homer1 Prkca Gabrg2 Camk2a Ppp3ca Gad1 Grin2a Gabrb2 Grm5 Syngap1 Gabra1 Cyfip2 Shank1 Aldh5a1 Grin2b Eif4e Gria2 Gphn Mapk1 Gria1 Hcn1 Grb2 Itpr1 Rptor Mecp2 Ap2b1 Prkar1a Plcb1 Pik3cb Epha4 Dlg4 Cyfip1 Dnm1 Akap5 Grip1 Camk2b Abat Ap2a1 Dlgap3 Cnr1 Mlst8 Tti1 Akt1 Nf1 Grin1 Rps6kb1 Mtor Telo2 Akt1s1 Cltc App Eef1a1 Braf Kras Bdnf Ppp1ca Ago2 Arc Map2k1 Arhgap32 | 14 |
| REACTOME ECM PROTEOGLYCANS | 3.96E-11 | 4.88E-09 | 8.51E-01 | -0.69 | -2.392 | 70 | Col4a5 Lama1 Col4a1 Col1a1 Col6a3 Lama4 Col4a6 Bgn Lum Vtn Lamb1 Col3a1 Lamc1 Col4a4 Fn1 Lama5 Fmod Tgfb1 Itga2b Col4a2 Col9a2 Col6a2 Sparc Lamb2 Dmp1 Col5a3 Dcn Lama2 Itgax Tnc Col9a1 Col5a1 Itgb1 Col6a1 Col2a1 Col4a3 | 15 |
| KEGG ECM RECEPTOR INTERACTION | 4.40E-11 | 5.09E-09 | 8.51E-01 | -0.686 | -2.402 | 74 | Lama1 Col4a1 Itgb4 Col1a1 Col6a3 Lama4 Itga5 Col4a6 Vtn Lamb1 Col3a1 Lamc1 Vwf Col4a4 Fn1 Lama5 Itga2b Col4a2 Col6a2 Lamb2 Col5a3 Lama2 Lamc3 Tnc Itga1 Sdc4 Col5a1 Thbs2 Cd44 Itgb1 Col6a1 Col2a1 | 16 |
| REACTOME LAMININ INTERACTIONS | 5.64E-11 | 6.14E-09 | 8.51E-01 | -0.839 | -2.464 | 30 | Col4a5 Nid1 Lama1 Col4a1 Itgb4 Lama4 Col4a6 Lamb1 Col7a1 Col18a1 Lamc1 Col4a4 Nid2 Lama5 Col4a2 Lamb2 Lama2 Lamc3 Itga1 | 17 |
| NABA BASEMENT MEMBRANES | 7.83E-11 | 8.05E-09 | 8.39E-01 | -0.8 | -2.467 | 38 | Col4a5 Nid1 Lama1 Col4a1 Col6a3 Lama4 Col4a6 Lamb1 Col18a1 Lamc1 Colq Col4a4 Nid2 Lama5 Col4a2 Col6a2 Hmcn1 Lamb2 Npnt Lama2 Lamc3 Papln Col6a1 Col4a3 | 18 |
| REACTOME NEUTROPHIL DEGRANULATION | 1.11E-10 | 1.08E-08 | 8.39E-01 | -0.44 | -1.904 | 404 | Anpep P2rx1 Fgl2 Lgals3 Anxa2 Frk Ctsh Grn Chrnb4 Vcl Txndc5 Lyz2 Rhoa Folr2 Gyg1 Unc13d Gsn Itgal Gstp1 Ist1 Hspa1a Krt1 Cfd Vamp8 Rhog Ostf1 Arhgap45 Tcirg1 Stom Galns Dpp7 Gla Cd63 Gm2a Capn1 Gmfg Hspa8 Vat1 Degs1 Itgb2 Pkp1 Adgre5 Srp14 Magt1 Pgm2 Cd93 Crispld2 Cyb5r3 Tmbim1 Cd14 Padi2 Mgst1 Ctsa Ticam2 Ap1m1 Serpinb6b Ctsc Cybb Ftl1 Cnn2 Snap29 Iqgap1 Dera Clec12a RT1-A2 Cr1l Ptprb Ano6 B2m Lrg1 Npc2 Itgax Tmem179b Aga Xrcc5 S100a9 Surf4 Pgrmc1 Mvp Rnaset2 Hbb Tlr2 Oscar Cd68 Chit1 Rab27a Bri3 Nbeal2 S100a11 Itgam Glb1 Cd300a Tubb5 Cd44 B4galt1 Slc44a2 Gsdmd Cstb Cpped1 Gpr84 Mettl7a Tmem63a Cpne3 Hexb Cyba Idh1 C3 Dynlt1 | 19 |
| PID INTEGRIN3 PATHWAY | 1.55E-09 | 1.43E-07 | 7.88E-01 | -0.771 | -2.367 | 37 | Col4a5 Col4a1 Tgfbi Col1a1 Lama4 Col4a6 Vtn Lamb1 Fbn1 Lamc1 Col4a4 Fn1 Ccn1 Itga2b Pdgfrb Sphk1 Tnc Sdc4 Kdr Pvr F11r Col4a3 | 20 |
| WP GABA RECEPTOR SIGNALING | 1.67E-09 | 1.47E-07 | 7.88E-01 | 0.813 | 2.427 | 30 | Gabrg2 Gad1 Gabrb2 Gabrb1 Gabra1 Gabrb3 Gabra5 Gabra2 Gphn Gabrg1 Gabrg3 Gabra3 Gad2 Ap2b1 Gabbr1 Gabrq Abat Ap2a1 | 21 |
| REACTOME PROTEIN PROTEIN INTERACTIONS AT SYNAPSES | 3.32E-09 | 2.79E-07 | 7.75E-01 | 0.624 | 2.272 | 83 | Slitrk2 Homer1 Grin2a Nrxn1 Grm5 Dlgap2 Shank1 Grin2b Nlgn1 Slitrk1 Dlgap1 Lrfn4 Apba2 Lrrtm3 Dlg3 Slitrk5 Il1rapl1 Gria1 Nlgn2 Sipa1l1 Gria3 Epb41l3 Dlg2 Lrfn3 Dlg4 Homer2 Nlgn3 Lin7c Gria4 Ppfia2 Ntrk3 Dlgap4 Dlgap3 Rtn3 Shank2 Flot1 Syt2 Apba1 Grin1 Syt1 Syt10 | 22 |
| REACTOME COLLAGEN DEGRADATION | 3.68E-09 | 2.96E-07 | 7.75E-01 | -0.72 | -2.339 | 49 | Col4a5 Col4a1 Col14a1 Col1a1 Col6a3 Col4a6 Col26a1 Ctsk Col12a1 Col7a1 Col18a1 Col3a1 Col13a1 Col4a4 Col4a2 Col9a2 Col6a2 Col5a3 Ctsl Col9a1 Col5a1 Col6a1 Furin Col2a1 Col4a3 | 23 |
| NABA ECM REGULATORS | 3.96E-09 | 3.05E-07 | 7.62E-01 | -0.545 | -2.106 | 144 | Pappa2 Serping1 Ctsh Cela3b Adamts1 Ctsk Lox Adam2 P3h1 Tgm3 Loxl2 Adamts12 Plat Pamr1 Timp1 Adamts15 Loxl1 P4ha2 Plod1 Htra3 Timp3 Adamts14 Serpinb9 F13a1 Tgm2 St14 Cd109 Ctsa Serpinb6b Kazald1 Ctsc Serpinf1 Adamtsl2 Adamts7 Adamts10 Ctsl Serpinf2 Serpinh1 Adam19 Hyal2 Cstb Plod3 Serpind1 Cela1 Adamtsl4 Mmp23 Serpinb1a Serpine1 Adam32 Mmp19 Agt Cst6 Ctsz Fam20a Adamtsl5 Plod2 Hyal1 Adamts2 Bmp1 | 24 |
| KEGG LONG TERM POTENTIATION | 9.13E-09 | 6.76E-07 | 7.48E-01 | 0.67 | 2.34 | 65 | Adcy1 Prkcb Camk2g Gnaq Prkca Camk2a Ppp3ca Grin2a Grm5 Adcy8 Grin2b Gria2 Mapk1 Ppp3r1 Gria1 Itpr1 Calm2 Plcb1 Prkacb Prkx Camk2b Ppp1cb Ppp3cb Grin1 Calm3 Nras Braf Rps6ka6 Kras Ppp1ca Calm1 Camk2d Map2k1 Prkcg Rps6ka1 Plcb2 Camk4 Rps6ka3 Grin2c | 25 |

**Table S3D. Top 25 Pathways from GSEA (based on p-value) using Gene Sets of MSigDB GO Annotations for PND 46**

| Pathway | pval | padj | log2err | ES | NES | size | Genes | Rank |
| --- | --- | --- | --- | --- | --- | --- | --- | --- |
| GOCC COLLAGEN CONTAINING EXTRACELLULAR MATRIX | 1.59E-42 | 7.78E-39 | 1.70E+00 | -0.677 | -2.865 | 319 | Postn Angptl2 Ptn Clec3b Lgals1 Plscr1 Col4a5 Fgl2 Nid1 Lgals3 Srpx Anxa2 Lama1 Col4a1 Pcolce Serping1 Mfap5 Itgb4 Tgfbi Impg2 Egfl7 Anxa6 Ctsh Col14a1 Mmrn2 Col1a1 Rarres2 Dpt Col6a3 Lama4 Fbn2 Adamts1 Tinagl1 Col4a6 Bgn Col26a1 Mgp Lum Col12a1 Fbln5 S100a10 P3h1 Vtn Krt1 Lamb1 Angpt2 Col7a1 Anxa1 Col18a1 Loxl2 Fbn1 Col3a1 Egflam Aebp1 S100a4 Lamc1 Abi3bp Emilin1 Col13a1 Colq Vwf Bcam Timp1 Adamts15 Loxl1 Col4a4 Fn1 Ccn1 Rbp3 Nid2 Lama5 Wnt2b Timp3 Prelp Clec14a Igfbp7 Gdf10 Fmod Sema3b Tgfb1 Impg1 Eln Mfap2 Myoc Serpinb9 Col4a2 F13a1 Col9a2 Col6a2 Tgm2 Cilp Ecm1 Ltbp1 Sparc Hmcn1 Serpinb6b Kazald1 Ctsc Col24a1 Serpinf1 Lamb2 Npnt Col5a3 Sfrp1 Ccn2 Dcn Vwa1 Anxa5 Adamts10 Ctsl Serpinf2 Lama2 Ltbp4 Lamc3 Wnt5a Tnc S100a9 Col9a1 Serpinh1 Col5a1 C1qc Ssc5d Anxa8 S100a6 C1qa Adam19 Thbs2 Fbln1 Cstb Plod3 Col6a1 Tecta Ccdc80 Col2a1 Wnt5b Col4a3 Mfge8 Cspg4 Dst Adamtsl4 Scp2 Mmp23 Serpinb1a Serpine1 | 1 |
| GOCC EXTERNAL ENCAPSULATING STRUCTURE | 1.77E-41 | 4.34E-38 | 1.68E+00 | -0.627 | -2.723 | 424 | Postn Angptl2 Ptn Nyx Clec3b Tnfrsf11b Lgals1 Plscr1 Col4a5 Fgl2 Nid1 Lgals3 Srpx Anxa2 Lama1 Epyc Col4a1 Pcolce Serping1 Tgfbr3 Mfap5 Itgb4 Tgfbi Impg2 Egfl7 Lrrc17 Anxa6 Ccn4 Ctsh Col14a1 Mmrn2 Col1a1 Rarres2 Dpt Col6a3 Lama4 Fbn2 Adamts1 Tinagl1 Col4a6 Bgn Col26a1 Mgp Lrrc32 Lum Col12a1 Lox Fbln5 S100a10 P3h1 Vtn Krt1 Lamb1 Angpt2 Col7a1 Anxa1 Col18a1 Olfml2a Loxl2 Fbn1 Col3a1 Egflam Aebp1 S100a4 Lamc1 Tfpi2 Abi3bp Emilin1 Col13a1 Colq Vwf Adamts12 Bcam Timp1 Adamts15 Loxl1 Col4a4 Fn1 Pkhd1l1 Ccn1 Ucma Rbp3 Nid2 Lama5 Wnt2b Timp3 Prelp Clec14a Igfbp7 Gdf10 Fmod Sema3b Tgfb1 Impg1 Eln Lingo4 Mfap2 Myoc Adamts14 Serpinb9 Optc Col4a2 F13a1 Col9a2 Col6a2 Tgm2 Cilp Ecm1 Ltbp1 Hapln2 Spon2 Crispld2 Sparc Hmcn1 Serpinb6b Kazald1 Ctsc Col24a1 Serpinf1 Lamb2 Adamtsl2 Dmp1 Fgf1 Npnt Col5a3 Sfrp1 Ccn2 Dcn Vwa1 Anxa5 Adamts7 Adamts10 Ctsl Serpinf2 Lama2 Ltbp4 Lamc3 Wnt5a Tnc S100a9 Col9a1 Serpinh1 Col5a1 C1qc Ssc5d Anxa8 Tlr3 S100a6 Papln C1qa Adam19 Wnt6 Thbs2 Fbln1 Colec12 Cstb Plod3 Vasn Col6a1 Tecta Ccn5 Ccdc80 Col2a1 Wnt5b Col4a3 Mfge8 Cspg4 Dst Wnt3 Adamtsl4 Lrig3 Scp2 Mmp23 Serpinb1a Serpine1 | 2 |
| GOMF EXTRACELLULAR MATRIX STRUCTURAL CONSTITUENT | 5.28E-32 | 8.62E-29 | 1.47E+00 | -0.783 | -2.983 | 129 | Postn Col4a5 Fgl2 Nid1 Srpx Lama1 Col4a1 Pcolce Mfap5 Tgfbi Impg2 Col14a1 Mmrn2 Col1a1 Dpt Col6a3 Lama4 Fbn2 Tinagl1 Col4a6 Bgn Mgp Lum Col12a1 Fbln5 Vtn Lamb1 Col7a1 Col18a1 Fbn1 Col3a1 Aebp1 Lamc1 Tfpi2 Abi3bp Emilin1 Col13a1 Colq Vwf Cd4 Col4a4 Fn1 Ccn1 Nid2 Lama5 Prelp Igfbp7 Fmod Impg1 Eln Mfap2 Optc Col4a2 Col9a2 Col6a2 Cilp Ecm1 Ltbp1 Sparc Hmcn1 Col24a1 Lamb2 Npnt Col5a3 Dcn Vwa1 Lama2 Ltbp4 Tnc Umodl1 Col9a1 Col5a1 | 3 |
| GOBP EXTERNAL ENCAPSULATING STRUCTURE ORGANIZATION | 5.73E-31 | 7.02E-28 | 1.45E+00 | -0.617 | -2.619 | 329 | Postn Tnfrsf11b Col4a5 Nid1 Lama1 Tie1 Col4a1 Mfap5 Itgb4 Tgfbi Col14a1 Cyp1b1 Col1a1 Dpt Col6a3 Lama4 Foxf1 Crtapl1 Itga5 Fbn2 Adamts1 Foxc1 Col4a6 Bgn Itgal Lum Ctsk Col12a1 Lox Fbln5 Vtn Lamb1 Col7a1 Col18a1 Olfml2a Tgm3 Loxl2 Fbn1 Col3a1 Egflam Nphs1 Aebp1 Lamc1 Emilin1 Col13a1 Colq Vwf Adamts12 Timp1 Pdpn Adamts15 Loxl1 Col4a4 Fn1 Ccn1 Eng Capn1 Slc39a8 Nid2 Lama5 Fmod Myh11 Tgfb1 Phldb1 Eln Itgb2 Ets1 Mfap2 Itga2b Fkbp10 Adamts14 Optc Col4a2 Col9a2 Col6a2 Gas2 Hapln2 Bcl3 Crispld2 Bsg Sparc Scube3 Kazald1 Col24a1 Lamb2 Adamtsl2 Dmp1 Npnt Col5a3 Ccn2 Dcn Vwa1 Adamts7 Adamts10 Ctsl Serpinf2 Lama2 Itgax Lamc3 Tnc Phldb2 Notch1 Col9a1 Serpinh1 Itga1 Reck Col5a1 Fgfr4 Colgalt1 Ddr1 Papln Adam19 Kdr P3h4 Itgam Tnfrsf1a Cd44 Fbln1 B4galt1 Itgb1 F11r Plod3 Col6a1 Ccdc80 Scube1 Furin Col2a1 | 4 |
| GOCC NEURON TO NEURON SYNAPSE | 2.50E-24 | 2.45E-21 | 1.28E+00 | 0.575 | 2.507 | 327 | Adcy1 Homer1 Drp2 Dgki Magi2 Lrrc4c Camk2a Actr2 Ppp1r9a Ptprt Pdpk1 Shisa7 Grin2a Grm5 Vamp7 Add2 Syngap1 Pak3 Adora1 Adgrl1 Dlgap2 Lrfn5 Sorcs3 Pja2 Shank1 Slc8a1 Add1 Ntsr1 Epha7 Lrrc7 Adcy8 ND2 Cacng3 Grid1 Sh2d5 Grin2b Nlgn1 Slitrk1 Dlgap1 Gria2 Rapgef4 Lrfn4 Grm3 Inpp4a Lrrtm3 Gphn Lrp4 Mapk1 Erbb4 Grm7 Anks1b Grik2 Nck2 Arhgap44 Lrrc4 Mx2 Bcl11a Cdkl5 Gria1 Dab1 Atp2b2 Slc1a1 Itpr1 Slc8a2 Chrm1 Dclk1 Cap2 Tnik Adam22 Bmpr2 Nlgn2 Sipa1l1 Dnajc6 Clstn1 Shisa8 Dagla Ppp1r9b Lzts3 Arhgef9 Dlg2 Mapk8ip2 Epha4 Lrfn3 Rgs7bp Dlg4 Homer2 Ctnnd2 Nlgn3 Dnm1 Cacng8 Akap5 Lin7c Shisa6 Prkar1b Nptn Rgs14 Abi1 Cdk5r1 Ptk2b Psd3 Arrb1 Dcc Grip1 Cpeb4 Insyn1 Hnrnph2 Plppr4 Nsf Drosha Cnih2 Nsmf Mink1 Arf1 Dlgap3 Srcin1 Shank2 Iqsec3 Dmtn Neto2 Lrp8 Rtn4 Slc4a8 Grin1 Syt1 Kcnh1 Neto1 Baalc Dnm3 Grid2 Septin11 Cacng7 Actn2 Stxbp5 Iqsec1 Igsf9b Lrrtm2 Zdhhc2 Atp1a1 Gopc Dbn1 Psd Crtc1 Tamalin Syn1 Tmem108 Slitrk3 Efnb3 Lrfn1 Shank3 Zdhhc15 Syn3 Mib1 Tiam1 Prnp Arc Arhgap32 Adgra1 Ncs1 Syndig1 Ptprd Atp1a3 Calb1 Nefh Mpp2 Disc1 Oprd1 Strn Syt7 Plekha5 Syn2 Prkcg | 5 |
| GOCC SYNAPTIC MEMBRANE | 6.19E-21 | 5.06E-18 | 1.19E+00 | 0.535 | 2.344 | 345 | Slitrk2 Adcy1 Dgki Lrrc4c Gabrg2 Kcnb1 Cdh10 Atp2b1 Chrna5 Ptprt Shisa7 Grin2a Nrxn1 Gabrb2 Grm5 Stx1b Cntnap4 Adora1 Adgrl1 Gabrb1 Dnm1l Lrfn5 Sorcs3 Gabra1 Slc1a2 Shank1 Slc8a1 Gabrb3 Epha7 Adcy8 Cacng3 Grid1 Grin2b Nlgn1 Slitrk1 Gria2 Lrfn4 Grm3 Ddn Gabra5 Gabra2 Kcnj3 Lrrtm3 Ank1 Kcnc3 Grik3 Gphn Gabrg1 Gabrg3 Lrp4 Erbb4 Grm7 Grik2 Gabra3 Unc13a Lrrc4 Mx2 Il1rapl1 Gad2 Gria1 Dgkb Htr1b Gpm6a Atp2b2 Chrm1 Adam22 Nlgn2 Glrb Clstn1 Adra1a Shisa8 Adgrl2 Dagla Gria3 Chrm3 Ppp1r9b Gabbr1 Cdh9 Dlg2 Epha4 Lrfn3 Rgs7bp Dlg4 Nlgn3 Dnm1 Ntng2 Cacng8 Kcna3 Dnaja3 Kcna1 Lin7c Shisa6 Nptn Htr2a Arrb1 Gria4 Pcdh8 Itsn1 Snph Dcc Zdhhc17 Exoc3 Grip1 Magee1 Picalm Kcna2 Grik1 Gabrq Plppr4 Ptpra Chrna2 Cnih2 Pi4k2a Sncaip Shank2 Crkl Cnr1 Adam23 Glra2 Erc2 Apba1 Scn2a Rph3a Slc6a5 Slc4a8 Chrm5 Grin1 Syt1 Kcnh1 Neto1 Dnm3 Chrm2 Grid2 Kcnma1 Cacng7 Actn2 Stxbp5 Cbln1 | 6 |
| GOBP REGULATION OF TRANS SYNAPTIC SIGNALING | 2.30E-20 | 1.61E-17 | 1.17E+00 | 0.518 | 2.292 | 386 | Adcy1 Prkcb Homer1 Dgki Lrrc4c Camk2a Kcnb1 Ppp1r9a Ppp3ca Pnkd Chrna5 Jph3 Iqsec2 Cx3cl1 Lrrk2 Shisa7 Grin2a Nrxn1 Grm5 Stx1b Syngap1 Cntnap4 Adora1 Dlgap2 Dnm1l Sorcs3 Npy5r Shank1 Usp46 Btbd9 Ophn1 Adcy8 Cacng3 Grid1 Grin2b Eif4e Cacnb2 Prepl Nlgn1 Dlgap1 Grm3 Grik3 Mapk1 Grm7 Grik2 Unc13a Git1 Lrrc4 Plcl2 Gria1 Dgkb Htr1b Prkce Slc1a1 Slc8a2 Rab5al1 Adcyap1 Mecp2 Nlgn2 Sipa1l1 Ncdn Eif2ak4 Clstn1 Calm2 Plcb1 Gnai1 Adra1a Shisa8 Itpka Phf24 Ppp1r9b Rasgrf1 Kctd13 Stau1 Mapk8ip2 Epha4 Dlg4 Cyfip1 Nlgn3 Ntng2 Cacng8 Kcnj10 Akap5 Egfr Abr Hcrt Mctp1 Shisa6 Prkar1b Nptn Rgs14 Ptk2b Htr2a Jph4 Ppfia2 Cplx2 Lilrb3 Dcc Neurod2 Tshz3 Cntn4 Grik1 Dlgap4 Ptpra Camk2b Mme Cnih2 Nsmf Slc4a10 Arf1 Dlgap3 Ythdf1 Rapgef2 Sncaip Atp2a2 Shank2 Flot1 Cnr1 Celf4 Kif5b Gsk3b Cux2 Brsk1 Rab11a Lrp8 Plk2 Erc2 Stxbp5l Apba1 Cyp46a1 Slc4a8 Cdh11 Fgf14 Ppp3cb Tnr Grin1 Syt1 Calm3 Pink1 Neto1 Mtor Ntf3 Anapc2 Grid2 Psmc5 Cacng7 Stxbp5 Cbln1 Asic1 Tubb2b App Car7 Nrg3 Lrrtm2 Zdhhc2 Ache Dbn1 Kras Rims1 Crtc1 Bdnf Lgi1 Syn1 Tmem108 Syngr1 Wnt7a Shank3 Syn3 Pcdh17 Grm8 Calm1 Arc Dgke Sqstm1 Tacr2 Ncs1 Vgf Nsg1 Ptprd Calb1 Pten | 7 |
| GOCC GLUTAMATERGIC SYNAPSE | 2.53E-18 | 1.55E-15 | 1.11E+00 | 0.537 | 2.32 | 297 | Slitrk2 Adcy1 Homer1 Drp2 Lrrc4c Ppp1r9a Ppp3ca Cdh10 Atp2b1 Ptprt Lrrk2 Shisa7 Grin2a Syngap1 Pak3 Prkci Dlgap2 Lrfn5 Sorcs3 Napa Ppm1h Usp46 Ophn1 Epha7 Adcy8 Cacng3 Grid1 Eif4e Nlgn1 Slitrk1 Dlgap1 Lrfn4 Grm3 Lrrtm3 Grik3 Elavl1 Sptbn2 Erbb4 Grik2 Cadps2 Arhgap44 Lrrc4 Il1rapl1 Cdkl5 Gria1 Dgkb Gpm6a Atp2b2 Chrm1 Tnik Adam22 Ap2b1 Clstn1 Prkar1a Plcb1 Adra1a Adgrl2 Stau1 Epha4 Lrfn3 Rgs7bp Dlg4 Homer2 Nlgn3 Ntng2 Kcna3 Cadps Abr Hpca Eps15 Arpc2 Kcna1 Lin7c Shisa6 Prkar1b Nptn Rgs14 Cttnbp2 Nos1ap Ptk2b Camkv Htr2a Ppfia2 Cplx2 Pcdh8 Itsn1 Grip1 Plppr4 Dlgap4 Bcan Cnih2 Arf1 Dlgap3 Flot1 Sv2a Cnr1 Adam23 Gsk3b Rab11a Erc2 Apba1 Scn2a Caly Slc4a8 Cdh11 Cbln2 Ppp3cb Ywhae Tnr Syt1 Clcn3 Neto1 Mtor Wasf3 Dnm3 Ppp2r2a Grid2 Septin11 Cacng7 Actn2 Cbln1 Nrg3 Lrrtm2 Dbn1 Crtc1 Tamalin Lgi1 Ppp1r1b Efnb3 Wnt7a Psd2 Vcp Syn3 Tiam1 Pcdh17 Ppp1ca Arc Dgke Adgra1 Ncs1 Adgrl3 Nsg1 Ptprd Calb1 Rac1 Ywhaz | 8 |
| GOBP SYNAPSE ORGANIZATION | 6.37E-18 | 3.47E-15 | 1.10E+00 | 0.496 | 2.196 | 387 | Slitrk2 Kalrn Homer1 Drp2 Abi2 Lrrc4c Gabrg2 Actr2 Ppp1r9a Ptprt Frmpd4 Cx3cl1 Lrrk2 Shisa7 Nrxn1 Gabrb2 Grm5 Add2 Cacnb4 Dhx36 Syngap1 Pak3 Wasl Adgrl1 Dnm1l Lrfn5 Opa1 Lrrn3 Gabra1 Sema3e Shank1 Gabrb3 Kif1a Ophn1 Vps35 Epha7 Grin2b Cacnb2 Nlgn1 Slitrk1 Lrfn4 Gabra2 Lrrtm3 Pcdhb16l Gphn Lrp4 Sptbn2 Erbb4 Unc13a Arhgap44 Slitrk5 Flrt2 Lrrc4 Il1rapl1 Cdkl5 Dgkb Gpm6a Slc8a2 Mecp2 Pcdhb2 Ube3a Nlgn2 Sipa1l1 Wnt7b Elfn1 Adgrb3 Glrb Myh10 Clstn1 Lgi2 Adgrl2 Itpka Lrrn1 Nrcam Lzts3 Klk8 Cdh9 Stau1 Epha4 Lrfn3 Dlg4 Pcdhb21 Cyfip1 Ctnnd2 Amigo1 Nlgn3 Ntng2 Dnaja3 Shisa6 Nptn Sybu Cttnbp2 Nos1ap Cdk5r1 Camkv Ppfia2 Pcdh8 Itsn1 Lilrb3 Cbln4 Neurod2 Zfp365 Bcan Robo2 Camk2b Arhgap33 Setd5 Arf1 Dlgap3 Srcin1 Shank2 Crkl Wasf1 Kirrel3 Asic2 Cux2 Lrp8 Erc2 Gnpat Cacnb1 Cbln2 Tnr Ube2v2 Wasf3 Dnm3 Anapc2 L1cam Grid2 Septin11 Cbln1 Pcdhb19 Ryk App Lrrtm2 Zdhhc2 Ache Dbn1 Ctnna2 Bdnf Syn1 Tmem108 Slitrk3 Pcdhb5 Wnt7a Vcp Lrfn1 Shank3 Zdhhc15 Amigo2 Pcdhb6l Tiam1 Pcdhb9 Pcdh17 Prnp Sez6 Arc Tanc2 Farp1 Dock10 Adgrl3 Syndig1 Ptprd Ppfia4 Pten Nefh Lhfpl4 Ywhaz Disc1 Afg3l2 Bhlhb9 Gpc4 | 9 |
| GOCC ENDOPLASMIC RETICULUM LUMEN | 5.38E-17 | 2.62E-14 | 1.07E+00 | -0.557 | -2.305 | 255 | Pdgfd Lgals1 Col4a5 P4hb Col4a1 Fmo1 Col14a1 Cp Fstl1 Col1a1 Txndc5 Col6a3 Casq1 Crtapl1 Sumf2 Col4a6 Ppib Col26a1 Col12a1 Bmp15 P3h1 Lamb1 Col7a1 Col18a1 Fbn1 Col3a1 Lamc1 Col13a1 Timp1 Calu Cd4 Col4a4 Fn1 Ccn1 P4ha2 Csf1 Igfbp7 Arsg Penk C4b Rdh5 Tspan15 Apol3 Fkbp10 Pdia5 Col4a2 Col9a2 Col6a2 Cnpy3 Fkbp14 Ltbp1 Hrc Ces1d Cercam Erap1 Ctsc Col24a1 Lamb2 Dmp1 Casq2 Col5a3 Poglut3 Vwa1 Mxra8 Bche Gpx7 B2m Adamts7 Fstl3 Sts Wnt5a Tnc Col9a1 Notum Serpinh1 Col5a1 Rnaset2 Colgalt1 Wnt6 Dbi Sdf2l1 Cln6 Plod3 Selenom Sumf1 Serpind1 Col6a1 Col2a1 Wnt5b Poglut2 Col4a3 Mfge8 Ganab C3 Wnt3 Tor2a Adamtsl4 | 10 |
| GOCC BASEMENT MEMBRANE | 5.88E-17 | 2.62E-14 | 1.06E+00 | -0.739 | -2.646 | 85 | Ptn Col4a5 Nid1 Anxa2 Lama1 Col4a1 Itgb4 Tgfbi Mmrn2 Lama4 Adamts1 Col4a6 Lamb1 Col7a1 Col18a1 Loxl2 Fbn1 Egflam Lamc1 Colq Timp1 Loxl1 Col4a4 Fn1 Nid2 Lama5 Col4a2 Sparc Hmcn1 Serpinf1 Lamb2 Vwa1 Lama2 Lamc3 Tnc Col9a1 Col5a1 | 11 |
| GOCC POSTSYNAPTIC MEMBRANE | 1.52E-16 | 6.22E-14 | 1.05E+00 | 0.55 | 2.33 | 246 | Slitrk2 Adcy1 Lrrc4c Gabrg2 Kcnb1 Cdh10 Chrna5 Ptprt Shisa7 Grin2a Gabrb2 Grm5 Adora1 Gabrb1 Lrfn5 Sorcs3 Gabra1 Shank1 Slc8a1 Gabrb3 Epha7 Cacng3 Grid1 Grin2b Nlgn1 Slitrk1 Gria2 Lrfn4 Grm3 Ddn Gabra5 Gabra2 Lrrtm3 Ank1 Kcnc3 Grik3 Gphn Gabrg1 Gabrg3 Erbb4 Grm7 Grik2 Gabra3 Lrrc4 Mx2 Il1rapl1 Gria1 Dgkb Atp2b2 Chrm1 Adam22 Nlgn2 Glrb Clstn1 Adra1a Shisa8 Adgrl2 Dagla Gria3 Chrm3 Gabbr1 Cdh9 Dlg2 Epha4 Lrfn3 Rgs7bp Dlg4 Nlgn3 Dnm1 Cacng8 Kcna3 Dnaja3 Kcna1 Lin7c Shisa6 Nptn Htr2a Arrb1 Gria4 Pcdh8 Dcc Grip1 Magee1 Picalm Grik1 Gabrq Plppr4 Chrna2 Cnih2 Shank2 Crkl | 12 |
| GOCC NEURON SPINE | 4.31E-15 | 1.62E-12 | 9.97E-01 | 0.593 | 2.403 | 171 | Homer1 Dgki Abi2 Camk2a Ppp1r9a Ppp3ca Atp2b1 Frmpd4 Shisa7 Grin2a Adora1 Shank1 Cd3e Slc8a1 Ntsr1 Ophn1 Asap1 Nlgn1 Gria2 Grm3 Apba2 Ddn Ptk2 Kcnc3 Anks1b Arhgap44 Lrrc4 Mx2 Gria1 Gpm6a Slc1a1 Slc8a2 Sipa1l1 Clstn1 Shisa8 Itpka Dagla Gria3 Ppp1r9b Lzts3 Epha4 Dlg4 Cyfip1 Dnm1 Akap5 Abr Hpca Shisa6 Rgs14 Cttnbp2 Cdk5r1 Ptk2b Arrb1 Gria4 Ppfia2 Itsn1 Kcna4 Cpeb4 Kcnn2 Arhgap33 Cnih2 Shank2 Asic2 Apba1 Rph3a Grin1 Dnm3 Grid2 Septin11 Actn2 App Psmc2 Psd Ppp1r1b Shank3 Tiam1 Tenm2 Oprm1 Ppp1ca Sez6 Arc Tanc2 Arhgap32 Farp1 Dock10 Syndig1 Calb1 Pten Rac1 Mpp2 Oprd1 Strn | 13 |
| GOBP BEHAVIOR | 2.02E-14 | 7.06E-12 | 9.76E-01 | 0.442 | 1.989 | 468 | Adcy1 Kalrn Homer1 Ext1 Nr4a3 Gabrg2 Actr2 Slc12a5 Tmod2 Chrna5 Jph3 Gad1 Ncoa2 Lrrk2 Shisa7 Grin2a Nrxn1 Grm5 Syngap1 Sgip1 Htr1a Cntnap4 Adora1 Brinp1 Sorcs3 Rnf180 Pja2 Npy5r Ttbk1 Shank1 Usp46 Btbd9 Ntsr1 Adcy8 Grid1 Grin2b Eif4e Sobp Nlgn1 Meis2 Slitrk1 Cic Id2 Gnao1 Ncoa1 Grp Pak5 Gabra5 Vps13a Mapk1 Sptbn2 Ptprz1 Grik2 Git1 Mbd5 Slitrk5 Gria1 Tafa2 Htr1b Prkce Dab1 Slc1a1 Slc8a2 Ncor1 Adcyap1 Nova1 Mecp2 Adam22 Nlgn2 Thrb Oxr1 Adgrb3 Glrb Fgf12 Eif2ak4 Plcb1 Sgk1 Nr4a2 Pias1 Pde1b Ppp1r9b Fezf2 Klk8 Rasgrf1 Adcy3 Mapk8ip2 Epha4 Negr1 Dlg4 Homer2 Pafah1b1 Nlgn3 Tbr1 Ubr3 Kcnj10 Egfr Ube2q1 Hcrt Prex2 Prkar1b Nptn Rgs14 Arrdc3 RGD1562339 Htr2a Atxn1 Lilrb3 Neurod2 Picalm Strbp Ankh Npas2 Ddhd2 Hcrtr1 Mme Abat Slc4a10 Ythdf1 Shank2 Cnr1 Kirrel3 Nqo2 Cux2 Brsk1 Plk2 Egr1 Apba1 Scn2a Gfral Abl2 Slc11a2 Ppp3cb Tnr Mchr1 Gpr37 Nf1 Grin1 Clcn3 Rps6kb1 Neto1 Mtor Ntf3 | 14 |
| GOCC INTRINSIC COMPONENT OF SYNAPTIC MEMBRANE | 4.61E-14 | 1.51E-11 | 9.65E-01 | 0.629 | 2.463 | 132 | Slitrk2 Adcy1 Lrrc4c Cdh10 Atp2b1 Ptprt Grin2a Gabrb2 Adora1 Lrfn5 Sorcs3 Slc1a2 Slc8a1 Epha7 Adcy8 Cacng3 Grid1 Nlgn1 Slitrk1 Lrfn4 Grm3 Gabra5 Kcnj3 Lrrtm3 Gphn Gabrg3 Erbb4 Gabra3 Lrrc4 Gria1 Htr1b Gpm6a Chrm1 Adam22 Nlgn2 Clstn1 Adra1a Adgrl2 Dagla Cdh9 Epha4 Lrfn3 Rgs7bp Nlgn3 Ntng2 Kcna3 Kcna1 Shisa6 Nptn Htr2a Pcdh8 Dcc Plppr4 Ptpra Cnih2 Cnr1 Adam23 | 15 |
| GOBP SYNAPSE ASSEMBLY | 2.29E-13 | 6.66E-11 | 9.33E-01 | 0.582 | 2.338 | 160 | Slitrk2 Gabrg2 Ppp1r9a Nrxn1 Gabrb2 Add2 Lrfn5 Lrrn3 Gabra1 Shank1 Gabrb3 Vps35 Epha7 Nlgn1 Slitrk1 Lrfn4 Gabra2 Lrrtm3 Pcdhb16l Lrp4 Sptbn2 Erbb4 Slitrk5 Flrt2 Lrrc4 Il1rapl1 Gpm6a Mecp2 Pcdhb2 Nlgn2 Adgrb3 Clstn1 Lgi2 Adgrl2 Lrrn1 Nrcam Cdh9 Lrfn3 Pcdhb21 Amigo1 Nlgn3 Ntng2 Nptn Cbln4 Robo2 Setd5 Shank2 Crkl Kirrel3 Asic2 Cux2 Gnpat Cbln2 Ube2v2 Dnm3 Grid2 Cbln1 Pcdhb19 Ryk App Lrrtm2 Ache Bdnf Slitrk3 Pcdhb5 Wnt7a Lrfn1 Shank3 Amigo2 Pcdhb6l Pcdhb9 Pcdh17 Farp1 Adgrl3 Syndig1 Ptprd Pten Lhfpl4 Bhlhb9 Gpc4 Pcdhb20 | 16 |
| GOCC CATION CHANNEL COMPLEX | 2.31E-13 | 6.66E-11 | 9.33E-01 | 0.553 | 2.277 | 193 | Trpc5 Kcnab2 Kcnb1 Kcnc4 Kcns3 Kcnq2 Shisa7 Grin2a Kcnq3 Cacnb4 Scn3b Micu3 Shank1 Scn1a Cacng3 Grin2b Cacnb2 Nlgn1 Gria2 Kcnj3 Pde4d Kcnc3 Grik3 Trpc4 Trpc6 Grik2 Kcnh2 Vwc2l Scn2b Gria1 Hcn1 Olfm3 Cacna1i Kcnj6 Sestd1 Cacna2d1 Scn1b Calm2 Shisa8 Dpp10 Trpc1 Gria3 Dlg2 Kcnab1 Dlg4 Cacna2d3 Kcnn1 Amigo1 Cacng8 Kcna3 Trpv6 Ccdc51 Kcna1 Fkbp1a Shisa6 Akap6 Fkbp1b Nos1ap Ptk2b Gria4 Kcng1 Kcna4 Kcnh4 Kcna2 Kcnj11 Cnih2 Ptpa Kcnmb2 Kcns2 Scn2a Cacnb1 Scn8a Kcnab3 Trpc7 Dpp6 Grin1 Kcnh1 Calm3 Micu1 Cacna1h Ryr1 Kcnma1 Cacng7 | 17 |
| GOBP CELL SUBSTRATE ADHESION | 4.31E-13 | 1.17E-10 | 9.33E-01 | -0.484 | -2.052 | 323 | Postn Jag1 Apod Ptn Lgals1 Nid1 P4hb Vamp3 Itgb4 Vcl Acvrl1 Col1a1 Foxf1 Rhoa Itga5 Unc13d Col26a1 Itgal Fbln5 Id1 S100a10 Vtn Lamb1 Smad6 Angpt2 Sned1 Col3a1 Egflam Lamc1 Fermt3 Emilin1 Col13a1 Vwf Adamts12 Bcam Kif14 Pdpn Flna Fn1 Ccn1 Cd63 Bcl2l11 Dnm2 Nid2 Lama5 Alox15 Fzd4 Csf1 Wdpcp Arl2 Itgb2 Itga2b Myoc Tesk2 Zyx Cib1 Efna1 Lims2 Lamb2 Dock5 Dmp1 Iqgap1 Parvg Npnt Col5a3 Sfrp1 Ccn2 Fam107a Src Ppard Lamc3 Bves Phldb2 Notch1 Itga1 Sdc4 Trip6 Ddr1 Timm10b Kdr Fat2 Cass4 Dusp22 Cd44 Fbln1 Utrn Rras Itgb1 Akip1 Sorbs3 Axl Tecta Ccdc80 Pkd1 | 18 |
| GOBP DENDRITE DEVELOPMENT | 4.79E-13 | 1.23E-10 | 9.21E-01 | 0.533 | 2.228 | 220 | Trpc5 Kalrn Abi2 Camk2a Actr2 Ppp1r9a Ppp3ca Slc12a5 Lrrk2 Dhx36 Syngap1 Pak3 Wasl Dnm1l Opa1 Shank1 Cd3e Kif1a Hecw2 Asap1 Rtn4ip1 Nlgn1 Mef2a Btbd3 Lrp4 Trpc6 Pacsin1 Nck2 Arhgap44 Slitrk5 Il1rapl1 Bcl11a Cdkl5 Dgkg Dab1 Mecp2 Dclk1 Tnik Ube3a Sipa1l1 Adgrb3 Hecw1 Itpka Ppp1r9b Lzts3 Fezf2 Tmem106b Mapk8ip2 Epha4 Dlg4 Flrt1 Cyfip1 Ctnnd2 Camsap2 Kidins220 Prex2 Sarm1 Cdk5r1 RGD1307443 Ppfia2 Itsn1 Dcc Grip1 Picalm Acsl4 Zfp365 Crk Camk2b Arhgap33 Nsmf Mink1 Arf1 Rapgef2 Srcin1 Shank2 Prex1 Crkl Gsk3b Cux2 Lrp8 Plk2 | 19 |
| GOCC POSTSYNAPTIC SPECIALIZATION MEMBRANE | 5.37E-13 | 1.30E-10 | 9.21E-01 | 0.655 | 2.464 | 100 | Adcy1 Lrrc4c Cdh10 Ptprt Shisa7 Grin2a Gabrb2 Grm5 Lrfn5 Sorcs3 Epha7 Cacng3 Grid1 Grin2b Nlgn1 Slitrk1 Gria2 Lrfn4 Gabra5 Lrrtm3 Gphn Erbb4 Gabra3 Lrrc4 Gria1 Atp2b2 Chrm1 Adam22 Nlgn2 Clstn1 Dagla Dlg2 Epha4 Lrfn3 Rgs7bp Dlg4 Cacng8 Lin7c Shisa6 Nptn Dcc Plppr4 Cnih2 | 20 |
| GOBP SENSORY PERCEPTION OF LIGHT STIMULUS | 5.58E-13 | 1.30E-10 | 9.21E-01 | -0.577 | -2.282 | 171 | Nyx Rs1 Slc24a1 Trpm1 Tgfbi Impg2 Ppef2 Cnga1 Cryaa Grk1 Cyp1b1 Col1a1 Cngb1 Pde6g Lum Rom1 Cplx4 Gnat2 Arr3 Cplx3 Guca1a Col18a1 Cacna2d4 Dll4 Crybg3 Adgrv1 Cacna1f Rgs9 Gucy2d Lrit3 Rbp3 Timp3 Impg1 Crb1 Tulp1 Rdh5 Rpgrip1 Cabp1 Prcd Zic2 Rax Rp1l1 Hmcn1 Aipl1 Rlbp1 Pde6c Clic5 Lamb2 Unc119 Clrn1 Guca1b Lamc3 Cngb3 Crx Opn1sw Rgs16 Rd3 Arl6 Crygs Cln6 Pdc Fam161a Pde6d Cnnm4 Slitrk6 Pde6a Col2a1 Rbp4 Hps1 Gnat1 | 21 |
| GOCC PRESYNAPSE | 8.53E-13 | 1.90E-10 | 9.21E-01 | 0.427 | 1.92 | 463 | Prkcb Dgki Rab8b Cdh10 Atp2b1 Synj2 Gad1 Lrrk2 Doc2b Grin2a Nrxn1 Vamp7 Stx1b Cntnap4 Adora1 Adgrl1 Dnm1l Napa Atcay Slc1a2 Vamp4 Slc8a1 Kif1a Ntsr1 Ophn1 Rab10 Adcy8 Gucy1b1 Cacnb2 Nlgn1 Sv2c Grm3 Apba2 Gabra5 Gabra2 Kcnj3 Kcnc3 Nts Grik3 Septin6 Pacsin1 Sptbn2 Erbb4 Grm7 Grik2 Unc13a Git1 Cadps2 Arhgap44 Ston2 Mx2 Gad2 Gria1 Htr1b Gpm6a Atp2b2 Slc1a1 Slc8a2 Chrm1 Rab5al1 Adcyap1 Tnik Stx6 Septin3 Gnb5 Nlgn2 Dnajc6 Calm2 Adra1a Vamp1 Dnajc5 Gabbr1 Cdh9 Epha4 Lrfn3 Btbd8 Ndel1 Rgs7bp Dlg4 Cyfip1 Nlgn3 Dnm1 Ntng2 Kcna3 Kcnj10 Cadps Stx12 Hcrt Mctp1 Kcna1 Lin7c Nptn Cttnbp2 Cdk5r1 Htr2a Ppfia2 Cplx2 Pcdh8 Itsn1 Snph Zdhhc17 Exoc3 Cops5 Picalm Kcna2 Gak Mme Sv2b Slc4a10 Pi4k2a Sncaip Srcin1 Flot1 Sv2a Cnr1 Kirrel3 Adam23 Brsk1 Erc2 Syt2 Aak1 Clcn4 Apba1 Cyp46a1 Scn2a Rph3a Slc6a5 Slc4a8 Rnf40 Sypl2 Grin1 Syt1 Kcnh1 Clcn3 Calm3 Ntf3 Syt10 Dnm3 Hcn3 Fzd3 Stxbp5 Cntn6 Iqsec1 Trim9 App Cbarp Synpr Rims1 Svop Bdnf Syn1 Efnb3 Syngr1 Wnt7a Syn3 Phaf1 Oprm1 Pcdh17 Ppp1ca Calm1 Rab6b Ncs1 Syndig1 Ptprd Dnmbp Ppfia4 Calb1 Synj1 Disc1 Oprd1 Rab35 Slc6a17 Gpc4 Pde2a Dmxl2 Syt7 Rab40b Sncb Syn2 Prkcg Syt11 Septin8 Cask Rab11b Slc17a5 Ppfia3 Rab2b Lrrc4b Cntnap1 Htt Fcho2 Wdr7 Doc2a | 22 |
| GOCC INTRINSIC COMPONENT OF POSTSYNAPTIC MEMBRANE | 9.98E-13 | 2.13E-10 | 9.10E-01 | 0.665 | 2.492 | 97 | Slitrk2 Adcy1 Lrrc4c Cdh10 Ptprt Grin2a Gabrb2 Adora1 Lrfn5 Sorcs3 Slc8a1 Epha7 Cacng3 Grid1 Nlgn1 Slitrk1 Lrfn4 Grm3 Gabra5 Lrrtm3 Erbb4 Gabra3 Lrrc4 Gria1 Chrm1 Adam22 Nlgn2 Clstn1 Adra1a Adgrl2 Dagla Cdh9 Epha4 Lrfn3 Rgs7bp Nlgn3 Kcna3 Kcna1 Shisa6 Nptn Htr2a Pcdh8 Dcc Plppr4 Cnih2 | 23 |
| GOMF GLYCOSAMINOGLYCAN BINDING | 2.02E-12 | 4.13E-10 | 8.99E-01 | -0.565 | -2.229 | 169 | Postn Ptn Clec3b Epyc Pcolce Tgfbr3 Impg2 Anxa6 Ccn4 Fstl1 Adamts1 Bgn Vtn Stab1 Fbn1 Egflam Col13a1 Colq Adamts15 PCOLCE2 Fn1 Ccn1 Eng Prelp Impg1 Hapln2 Crispld2 Cxcl10 Layn Pgf Fgf1 Col5a3 Sfrp1 Ccn2 Dcn Eva1c Ltbp4 Col5a1 Fgfr4 Tlr2 Cfh Thbs2 Hyal2 Cd44 Cxcl13 Cemip Serpind1 Ccn5 Ccdc80 Nod1 | 24 |
| GOCC INTRINSIC COMPONENT OF POSTSYNAPTIC SPECIALIZATION MEMBRANE | 2.16E-12 | 4.23E-10 | 8.99E-01 | 0.733 | 2.543 | 62 | Adcy1 Lrrc4c Cdh10 Ptprt Grin2a Gabrb2 Lrfn5 Sorcs3 Epha7 Cacng3 Grid1 Nlgn1 Slitrk1 Lrfn4 Gabra5 Lrrtm3 Erbb4 Gabra3 Lrrc4 Gria1 Chrm1 Adam22 Nlgn2 Clstn1 Lrfn3 Rgs7bp Shisa6 Nptn Dcc Plppr4 Cnih2 | 25 |
