## Supplemental S1-S4 for "Comparative gene pathway analysis during adolescent binge-EtOH exposure, withdrawal, and following abstinence": Table_S4.docx

**Table S4A. Top 25 Pathways from GSEA (based on p-value) using Gene Sets of MSigDB ImmuneSigDB for PND 70**

| Pathway | pval | padj | log2err | ES | NES | size | Genes | Rank |
| --- | --- | --- | --- | --- | --- | --- | --- | --- |
| GSE42021 TREG PLN VS TREG PRECURSORS THYMUS DN | 2.14E-09 | 1.05E-05 | 7.75E-01 | 0.507 | 2.112 | 171 | Zc3hav1 Irf1 Pml RT1-M3-1 Ifitm3 Dusp22 Ifit3 Ifi44 Irf9 Blzf1 RT1-CE5 Ifit2 Stat2 Gtf2a1 Rasgrp3 B2m Dsp Tnfsf10 Ifi35 Plscr1 Ube2l6 Trafd1 Heg1 Tle3 Sqor Jade2 Tap1 Sp110 Lypd3 Slc25a28 Dcp1a Nrp1 Bst2 Tlr3 Tapbp Elf1 RT1-CE7 RT1-CE4 Lgals3bp Samhd1 Parva Gna14 Cd3e Bcl2l13 Decr2 Serpinb9 Oas2 Isg20 Tmem140 St3gal4 Smarca5 Ogfr Ifitm2 Nfe2l3 Adgre1 Dedd Eif2ak2 Tent5a Bmpr2 Oasl Prkd2 Oas1a | 1 |
| GSE37533 PPARG2 FOXP3 VS FOXP3 TRANSDUCED CD4 TCELL DN | 4.08E-08 | 5.78E-05 | 7.20E-01 | 0.48 | 2.003 | 176 | Irf1 Pml Gtf2b RT1-M3-1 Ifit3 Ifi44 Irf9 Blzf1 Slfn4 Ripk1 RT1-CE5 Ifit2 Stat2 Ifi35 Plscr1 Ube2l6 Trafd1 Uba7 Pla1a Fhl3 Sqor Jade2 Tap1 Sp110 Slc25a28 Vezf1 Cyp2j4 Tlr3 Pcf11 Elf1 RT1-CE7 RT1-CE4 Samhd1 Dhx58 Rtp4 Nf1 Bcl2l13 Fbxl7 Oas2 Isg20 Gsdmd Ogfr Stard5 Jak2 Eif2ak2 Tent5a Oasl Prkd2 Oas1a Zfyve26 Msx1 Kansl1l Spats2l Csf1 Fes Napa Cnp Siah2 Trim21 Wars1 N4bp1 Ceacam1 Themis2 Tymp Psmb9 RT1-CE10 Ddx58 Nmi Il15ra Plekha4 Art3 Sp100 Rnf19b Irs1 Tdrd7 Pdzd2 Ly6e Usp25 Apol9a Rps18 | 2 |
| GSE42021 CD24HI VS CD24INT TREG THYMUS DN | 4.62E-08 | 5.78E-05 | 7.20E-01 | 0.474 | 1.978 | 173 | Erap1 Irf1 Ctsl Pml RT1-M3-1 Ifitm3 Ifit3 Ifi44 Irf9 RT1-CE5 Ifit2 Rasgrp3 B2m Tapbpl Tnfsf10 Psmb3 Ifi35 Plscr1 Zfp36 Ube2l6 Trafd1 Cd38 Uba7 Sqor Zfp710 Jade2 Hist1h2bo Tap1 Sp110 Slc25a28 Bst2 Tlr3 Tapbp RT1-CE7 Tasor2 Parp3 RT1-CE4 Lgals3bp Samhd1 Serping1 Max Rtp4 Il6st Bcl2l13 Plaur Serpinb9 Cfh Oas2 Isg20 | 3 |
| GSE36527 CD69 NEG VS POS TREG CD62L LOS KLRG1 NEG UP | 4.75E-08 | 5.78E-05 | 7.20E-01 | 0.472 | 1.977 | 180 | Crybg1 Cdc42se2 Irf1 Pml Ifit3 Ifi44 Irf9 Slfn4 Ripk1 Ifit2 Stat2 Socs3 Myof Cmtm6 Gbp6 Notch1 Tnfsf10 Tex2 Rad51ap1 Ifi35 Tor3a Trafd1 Tor1aip2 Pttg1 Tap1 Lysmd2 Lipa Irgm Tor1aip1 Exo1 Map3k8 Casp4 Gadd45g Tapbp Tasor2 Lyzl4 Lgals3bp Znfx1 Samhd1 Dhx58 Pex26 Rtp4 Ttc39b Asb3 Fnbp4 Morc3 Gbp4 Oas2 Isg20 Tmem140 Arhgef10 Ogfr Frmd4b Pnpt1 Slfn13 | 4 |
| GSE18791 CTRL VS NEWCASTLE VIRUS DC 8H DN | 9.03E-08 | 8.80E-05 | 7.05E-01 | 0.489 | 2.009 | 156 | Irf1 Pml Ifitm3 Ifit3 Ifi44 Rnf213 Ifit2 Stat2 Siglec1 Ints12 Tnfsf10 Agap2 Pi4k2b Slfn5 Plscr1 Ube2l6 Hk2 Grhl1 Dusp5 Ddit4 Tap1 Sp110 Cdh11 Sema4d Bcl6b Pelo Lgals3bp Znfx1 Dhx58 Serping1 Nlrc5 Amotl2 Rtp4 Gbp4 Oas2 Isg20 Cnih2 Nppa Tmem140 Ptk2b Gadd45b Pnpt1 Dtx3l Stard5 Tent5a Fam135a Oasl | 5 |
| GSE21546 WT VS SAP1A KO DP THYMOCYTES UP | 2.29E-07 | 1.86E-04 | 6.90E-01 | 0.476 | 1.97 | 163 | Zc3hav1 Pml Serpine1 RT1-M3-1 Ifitm3 Ifit3 Ifi44 Irf9 RT1-CE5 Rnf213 Ifit2 Stat2 Tjp1 B2m Gsg1 Tapbpl Ifi35 Plscr1 Ube2l6 Heg1 Agtrap Trim69 Tap1 Sp110 Slc25a28 Vezf1 Gucd1 Bst2 RT1-CE7 RT1-CE4 Lgals3bp Znfx1 Samhd1 Dhx58 Bdh2 Nlrc5 Bcl2l13 Baz1a Oas2 Cdk5rap1 Tmem140 Mt1 Ogfr Ifitm2 Pnpt1 Dtx3l Angpt2 Eif2ak2 Tent5a Adgrg6 Oasl Prkd2 Psmg2 Wdr25 Oas1a Pfkfb3 Msx1 Sox9 | 6 |
| GSE42724 NAIVE BCELL VS PLASMABLAST UP | 2.70E-07 | 1.88E-04 | 6.75E-01 | 0.466 | 1.943 | 172 | Irf1 Pml Ifitm3 Ifit3 Ifi44 Irf9 Ripk1 RT1-CE5 Ifit2 Stat2 Rab8a Siglec1 Tapbpl Casp12 Tnfsf10 Ifi35 Plscr1 Ube2l6 Trafd1 Parp10 Cd38 Uba7 Mvp Arpin Tap1 Sp110 Slc25a28 Bst2 Tlr3 Tuft1 Lgals3bp Znfx1 Tyw5 Dhx58 Serping1 Nlrc5 Rtp4 Shisa5 Axl Gbp4 Oas2 Isg20 Tmem140 Ifitm2 Stom Pnpt1 Dtx3l Stard4 Acsm5 Eif2ak2 Ephb2 Tent5a Oasl Oas1a | 7 |
| GSE37533 PPARG1 FOXP3 VS FOXP3 TRANSDUCED CD4 TCELL DN | 6.37E-07 | 3.88E-04 | 6.59E-01 | 0.458 | 1.917 | 178 | Zc3hav1 Irf1 Pml Gtf2b RT1-M3-1 Ifitm3 Ifit3 Ifi44 Irf9 Nfyb Slfn4 Ripk1 RT1-CE5 Ifit2 Stat2 Lrrc32 Tnfsf10 Ifi35 Plscr1 Ube2l6 Trafd1 Uba7 Pla1a Fhl3 Sqor Jade2 Tap1 Sp110 Slc25a28 Vezf1 Bst2 Tlr3 Elf1 RT1-CE7 RT1-CE4 Samhd1 Dhx58 Max Rtp4 Bcl2l13 Fbxl7 Baz1a Oas2 Isg20 Gsdmd Tmem140 Ogfr Ifitm2 Dedd Jak2 Eif2ak2 Tent5a Lgmn Oasl Prkd2 Oas1a Zfyve26 | 8 |
| GSE22140 GERMFREE VS SPF MOUSE CD4 TCELL UP | 1.00E-06 | 5.43E-04 | 6.44E-01 | 0.451 | 1.881 | 175 | Cd7 Zc3hav1 Gem Irf1 Pml Ifitm3 Ifit3 Ifi44 Irf9 Blzf1 Ifit2 Stat2 Socs3 Furin Tnfsf10 Jmjd6 Psmb3 Ifi35 Zfp36 Ube2l6 Trafd1 Heg1 Mvp Mrc1 P2rx4 Flot1 Stx3 Dusp5 Tap1 Sp110 Ctns Lmo4 C1qb Anks1a Mak16 Klf4 Casp4 Serping1 Smpd1 Naa80 Oas2 Isg20 Il3ra Junb Ifitm2 Pdgfb Gadd45b Oasl Rere Kif2a Oas1a Csf2rb Sft2d2 | 9 |
| GSE18791 UNSTIM VS NEWCATSLE VIRUS DC 10H DN | 1.14E-06 | 5.57E-04 | 6.44E-01 | 0.464 | 1.904 | 153 | Ier2 Zc3hav1 Pml Gtf2b Ifit3 Ifi44 Ifit2 Stat2 Siglec1 Tnfsf10 Pi4k2b Slfn5 Dok7 Plscr1 Stx11 Cd38 Uba7 Csrp2 Jade2 Trim69 Tap1 Sp110 Lysmd2 Map3k8 Casp4 Tlr3 Lgals3bp Znfx1 Dhx58 Serping1 Rtp4 Tmem19 Gbp4 Oas2 Atp10a Isg20 Gadd45b Adgre1 Pnpt1 Dtx3l Stard5 Oasl Hexd Gbp5 Nrip1 Sardh Fas Fgfr3 Cxcl9 Cnp Synpo2 Srgap2 Wars1 Mmp14 | 10 |
| GSE1432 1H VS 24H IFNG MICROGLIA DN | 1.57E-06 | 6.95E-04 | 6.44E-01 | 0.441 | 1.839 | 173 | Crybg1 Erap1 Map3k11 Zfp36l1 Ifitm3 Agpat5 Ifi44 RT1-CE5 Btn2a2 Rfk Guf1 Myof Tapbpl Tnfsf10 RT1-Da Cd38 Mvp Dennd1b Gimap6 Jade2 Tap1 Gimap4 Sp110 Ralb Smurf1 Casp4 RT1-Bb Lgals3bp Pla2g4a Dhx58 Serping1 Max Dipk1a Rad50 Cd74 Prpf39 Baz1a Isg20 C1rl Ifitm2 Stom Fan1 Osbpl11 RT1-Db1 Tma16 Atg3 | 11 |
| GSE41978 ID2 KO VS BIM KO KLRG1 LOW EFFECTOR CD8 TCELL UP | 2.20E-06 | 8.71E-04 | 6.27E-01 | 0.45 | 1.871 | 170 | Zc3hav1 Irf1 Cebpb Ctsl Pml Gtf2b RT1-M3-1 Ifitm3 Hyal1 Trio Ifit3 Irf9 B2m Ifi35 Ccdc68 Plscr1 Zfp36 Ube2l6 Arhgap29 Trafd1 Heg1 Pex11a Sqor Scn9a Tap1 Sp110 Cebpd Slc25a28 Klf4 Bst2 Tlr3 Tapbp RT1-CE7 Tasor2 RT1-CE4 Lgals3bp Serping1 Max Cep63 Met Rtp4 Il1r1 Fnbp4 Cfh Isg20 | 12 |
| GSE34006 A2AR KO VS A2AR AGONIST TREATED TREG UP | 2.33E-06 | 8.71E-04 | 6.27E-01 | 0.437 | 1.844 | 192 | Irf1 Pml RT1-M3-1 Ifit3 Ifi44 Irf9 Slfn4 Ripk1 Ifit2 Stat2 Cmtm6 Gbp6 B2m Notch1 Tnfsf10 Tex2 Rad51ap1 Ifi35 Tor3a Ss18 Trafd1 Tor1aip2 Tap1 Rpl13 Lipa Slc25a28 Irgm Tor1aip1 Bst2 Tapbp RT1-CE7 Tasor2 RT1-CE4 Lgals3bp Znfx1 Samhd1 Dhx58 Dicer1 Pex26 Rtp4 Tmtc4 Il11 Sertad3 Fnbp4 Morc3 Gbp4 Oas2 Isg20 Rgs10 Acrbp Tmem140 Ogfr Pnpt1 Slfn13 Eif2ak2 Noc4l Oasl Cd164 Oas1a Vps54 Pdk1 Dop1b Phlpp1 Atp11b Ppic Nif3l1 Cnp Gbp2 Trim21 Nub1 | 13 |
| GSE37301 MULTIPOTENT PROGENITOR VS GRAN MONO PROGENITOR DN | 3.83E-06 | 1.33E-03 | 6.11E-01 | 0.435 | 1.821 | 181 | S100a4 Crybg1 Irf1 Pml Tab2 RT1-M3-1 Trio Ifit3 Cebpg Adam23 Ripk1 Ifit2 Birc3 Polr2a Rela Hpca Xdh Itpr1 Trafd1 Uba7 Tor1aip2 Qsox1 Tap1 Lysmd2 Pkdcc Nfkb2 Irgm Lcn2 Exo1 Abhd16a Casp4 Gadd45g Tapbp Gja4 Alox12 Lgals3bp Znfx1 Samhd1 Tbk1 Pla2g4a Serping1 Nck1 Il6st Trdmt1 Ddr1 Gbp4 Isg20 Tsr1 Ly75 Ogfr Rad1 Tfdp1 Nr1h2 Jun Dkk3 Eif2ak2 | 14 |
| GSE4748 CTRL VS LPS STIM DC 3H UP | 4.74E-06 | 1.54E-03 | 6.11E-01 | 0.419 | 1.765 | 189 | Elk3 Gtf2b Ccn1 Serpine1 Lifr Ifit3 Fbn1 Slc16a3 Ifi44 Tead3 Ripk1 Sufu Ccn2 Sav1 Bicc1 Polr2a Angptl2 Wasf2 Sesn2 Ifi35 Timp3 Emp1 Adamts1 Ahnak Tll1 Pygl Irgm Cdh11 Ntmt1 Inhba Fzd7 Amfr Iqgap1 Serping1 Zfp532 Amotl2 Nid1 Rtp4 Enc1 Trim2 Stx5 Sertad3 Ghr Arhgef10 Mt1 Rassf8 Ackr3 Pdgfra Nab2 Has2 Dkk3 Col3a1 Prss23 Smad7 Cyp1b1 Col5a2 Cmas Lrrc58 Scara5 Sox9 Pdk1 Rin2 Rhoj Hbegf Lrba | 15 |
| GSE37534 UNTREATED VS PIOGLITAZONE TREATED CD4 TCELL PPARG1 AND FOXP3 TRASDUCED DN | 5.47E-06 | 1.60E-03 | 6.11E-01 | 0.435 | 1.826 | 182 | Atf3 Erap1 Irf1 Pml Ifitm3 Ifit3 Ifi44 Irf9 RT1-CE5 Ifit2 Stat2 Frs2 Anxa2 Sigirr Ifi35 Plscr1 Ube2l6 Cflar Trafd1 Hk2 P2rx4 Jade2 Hist1h2bo Ddit4 Ptn Tap1 Rpl13 Sp110 Slc25a28 Dcp1a Bst2 Tlr3 Id3 Elf1 Rtp4 Acat2l1 Rab20 Cep85 Bcl2l13 Oas2 Atp10a Isg20 Gsdmd Tmem140 Mt1 Ogfr Ifitm2 Gadd45b Stard5 Carhsp1 Eif2ak2 Oasl Prkd2 Oas1a | 16 |
| GSE42021 TREG VS TCONV PLN UP | 5.62E-06 | 1.60E-03 | 6.11E-01 | 0.442 | 1.849 | 177 | Atf3 Irf1 Ctsl Pml Gtf2b RT1-M3-1 Ifitm3 Ifit3 Ifi44 Irf9 RT1-CE5 Ifit2 Stat2 Rasgrp3 B2m Tnfsf10 Ifi35 Ccdc68 Plscr1 Ube2l6 Trafd1 Heg1 Sqor Jade2 Tap1 Sp110 Cebpd Slc25a28 Dcp1a Bst2 Tlr3 Tapbp RT1-CE7 Tasor2 RT1-CE4 Lgals3bp Samhd1 Serping1 Max Rtp4 Bcl2l13 Plaur Nod1 Cfh Oas2 Isg20 Tmem140 Ogfr Ifitm2 Lsm6 Nfe2l3 Jak2 Eif2ak2 Bmpr2 Slc12a7 Oasl Prkd2 Oas1a Msx1 Pclo Spats2l Msrb1 Csf1 Fas Napa Panx1 | 17 |
| GSE19888 ADENOSINE A3R INH VS TCELL MEMBRANES ACT MAST CELL UP | 5.90E-06 | 1.60E-03 | 6.11E-01 | 0.442 | 1.845 | 174 | Zc3hav1 Irf1 Sppl2a Pml Ifit3 Ifi44 Irf9 Rnf213 Egfl6 Ifit2 Stat2 Gbp6 Tnfsf10 Khnyn Myh1 Ifi35 Slfn5 Ube2l6 Uba7 Tor1aip2 Mvp Mgme1 Stx3 Tap1 Sp110 Irgm Casp4 Vwa3b LOC691113 Tlr3 Gpr19 Terf2 Tapbp Ccser1 Hinfp Lto1 Samhd1 Dxo Dhx58 Nlrc5 Rtp4 Dusp18 Zfp287 Shisa5 Serpinb9 Itpka Fam241a Gbp4 Oas2 Tmem243 Myo9b Rhbdd1 Dtx3l Slfn13 Jak2 Ttc21b Ccdc18 Oas1a Steap1 Abra | 18 |
| GSE18791 UNSTIM VS NEWCATSLE VIRUS DC 6H DN | 6.76E-06 | 1.67E-03 | 6.11E-01 | 0.441 | 1.82 | 161 | Zc3hav1 Irf1 Pml Mill1 Gtf2b Ifit3 Ifi44 Irf9 Ripk1 Rnf213 Ifit2 Stat2 Siglec1 Tnfsf10 Agap2 Pi4k2b Plscr1 Trafd1 Uba7 Jade2 Trim69 Tap1 Sp110 Slc25a28 Sema4d Lgals3bp Znfx1 Dhx58 Nlrc5 Amotl2 Rtp4 Oas2 Atp10a Isg20 Elmo2 Stom Pnpt1 Dtx3l Stard5 Eif2ak2 Tent5a Oasl Oas1a Sft2d2 | 19 |
| GSE7218 IGM VS IGG SIGNAL THGOUGH ANTIGEN BCELL DN | 6.86E-06 | 1.67E-03 | 6.11E-01 | 0.473 | 1.893 | 128 | Erap1 Gem Ifitm3 Ifit3 Ifi44 Ifit2 Stat2 Gbp6 Birc3 B2m Tnfsf10 Ifi35 Ube2l6 Adora2a Trafd1 Pttg1 Gimap4 Plat Insrr Filip1l Ptpn6 Lcn2 Bst2 Abhd16a Aldh1a1 Rab19 Tapbp Parp3 Dhx58 Serping1 Nlrc5 Hspb1 Rtp4 Nod1 Serpinb9 Isg20 Gsdmd Ogfr Ifitm2 Rel Upp1 Septin1 Runx2 | 20 |
| GSE21360 NAIVE VS QUATERNARY MEMORY CD8 TCELL DN | 9.31E-06 | 2.16E-03 | 5.93E-01 | 0.433 | 1.805 | 172 | Pml Gtf2b Ifitm3 Dusp22 Ifit3 Ifi44 Irf9 Ripk1 Ifit2 Stat2 Rasgrp3 Ints12 Ifi35 Plscr1 Ube2l6 Trafd1 Cd38 Dennd1b Zzz3 Prrc1 Tap1 Sp110 Timm17a Dcp1a Mak16 Tmem127 Bst2 Casp4 Tlr3 Elf1 Pno1 Cdc73 Dhx58 Esf1 Srd5a1 Rtp4 Klhl20 Serpinb9 Oas2 Isg20 Fam98a Nsun3 Slc22a2 Ifitm2 Mrto4 Cdc14b Eif2ak2 Tent5a Rbm41 Oasl Cul4a Oas1a | 21 |
| GSE34156 NOD2 LIGAND VS TLR1 TLR2 LIGAND 6H TREATED MONOCYTE UP | 1.01E-05 | 2.25E-03 | 5.93E-01 | 0.423 | 1.784 | 192 | Crybg1 Map3k11 Irf1 Pml RT1-M3-1 Ifitm3 Fgf10 Ifit3 Tbpl1 Irf9 Slfn4 Ptpn13 Ifit2 Dusp1 Casp12 Tnfsf10 Mpeg1 Ifi35 Xdh Twist1 Zfp36 Tor3a Apod Ube2l6 Trafd1 Uba7 Tor1aip2 Stx3 Tap1 Pipox Irgm Klf4 Casp4 Rnpepl1 Gadd45g Tapbp RT1-CE7 RT1-CE4 Lgals3bp Znfx1 Samhd1 Il11 Irf5 Shisa5 Serpinb9 Gbp4 Isg20 | 22 |
| GSE19401 NAIVE VS IMMUNIZED MOUSE PLN FOLLICULAR DC UP | 1.12E-05 | 2.36E-03 | 5.93E-01 | 0.424 | 1.775 | 181 | S100a4 Atf3 Zc3hav1 Gem Dnajb1 Serpine1 Ifit3 Slc16a3 Ifi44 Npm3 Atp13a3 Dusp1 Ncor2 Zfp36 Heg1 Btg2 Mrc1 Emp1 P2rx4 Anpep Flot1 Dusp5 Ddit4 Ppp1r15a Plod3 Sp110 Pxdc1 Ppif Eng Lgals1 Phldb1 Crip1 Cd4 Ivns1abp Pdgfa Tubb4a Serpinb9 Il1rap Il3ra Ice1 Junb Cd99 Mxra7 Comt Cd163 Upp1 Il1b Lmna Rps4x Bmpr2 Sec24d Thbs1 Oasl Eif3k Prdm1 | 23 |
| GSE13485 CTRL VS DAY7 YF17D VACCINE PBMC DN | 1.29E-05 | 2.62E-03 | 5.93E-01 | 0.431 | 1.791 | 170 | Slc12a8 Ifitm3 Ifit3 Ifi44 Irf9 Rnf213 Ifit2 Stat2 Adap2 Myof C1galt1c1 Cysltr1 Siglec1 Fancl Trip6 Tnfsf10 Plac8 Rad51ap1 Pi4k2b Ifi35 Plscr1 Ube2l6 Parp10 Heg1 Cd38 Trim69 Tap1 Atp13a1 Sp110 Bst2 Casp4 Cyp2j4 Lgals3bp Dhx58 Serping1 Nid1 Rtp4 Shisa5 Axl Gbp4 Oas2 Atp10a Lilrb3 Ifitm2 Pnpt1 Dtx3l Jak2 Eif2ak2 Oasl Oas1a Ttc21a Grin3a Rin2 Spats2l Mvb12a Fbxo6 | 24 |
| GSE42021 TREG PLN VS CD24INT TREG THYMUS DN | 1.50E-05 | 2.92E-03 | 5.93E-01 | 0.43 | 1.79 | 171 | Erap1 Irf1 Ctsl Pml RT1-M3-1 Ifitm3 Ifit3 Ifi44 Irf9 RT1-CE5 B2m Tapbpl Psmb3 Ifi35 Plscr1 Zfp36 Ube2l6 Trafd1 Heg1 Cd38 Uba7 Gsto1 Sqor Tap1 Sp110 Slc25a28 Klf4 Bst2 Casp4 Tlr3 Tapbp Pelo RT1-CE7 Tasor2 Parp3 RT1-CE4 Lgals3bp Samhd1 Serping1 Max Rtp4 Plaur Cfh Cd74 Oas2 Isg20 | 25 |

**Table S4B. Top 14 Pathways from GSEA (based on p-value) using Gene Sets of MSigDB Hallmark Pathways for PND 70**

| Pathway | pval | padj | log2err | ES | NES | size | Genes | Rank |
| --- | --- | --- | --- | --- | --- | --- | --- | --- |
| HALLMARK EPITHELIAL MESENCHYMAL TRANSITION | 1.85E-12 | 9.23E-11 | 8.99E-01 | 0.533 | 2.237 | 184 | Lamc1 Cald1 Loxl1 Gem Ccn1 Serpine1 Col4a1 Fbn1 Mgp Tnfrsf11b Lama1 Ccn2 Ecm1 Tagln Col16a1 Fbln1 Pdgfrb Fn1 PCOLCE2 Fuca1 Timp3 Acta2 Wipf1 Myl9 Qsox1 Flna Gadd45a Loxl2 Anpep Nid2 Vim Sgcg Plod3 Plod1 Mylk Cdh11 Inhba Bmp1 Sdc4 Oxtr Itga5 Tgfbr3 Bgn Lgals1 Abi3bp Itgb1 Tpm2 Itga2 Sgcb Foxc2 Plaur P3h1 Tpm4 Vegfa Plod2 Col6a3 Lama2 Igfbp2 Fstl3 Sfrp1 Gadd45b Glipr1 Dab2 Col8a2 Jun Col5a3 Col3a1 Col5a1 Col6a2 Thbs1 Col1a1 Notch2 Pcolce Thbs2 Itgb5 Fbln5 Col5a2 Gpc1 Msx1 Col7a1 Fgf2 Fas Ecm2 Eln Emp3 Crlf1 Fbn2 Col4a2 Gas1 Mmp14 Serpinh1 Wnt5a Pmp22 Tgm2 Cd44 Lox Tnc | 1 |
| HALLMARK INTERFERON GAMMA RESPONSE | 2.02E-10 | 5.04E-09 | 8.27E-01 | 0.514 | 2.148 | 178 | Irf1 Sppl2a Pml RT1-M3-1 Ifitm3 Ifit3 Lats2 Ifi44 Irf9 Ripk1 Rnf213 Ifit2 Stat2 Ptpn1 Socs3 Gbp6 B2m Tnfsf10 Ifi35 Plscr1 Ube2l6 Trafd1 Cd38 Mvp RT1-Ba Tap1 Sp110 Lysmd2 Slc25a28 Vamp8 Ptpn6 Bst2 Casp4 Tapbp Il2rb RT1-CE7 RT1-CE4 Lgals3bp Znfx1 Samhd1 Cmklr1 Pla2g4a Dhx58 Serping1 Nlrc5 Rtp4 Irf5 Nod1 Cfh Cd74 Gbp4 Oas2 Isg20 Arl4a Stat1 Ogfr Ifitm2 RT1-Db1 Pnpt1 Upp1 Jak2 Eif2ak2 | 2 |
| HALLMARK INTERFERON ALPHA RESPONSE | 6.87E-09 | 1.14E-07 | 7.62E-01 | 0.588 | 2.235 | 92 | Irf1 Ifitm3 Ifit3 Ifi44 Irf9 Lpar6 Ifit2 Stat2 B2m Ifi35 Plscr1 Ube2l6 Trafd1 Uba7 Tap1 Sp110 Slc25a28 Bst2 Elf1 Lgals3bp Dhx58 Rtp4 Cd74 Gbp4 Isg20 Tmem140 Ogfr Ifitm2 Pnpt1 Eif2ak2 Tent5a Oasl Oas1a Csf1 Mvb12a Cnp Gbp2 Trim21 Nub1 Wars1 Casp8 Ccrl2 Psmb9 RT1-CE10 Nmi Tdrd7 Ly6e Txnip Procr Psme1 | 3 |
| HALLMARK TNFA SIGNALING VIA NFKB | 9.13E-07 | 1.14E-05 | 6.59E-01 | 0.449 | 1.878 | 178 | Klf2 Atf3 Ier2 Gem Irf1 Cebpb Ccn1 Serpine1 Trip10 Ifit2 Socs3 Sik1 Birc3 Dusp1 F3 Rela Tlr2 Zfp36 Cflar Btg2 Gadd45a Dusp5 Ppp1r15a Tap1 Nr4a1 Cebpd Nfkb2 Klf4 Inhba Sdc4 Bhlhe40 Map3k8 Snn Phlda1 Stat5a B4galt1 Sphk1 Tnfrsf9 Il6st Pdlim5 Plaur Plk2 Vegfa Litaf Efna1 Junb Il23a Kdm6b Mxd1 Rel Gadd45b Ackr3 Il1b Jun | 4 |
| HALLMARK OXIDATIVE PHOSPHORYLATION | 7.23E-05 | 7.23E-04 | 5.38E-01 | -0.404 | -1.673 | 193 | Ndufc1 Lrpprc Cox11 Atp5mg Atp5pd Ldha Cox6b1 Ndufab1 Cox7c Mgst3 Oxa1l Atp5me Dlst Prdx3 Atp6ap1 Ndufb4 Mfn2 Timm10 Sdhb Cox6c Glud1 Atp5f1d Cyb5a Uqcrb Ldhb Ndufv2 Tomm70 Atp5f1c Cox5a Iscu Grpel1 Polr2f Ndufa7 Hspa9 Ndufs4 Uqcrc1 Hadhb Idh3a Vdac2 Atp5mc3 Ndufs8 Mrps15 Afg3l2 Hadha Ndufs7 Sdhc Atp5f1a Atp5po Vdac1 Ndufs3 Atp6v0c Ndufb2 Cox6a1 Atp6v1e1 Atp6v1f Mrpl35 Phb2 Tomm22 Rhot1 Timm50 Cs Slc25a11 Slc25a5 Bckdha Uqcrfs1 Timm9 Uqcrq Sucla2 Cox7a2 Sdha Slc25a20 Abcb7 Cox8a Pdp1 | 5 |
| HALLMARK MYC TARGETS V1 | 3.15E-04 | 2.62E-03 | 3.55E-01 | -0.393 | -1.63 | 196 | Rps2 Dek Pcna Hspe1 Ppia Ldha Cad Pwp1 Ndufab1 Mrpl23 Pa2g4 Eif3j Etf1 Tufm Prdx3 Kars Srsf2 Mrps18b Lsm7 Cstf2 Pold2 Rad23b Hnrnpc Tomm70 Eif4e Cox5a Xpo1 Set Nap1l1 Psma7 Fbl C1qbp Eif2s1 Rack1 U2af1 Tra2b Npm1 Acp1 Psmc4 Ilf2 Nolc1 Rps5 Eif4h Aimp2 Dhx15 Eif3b Exosc7 Apex1 Vdac1 Mcm4 Gnl3 Ywhae Rpl18 Ddx21 Pabpc1 Clns1a Phb2 Snrpd2 Ptges3 | 6 |
| HALLMARK APOPTOSIS | 6.45E-04 | 4.61E-03 | 2.66E-01 | 0.396 | 1.618 | 150 | Atf3 Irf1 Ifitm3 Birc3 Faslg Anxa1 Gpx1 Rela Tnfsf10 Pdgfrb Erbb2 Timp3 Cflar Cd38 Btg2 Gsn Ptk2 Emp1 Gadd45a Rara Tap1 Plat Fdxr Casp4 Tgfbr3 Bgn Bid Hspb1 Dap Lgals3 Gstm1 Isg20 Wee1 Dpyd Igf2r Cdkn1b Gadd45b Il1b Jun Lmna Mgmt Smad7 | 7 |
| HALLMARK IL2 STAT5 SIGNALING | 8.24E-04 | 5.15E-03 | 2.36E-01 | 0.374 | 1.563 | 177 | Cdc42se2 Ifitm3 Ecm1 Enpp1 Furin Tnfsf10 Fah Plscr1 Itgae Hk2 Coch Emp1 P2rx4 Gsto1 Scn9a Wls Ahnak Ikzf2 Nrp1 Bhlhe40 Map3k8 Abcb1b Phtf2 Car2 Phlda1 Il2rb Cd48 Ahr Col6a1 Tnfrsf9 Ttc39b Tnfrsf8 Gbp4 Il3ra Twsg1 Arl4a Swap70 St3gal4 Igf2r Slc1a5 Slc39a8 Mxd1 Gadd45b Eef1akmt1 | 8 |
| HALLMARK COAGULATION | 5.12E-03 | 2.84E-02 | 9.27E-02 | 0.397 | 1.533 | 101 | Ctsl Serpine1 Fbn1 Ctsk Anxa1 F3 Furin Pros1 Fn1 Csrp1 Dusp6 Timp3 Hnf4a Gsn Rgn Plat Hpn Bmp1 Mmp9 Serping1 Acox2 Vwf Cd9 Itga2 Ctsh Cfh Tf Pdgfb C1qa Prss23 Lgmn Thbs1 Msrb2 Cfd Adam9 | 9 |
| HALLMARK APICAL SURFACE | 8.20E-03 | 4.10E-02 | 7.24E-02 | 0.518 | 1.646 | 36 | Crybg1 Mdga1 Sulf2 Scube1 Lypd3 Il2rb Ephb4 B4galt1 Hspb1 Plaur Slc2a4 Dcbld2 Srpx Rtn4rl1 | 10 |
| HALLMARK UV RESPONSE DN | 9.13E-03 | 4.15E-02 | 6.99E-02 | 0.354 | 1.437 | 143 | Lamc1 Ccn1 Serpine1 Ltbp1 Mapk14 Tjp1 Dusp1 Anxa2 F3 Ptpn21 Vav2 Pdgfrb Erbb2 Ythdc1 Nrp1 Synj2 Bhlhe40 Tgfbr3 Met Ptgfr Magi2 Cdon Kalrn Pdlim5 Prkca Sipa1l1 Id1 Mt1 Cdkn1b Dab2 Has2 Col3a1 Mgmt Smad7 Col1a1 Notch2 Tfpi Rbpms Fbln5 Col5a2 Atxn1 Apbb2 Ica1 Aggf1 | 11 |
| HALLMARK TGF BETA SIGNALING | 1.82E-02 | 7.60E-02 | 4.86E-02 | 0.442 | 1.516 | 52 | Serpine1 Tjp1 Furin Ncor2 Ppp1r15a Smurf1 Eng Cdh1 Id3 Rhoa Id1 Junb Wwtr1 Bmpr2 Smad7 Thbs1 Smad6 | 12 |
| HALLMARK HYPOXIA | 2.67E-02 | 1.03E-01 | 4.09E-02 | 0.314 | 1.316 | 182 | S100a4 Atf3 Ccn1 Serpine1 Ccn2 Dusp1 Anxa2 F3 Slc37a4 Plac8 Jmjd6 Zfp36 Tgfb3 Hk2 Ak4 Galk1 Csrp2 Ddit4 Ppp1r15a Sdc4 Bhlhe40 Bgn Pfkl P4ha2 Pgm2 Hexa Plaur Cavin1 Prkca Isg20 Vegfa Atp7a Ndst1 Pkp1 Efna1 Mt1 Pdgfb Cdkn1b Slc25a1 Grhpr Ackr3 Srpx Jun Col5a1 Slc2a1 Ilvbl Pfkfb3 Gpc1 Rbpj | 13 |
| HALLMARK ANGIOGENESIS | 3.09E-02 | 1.11E-01 | 3.70E-02 | 0.479 | 1.503 | 34 | S100a4 Fgfr1 Vav2 Ptk2 Pglyrp1 Nrp1 Pdgfa Vegfa Lpl Col3a1 Col5a2 Msx1 | 14 |

**Table S4C. Top 25 Pathways from GSEA (based on p-value) using Gene Sets of MSigDB Canonical Pathways for PND 70**

| Pathway | pval | padj | log2err | ES | NES | size | Genes | Rank |
| --- | --- | --- | --- | --- | --- | --- | --- | --- |
| REACTOME EXTRACELLULAR MATRIX ORGANIZATION | 8.08E-13 | 1.50E-09 | 9.21E-01 | 0.496 | 2.155 | 247 | Lamc1 Itgal Loxl1 Ctsl Serpine1 Col4a1 Col4a5 Ltbp1 F11r Bmp4 Fbn1 Dag1 Ctsk Lama1 Lamb2 Actn1 Loxl3 Furin Col16a1 Ddr2 Fbln1 Col25a1 Fn1 Itga1 P3h2 Itgae PCOLCE2 Itga8 Tgfb3 Scube1 Adamts1 Loxl2 Mfap2 Nid2 Kdr P4hb Col18a1 Plod3 Plod1 Tll1 Bmp1 Sdc4 Icam2 Itga5 Bgn Cdh1 Adamts16 Mmp9 Dmp1 Bmp7 Itgb1 P4ha2 Nid1 Pdgfa Lrp4 Col6a1 Vwf Bsg Itga2 Col13a1 Dmd P3h1 Ddr1 Prkca Pxdn Crtapl1 Plod2 Col6a3 Lamc3 Jam3 Ltbp3 Lama2 Capn6 Itga7 Matn1 Pdgfb Col8a2 Lama5 Scube3 Col5a3 Col3a1 Col5a1 Ctrb1 Col6a2 Emilin2 Cast Thbs1 Col1a1 Col4a3 Pcolce Itga3 Emilin1 Itgb5 Adam9 Fbln5 Itgb4 Col5a2 Col7a1 Col6a6 Capn2 Adamts3 Fgf2 Adamts14 Eln Itgb2 Jam2 Capn10 Fbn2 Col4a2 Mmp14 Serpinh1 Col4a6 Col9a2 Ceacam1 Musk App Itga11 Itgb6 Cd44 Lox Tnc Col9a3 | 1 |
| REACTOME RESPIRATORY ELECTRON TRANSPORT ATP SYNTHESIS BY CHEMIOSMOTIC COUPLING AND HEAT PRODUCTION BY UNCOUPLING PROTEINS | 3.35E-12 | 3.10E-09 | 8.99E-01 | -0.618 | -2.397 | 118 | COX2 ATP8 Ndufc1 CYTB Ndufs5 COX3 ND5 COX1 Lrpprc ND2 ND1 ATP6 Cox11 Atp5mg ND3 Atp5pd Ecsit Cox6b1 Ndufab1 Cox7c Atp5me Ndufb4 Ndufaf2 Sdhb Coq10a Cox6c Atp5f1d Ndufa13 Uqcrb Cox18 Ndufv2 Atp5f1c Cox5a Ndufa7 Ndufs4 Uqcrc1 | 2 |
| WP ELECTRON TRANSPORT CHAIN OXPHOS SYSTEM IN MITOCHONDRIA | 2.91E-11 | 1.79E-08 | 8.63E-01 | -0.654 | -2.45 | 94 | COX2 ATP8 Ndufc1 ND4L CYTB Ndufs5 COX3 ND5 COX1 ND2 ND1 ATP6 Cox11 Atp5mg ND3 Atp5pd Cox6b1 Ndufab1 Cox7c Atp5me Ndufb4 Atp5if1 Sdhb Cox6c Atp5f1d Uqcrb Ndufv2 Atp5f1c Cox5a Ndufa7 Ndufs4 Uqcrc1 | 3 |
| KEGG OXIDATIVE PHOSPHORYLATION | 1.62E-10 | 7.48E-08 | 8.27E-01 | -0.6 | -2.314 | 114 | COX2 ATP8 Ndufc1 ND4L CYTB Ndufs5 COX3 ND5 COX1 ND2 ND1 ATP6 Cox11 Atp5mg ND3 Atp5pd Cox6b1 Ndufab1 Cox7c Atp5me Atp6ap1 Ndufb4 Sdhb Cox6c Atp5f1d Uqcrb Atp6v1c2 Ndufv2 Atp5f1c Cox5a Ndufa7 Ndufs4 Uqcrc1 Atp5mc3 Ndufs8 Ndufs7 Sdhc Atp5f1a Atp5po Ndufs3 Atp6v0c Cox6a2 Ndufb2 Cox6a1 Atp6v1e1 Atp6v1f | 4 |
| REACTOME RESPIRATORY ELECTRON TRANSPORT | 4.98E-10 | 1.84E-07 | 8.01E-01 | -0.617 | -2.322 | 96 | COX2 Ndufc1 CYTB Ndufs5 COX3 ND5 COX1 Lrpprc ND2 ND1 Cox11 ND3 Ecsit Cox6b1 Ndufab1 Cox7c Ndufb4 Ndufaf2 Sdhb Coq10a Cox6c Ndufa13 Uqcrb Cox18 Ndufv2 Cox5a Ndufa7 Ndufs4 Uqcrc1 | 5 |
| KEGG PARKINSONS DISEASE | 1.21E-09 | 3.73E-07 | 7.88E-01 | -0.584 | -2.253 | 114 | COX2 ATP8 Ndufc1 ND4L CYTB Ndufs5 COX3 ND5 COX1 ND2 ND1 ATP6 ND3 Atp5pd Cox6b1 Ndufab1 Cox7c Pink1 Ndufb4 Th Slc18a2 Sdhb Ubb Cox6c Atp5f1d Uqcrb Ndufv2 Atp5f1c Cox5a Ndufa7 Sncaip Ndufs4 Uqcrc1 Ube2j1 Ube2j2 Vdac2 Atp5mc3 Ndufs8 Ndufs7 Sdhc Atp5f1a Atp5po Vdac1 Ndufs3 Cox6a2 Ndufb2 Cox6a1 | 6 |
| REACTOME THE CITRIC ACID TCA CYCLE AND RESPIRATORY ELECTRON TRANSPORT | 1.02E-08 | 2.68E-06 | 7.48E-01 | -0.514 | -2.087 | 165 | COX2 ATP8 Ndufc1 Adhfe1 CYTB Ndufs5 COX3 ND5 COX1 Lrpprc ND2 ND1 ATP6 Cox11 Atp5mg ND3 Atp5pd Ldha Ecsit Cox6b1 Ndufab1 Cox7c Atp5me Dlst Ndufb4 Ndufaf2 Sdhb Coq10a Cox6c Atp5f1d Ndufa13 Uqcrb Ldhb Cox18 Ndufv2 Atp5f1c Cox5a | 7 |
| NABA ECM GLYCOPROTEINS | 1.61E-07 | 3.71E-05 | 6.90E-01 | 0.481 | 1.968 | 151 | Lamc1 Tinagl1 Ccn1 Ltbp1 Fbn1 Mgp Rspo2 Kcp Lama1 Ccn2 Lamb2 Ecm1 Fbln1 Fn1 PCOLCE2 Fndc1 Coch Mfap2 Nid2 Aebp1 Cilp Vwa3b Lrg1 Dmp1 Abi3bp Sbspon Crispld2 Nid1 Ccn3 Vwf Sned1 Egflam Smoc2 Emid1 Pxdn Lamc3 Ltbp3 Lama2 Slit1 Igfbp2 Matn1 Lama5 Srpx Igfbp7 Emilin2 Thbs1 Vwce Pcolce Emilin1 Crispld1 Nell2 Ints14 Thbs2 Fbln5 Ints6l Spon2 Papln Dpt Tnfaip6 Gas6 Lgi4 Ecm2 Eln Fgl2 Vwa5b2 Fbn2 Igsf10 Colq Spon1 | 8 |
| NABA CORE MATRISOME | 2.08E-07 | 4.27E-05 | 6.90E-01 | 0.436 | 1.869 | 220 | Lamc1 Tinagl1 Ccn1 Col4a1 Col4a5 Ltbp1 Fbn1 Mgp Rspo2 Kcp Lama1 Ccn2 Lamb2 Ecm1 Col16a1 Fbln1 Col25a1 Fn1 PCOLCE2 Fndc1 Coch Mfap2 Nid2 Col18a1 Aebp1 Cilp Vwa3b Bgn Lrg1 Dmp1 Abi3bp Sbspon Crispld2 Nid1 Ccn3 Col6a1 Vwf Sned1 Col13a1 Egflam Smoc2 Emid1 Pxdn Col6a3 Lamc3 Ltbp3 Lama2 Slit1 Igfbp2 Matn1 Col8a2 Lama5 Srpx Col5a3 Col3a1 Igfbp7 Col5a1 Col6a2 Emilin2 Thbs1 Col1a1 Col4a3 Vwce Pcolce Emilin1 Crispld1 Nell2 Ints14 Hapln2 Thbs2 Fbln5 Ints6l Col5a2 Spon2 Papln Dpt Tnfaip6 Col7a1 Gas6 Col6a6 Lgi4 Ecm2 Prg4 Hapln3 Eln Fgl2 Vwa5b2 Fbn2 Col4a2 Igsf10 Colq Col4a6 Col9a2 Spon1 | 9 |
| REACTOME TRANSLATION | 4.38E-07 | 7.49E-05 | 6.75E-01 | -0.423 | -1.818 | 269 | Rps2 Rps23 Eif2s3 Mrps17 Mrpl46 Mrps35 Rplp1 Eif3c Mrps7 Cars Hars1 Rpl27a Rps15a Mrpl23 Mrpl41 Oxa1l Eif3j Lars1 Etf1 Tufm Mrps34 Rpl27 Rps13 Rpsa Kars Rpl22l1 Eif2b4 Mrpl19 Cars2 Mrpl32 Mrps18b Mrpl43 Rpl28 Lars2 Rpl15 Sec61a2 Rpl12 Mrpl40 Rps7 Rpl37a-ps1 Rps8 Eif4e Eef1g Nars2 Rps20 Eif3h Rpl29 Tram1 Mrpl33 Eif2s1 mrpl24 Rps12 Rpl10a Mrpl54 Srp68 Ears2 Srp72 Farsb Rpl19 Yars1 Rpl3 Mrps15 Fars2 Tars1 Mrps14 Rps5 Rpl23a Eif4h Fau Rps27 Aimp2 Mrpl57 Aars2 Eif3b Rps3a Rpl5 Rpl36al Gfm2 Mrpl35 Spcs3 Rpl18 Rps15 Pabpc1 Rps11 Mrpl4 Mrps27 Eef2 Eif2b1 Mrpl16 Rpl17 Spcs1 | 10 |
| PID INTEGRIN1 PATHWAY | 4.45E-07 | 7.49E-05 | 6.75E-01 | 0.613 | 2.18 | 62 | Lamc1 Col4a1 Col4a5 Fbn1 Lama1 Lamb2 Fn1 Itga1 Itga8 Col18a1 Itga5 Itgb1 Nid1 Col6a1 Itga2 Plaur Vegfa Col6a3 Lama2 Itga7 Lama5 Col3a1 Col5a1 Col6a2 Thbs1 Col1a1 Col4a3 Itga3 Thbs2 Col5a2 Col7a1 Igsf8 Jam2 Cspg4 Col4a6 Mdk Itga11 Tgm2 Tnc | 11 |
| WP OXIDATIVE PHOSPHORYLATION | 6.57E-07 | 1.01E-04 | 6.59E-01 | -0.644 | -2.199 | 55 | Ndufc1 ND4L Ndufs5 ND5 ND2 ND1 ATP6 Atp5mg ND3 Atp5pd Ndufab1 Atp5me Atp6ap1 Ndufb4 Atp5f1d Ndufv2 Ndufa7 Ndufs4 Atp5mc3 Ndufs8 Ndufs7 Atp5f1a Atp5po Ndufs3 Ndufb2 | 12 |
| REACTOME INTEGRIN CELL SURFACE INTERACTIONS | 2.96E-06 | 4.01E-04 | 6.27E-01 | 0.564 | 2.062 | 73 | Itgal Col4a1 Col4a5 F11r Fbn1 Col16a1 Fn1 Itga1 Itgae Itga8 Kdr Col18a1 Icam2 Itga5 Cdh1 Itgb1 Col6a1 Vwf Bsg Itga2 Col13a1 Col6a3 Jam3 Itga7 Col8a2 Col5a3 Col3a1 Col5a1 Col6a2 Thbs1 Col1a1 Col4a3 Itga3 Itgb5 Col5a2 Col7a1 Col6a6 Itgb2 Jam2 Col4a2 Col4a6 Col9a2 Itga11 Itgb6 Cd44 Tnc Col9a3 | 13 |
| REACTOME EUKARYOTIC TRANSLATION INITIATION | 3.04E-06 | 4.01E-04 | 6.27E-01 | -0.525 | -2 | 105 | Rps2 Rps23 Eif2s3 Rplp1 Eif3c Rpl27a Rps15a Eif3j Rpl27 Rps13 Rpsa Rpl22l1 Eif2b4 Rpl28 Rpl15 Rpl12 Rps7 Rpl37a-ps1 Rps8 Eif4e Rps20 Eif3h Rpl29 Eif2s1 Rps12 Rpl10a Rpl19 Rpl3 Rps5 Rpl23a Eif4h Fau Rps27 Eif3b Rps3a Rpl5 Rpl36al Rpl18 Rps15 Pabpc1 Rps11 Eif2b1 Rpl17 | 14 |
| REACTOME COLLAGEN FORMATION | 3.86E-06 | 4.77E-04 | 6.11E-01 | 0.547 | 2.033 | 80 | Loxl1 Ctsl Col4a1 Col4a5 Loxl3 Col16a1 Col25a1 P3h2 PCOLCE2 Loxl2 P4hb Col18a1 Plod3 Plod1 Tll1 Bmp1 Mmp9 P4ha2 Col6a1 Col13a1 P3h1 Pxdn Crtapl1 Plod2 Col6a3 Col8a2 Col5a3 Col3a1 Col5a1 Col6a2 Col1a1 Col4a3 Pcolce Itgb4 Col5a2 Col7a1 Col6a6 Adamts3 Adamts14 Col4a2 Serpinh1 Col4a6 Col9a2 Lox Col9a3 Ctsb Lama3 | 15 |
| REACTOME SMOOTH MUSCLE CONTRACTION | 9.10E-06 | 1.05E-03 | 5.93E-01 | 0.68 | 2.134 | 34 | Cald1 Pxn Anxa2 Anxa1 Lmod1 Myh11 Itga1 Vcl Acta2 Myl9 Actg2 Mylk Anxa6 Tpm2 Tpm4 Tln1 Myl6l Sorbs3 | 16 |
| WP MITOCHONDRIAL COMPLEX I ASSEMBLY MODEL OXPHOS SYSTEM | 1.44E-05 | 1.57E-03 | 5.93E-01 | -0.616 | -2.071 | 51 | Ndufc1 ND4L Ndufs5 ND5 ND2 ND1 Dmac2 Ecsit Ndufab1 Ndufb4 Ndufaf2 Ndufa13 Ndufv2 Ndufa7 Ndufs4 | 17 |
| REACTOME ELASTIC FIBRE FORMATION | 1.59E-05 | 1.63E-03 | 5.76E-01 | 0.648 | 2.107 | 40 | Loxl1 Ltbp1 Bmp4 Fbn1 Loxl3 Furin Fbln1 Fn1 Itga8 Tgfb3 Loxl2 Mfap2 Itga5 Bmp7 Itgb1 Ltbp3 Emilin2 Emilin1 Itgb5 Fbln5 | 18 |
| WP CYTOPLASMIC RIBOSOMAL PROTEINS | 1.98E-05 | 1.83E-03 | 5.76E-01 | -0.545 | -1.961 | 73 | Rps2 Rps23 Rplp1 Rps6ka6 Rpl27a Rps15a Rps6ka2 Rpl27 Rps13 Rpsa Mrpl19 Rpl28 Rpl15 Rpl12 Rps7 Rpl37a-ps1 Rps8 Rps20 Rpl29 Rps12 Rpl10a Rpl19 Rpl3 Rps5 Rpl23a Fau Rps27 Rps3a Rpl5 Rpl36al Rpl18 Rps15 Rps11 Rpl17 | 19 |
| REACTOME INTERFERON ALPHA BETA SIGNALING | 1.99E-05 | 1.83E-03 | 5.76E-01 | 0.583 | 2.04 | 57 | Irf1 RT1-M3-1 Ifitm3 Ifit3 Irf9 RT1-CE5 Ifit2 Stat2 Ptpn1 Socs3 Ifi35 RT1-M6-2 Ptpn6 Bst2 RT1-CE7 RT1-CE4 Samhd1 Irf5 RT1-M6-1 Oas2 Isg20 | 20 |
| REACTOME LAMININ INTERACTIONS | 2.07E-05 | 1.83E-03 | 5.76E-01 | 0.687 | 2.097 | 30 | Lamc1 Col4a1 Col4a5 Lama1 Lamb2 Itga1 Nid2 Col18a1 Itgb1 Nid1 Itga2 Lamc3 Lama2 Itga7 Lama5 Col4a3 Itga3 Itgb4 Col7a1 Col4a2 Col4a6 | 21 |
| REACTOME ACTIVATION OF THE MRNA UPON BINDING OF THE CAP BINDING COMPLEX AND EIFS AND SUBSEQUENT BINDING TO 43S | 2.99E-05 | 2.45E-03 | 5.76E-01 | -0.59 | -2.006 | 54 | Rps2 Rps23 Eif2s3 Eif3c Rps15a Eif3j Rps13 Rpsa Rps7 Rps8 Eif4e Rps20 Eif3h Eif2s1 Rps12 Rps5 Eif4h Fau Rps27 Eif3b Rps3a Rps15 Pabpc1 Rps11 | 22 |
| WP EBOLA VIRUS PATHWAY ON HOST | 3.04E-05 | 2.45E-03 | 5.76E-01 | 0.474 | 1.857 | 111 | Dab2ip Flnc Ctsl Clec10a RT1-M3-1 RT1-CE5 Socs3 Actn1 Rela Vav2 Itga1 RT1-Da Akt1 RT1-Ba Gsn Flna Cav2 Rhoc Eps15 Nfkb2 Bst2 Icam2 Itga5 RT1-Bb RT1-CE7 RT1-CE4 Itgb1 Iqgap1 Tbk1 Rhoa Itga2 Axl Asgr1 RT1-Db1 Rel Clta Eif2ak2 | 23 |
| REACTOME G ALPHA S SIGNALLING EVENTS | 3.21E-05 | 2.48E-03 | 5.57E-01 | -0.494 | -1.876 | 102 | Gng4 Adcyap1r1 Adcy8 Drd1 Ptger2 Pde7b Pomc Pde8b Calca Adcyap1 Rxfp2 Gng13 Gnaz Pde1b Calcr Cyct Vip Gnb1 Gpr45 Pde4d Pth2r | 24 |
| KEGG NEUROACTIVE LIGAND RECEPTOR INTERACTION | 4.05E-05 | 2.96E-03 | 5.57E-01 | -0.409 | -1.703 | 200 | Adcyap1r1 Chrna7 Drd1 Thrb Ptger2 Gabra3 Adra1a Adra2a Cckbr Gabrq Brs3 Ptafr Ntsr1 Rxfp2 Npy5r Oprl1 Sstr1 Grm8 Hcrtr1 F2r Bdkrb2 Calcr Gabra1 Agtr2 Grm2 Chrm3 Gabrg1 Pth2r Npy1r Npffr1 Oprk1 Grid1 Gabrb2 Chrna5 Tacr3 Trhr Gabrg2 Gabra4 Grik3 P2ry4 Gabre Galr1 Gabra2 Glrb Ptger1 | 25 |

**Table S4D. Top 25 Pathways from GSEA (based on p-value) using Gene Sets of MSigDB GO Annotations for PND 70**

| Pathway | pval | padj | log2err | ES | NES | size | Genes | Rank |
| --- | --- | --- | --- | --- | --- | --- | --- | --- |
| GOBP EXTERNAL ENCAPSULATING STRUCTURE ORGANIZATION | 1.29E-14 | 6.30E-11 | 9.87E-01 | 0.48 | 2.152 | 329 | Lamc1 Itgal Loxl1 Gfod2 Ctsl Sulf2 Ccn1 Serpine1 Col4a1 Col4a5 F11r Fbn1 Dag1 Ctsk Tnfrsf11b Lama1 Egfl6 Ccn2 Lamb2 Loxl3 Notch1 Furin Col16a1 Ddr2 Flrt2 Fbln1 Myh11 Fn1 Itga1 Itgae Foxc1 Itga8 Cflar Sh3pxd2b Scube1 Qsox1 Cav2 Adamts1 Loxl2 Klk4 Mfap2 Flot1 Nid2 Kdr Col18a1 Aebp1 Ercc2 Plod3 Tll1 Hpn Nfkb2 Bmp1 Eng Icam2 Itga5 Gas2 Bgn Cdh1 Adamts16 Mmp9 Dmp1 Phldb1 Itgb1 Meltf Adamts7 Phldb2 B4galt1 Crispld2 Nid1 Pdgfa Col6a1 Nf1 Vwf Bsg Reck Itga2 Col13a1 Foxc2 Ccdc80 Egflam Smoc2 Ddr1 Pxdn Crtapl1 Atp7a Col6a3 Lamc3 Jam3 Ltbp3 Lama2 Adamts10 Adamts20 Itga7 Matn1 Pdgfb Slc39a8 Tcf15 Cst3 Col8a2 Lama5 Scube3 Pdgfra Prdx4 Has2 Col5a3 Col3a1 Col5a1 Ctrb1 Col6a2 Sulf1 Thbs1 Col1a1 Col4a3 Itga3 Emilin1 Hapln2 Itgb5 Olfml2a Fbln5 Cyp1b1 Itgb4 Col5a2 Papln Sox9 Dpt Adamtsl4 Col7a1 Gas6 Capn2 Adamts3 Fgf2 Smpd3 Ecm2 Adamts14 Antxr1 Eln Itgb2 Adamts12 Jam2 Fbn2 Col4a2 Spint2 Mmp14 Colq Serpinh1 Col4a6 Col9a2 Tmem38b Fkbp10 Adamts13 App Itga11 Kif9 Tnfrsf1a Itgb6 Cd44 Lox Tnc | 1 |
| GOCC EXTERNAL ENCAPSULATING STRUCTURE | 2.09E-10 | 5.11E-07 | 8.27E-01 | 0.405 | 1.858 | 424 | S100a4 Wnt4 Lamc1 Loxl1 Tinagl1 Gfod2 Ctsl Ccn1 Serpine1 Col4a1 Col4a5 Fgf10 Ltbp1 Fbn1 Mgp Dag1 Tnfrsf11b Lama1 Egfl6 Bcam Lrrc32 Ccn2 Lamb2 Ecm1 Rarres2 Oc90 Anxa2 Anxa1 F3 Wnt5b Angptl2 Col16a1 Flrt2 Fbln1 Col25a1 Fn1 Plscr1 Egfl7 P3h2 Timp3 Tgfb3 Gdf10 Clec3b Coch Adamts1 Loxl2 Mfap2 Nid2 S100a10 Col18a1 Aebp1 Ptn Plod3 S100a6 C1qb Cilp Anxa6 Tgfbr3 Tlr3 Frem2 Bgn Lgals1 Adamts16 Mmp9 Dmp1 Bmp7 Lingo3 Lrrn2 Abi3bp Lgals3bp Adamts7 Hdgf Serping1 Sbspon Crispld2 Nid1 Ccn3 Anxa5 Otogl Col6a1 Vwf Lrrc24 Cdon Col13a1 Lgals3 Ccdc80 Egflam Serpinb9 Smoc2 Ctsh P3h1 Wnt6 Pxdn Vegfa Col6a3 Lamc3 Ltbp3 Colec12 Nppa Lama2 Adamts10 Adamts20 Mxra7 Matn1 Pdgfb Sfrp1 C1qa Pcsk6 Col8a2 Lama5 Cfp Srpx Col5a3 Col3a1 Igfbp7 Col5a1 Angpt2 Col6a2 Ssc5d Emilin2 Sulf1 Thbs1 Col1a1 Col4a3 Pcolce Emilin1 Lrrn1 Angptl1 C1qc Hapln2 Thbs2 Gpld1 Olfml2a Fbln5 Rtn4rl1 Itgb4 Col5a2 Gpc1 Gpc2 Spon2 Papln Flrt3 Dpt Sema7a Adamtsl4 Col7a1 Col6a6 Erbin Cstb Adamts3 Clu F12 Ecm2 Adamts14 Prg4 Hapln3 Eln Fgl2 Angpt1 Adamts12 Fbn2 Col4a2 Cspg4 Mmp14 Colq Sema3b Lrig3 Serpinh1 Col4a6 Col9a2 Spon1 | 2 |
| GOCC COLLAGEN CONTAINING EXTRACELLULAR MATRIX | 3.78E-10 | 6.18E-07 | 8.14E-01 | 0.44 | 1.964 | 319 | S100a4 Lamc1 Loxl1 Tinagl1 Ctsl Ccn1 Serpine1 Col4a1 Col4a5 Fgf10 Ltbp1 Fbn1 Mgp Dag1 Lama1 Egfl6 Bcam Ccn2 Lamb2 Ecm1 Rarres2 Anxa2 Anxa1 F3 Wnt5b Angptl2 Col16a1 Fbln1 Col25a1 Fn1 Plscr1 Egfl7 P3h2 Timp3 Tgfb3 Gdf10 Clec3b Coch Adamts1 Loxl2 Mfap2 Nid2 S100a10 Col18a1 Aebp1 Ptn Plod3 S100a6 C1qb Cilp Anxa6 Frem2 Bgn Lgals1 Mmp9 Bmp7 Abi3bp Lgals3bp Hdgf Serping1 Sbspon Nid1 Ccn3 Anxa5 Col6a1 Vwf Cdon Col13a1 Lgals3 Ccdc80 Egflam Serpinb9 Smoc2 Ctsh P3h1 Pxdn Col6a3 Lamc3 Ltbp3 Nppa Lama2 Adamts10 Adamts20 Mxra7 Matn1 Pdgfb Sfrp1 C1qa Pcsk6 Col8a2 Lama5 Cfp Srpx Col5a3 Col3a1 Igfbp7 Col5a1 Angpt2 Col6a2 Ssc5d Emilin2 Sulf1 Thbs1 Col1a1 Col4a3 Pcolce Emilin1 Angptl1 C1qc Thbs2 Fbln5 Itgb4 Col5a2 Gpc1 Gpc2 Dpt Sema7a Adamtsl4 Col7a1 Col6a6 Erbin Cstb Adamts3 Clu F12 Prg4 Eln Fgl2 Angpt1 Fbn2 Col4a2 Cspg4 Colq Sema3b Serpinh1 Col4a6 Col9a2 Spon1 | 3 |
| GOBP OXIDATIVE PHOSPHORYLATION | 6.48E-09 | 7.93E-06 | 7.62E-01 | -0.557 | -2.188 | 128 | COX2 ATP8 Ndufc1 ND4L CYTB Ndufs5 COX3 ND5 COX1 ND2 ND1 ATP6 Atp5mg ND3 Atp5pd Cox6b1 Atpsckmt Ndufab1 Actn3 Cox7c Slc25a23 Atp5me Pink1 Ndufb4 Cox6c Atp5f1d Ndufa13 Uqcrb Ndufv2 Atp5f1c Cox5a Iscu Ndufa7 Ndufs4 Uqcrc1 | 4 |
| GOCC MITOCHONDRIAL PROTEIN CONTAINING COMPLEX | 1.77E-08 | 1.73E-05 | 7.34E-01 | -0.458 | -1.947 | 241 | ATP8 Ndufc1 ND4L CYTB Ndufs5 ND5 COX1 ND2 ND1 Mrps17 ATP6 Mrpl46 Dmac2 Atp5mg ND3 Atp5pd Tomm6 Mrps35 Mrps7 Ndufab1 Cox7c Mrpl23 Mrpl41 Atp5me Chchd6 Mcub Mrps34 Ndufb4 Tomm40 Mrpl19 Mrpl32 Timm10 Sdhb Mrps18b Mrpl43 Immp1l Mrpl40 Atp5f1d Ndufa13 Timm23 Clpx Uqcrb Ndufv2 Tomm70 Atp5f1c Cox5a Grpel1 Dbt Ndufa7 Mrpl33 Hspa9 mrpl24 Ndufs4 Uqcrc1 Pam16 Suclg2 Mrpl54 Tomm40l Atp5mc3 Ndufs8 Mrps15 Mrps14 Afg3l2 Bckdhb Micos10 Hadha Ndufs7 Tomm20 Mtx3 Apoo Sdhc Mrpl57 Bckdk Atp5f1a Atp5po Vdac1 Ndufs3 Cox6a2 Ndufb2 Cox6a1 Mrpl35 Mrpl4 Phb2 Malsu1 Mrps27 Tomm22 Mrpl16 | 5 |
| GOCC RESPIRATORY CHAIN COMPLEX | 5.16E-08 | 4.21E-05 | 7.20E-01 | -0.615 | -2.222 | 76 | COX2 Ndufc1 ND4L CYTB Ndufs5 COX3 ND5 COX1 ND2 ND1 Dmac2 ND3 Cox6b1 Ndufab1 Cox7c Ndufb4 Sdhb Ndufa13 Uqcrb Ndufv2 Cox5a Ndufa7 Ndufs4 Uqcrc1 | 6 |
| GOBP ATP SYNTHESIS COUPLED ELECTRON TRANSPORT | 1.47E-07 | 9.37E-05 | 6.90E-01 | -0.581 | -2.156 | 89 | COX2 Ndufc1 ND4L CYTB Ndufs5 COX3 ND5 COX1 ND2 ND1 ND3 Cox6b1 Ndufab1 Cox7c Pink1 Ndufb4 Cox6c Ndufa13 Uqcrb Ndufv2 Cox5a Iscu Ndufa7 Ndufs4 Uqcrc1 | 7 |
| GOMF EXTRACELLULAR MATRIX STRUCTURAL CONSTITUENT | 1.58E-07 | 9.37E-05 | 6.90E-01 | 0.51 | 2.044 | 129 | Lamc1 Tinagl1 Ccn1 Col4a1 Col4a5 Ltbp1 Fbn1 Mgp Lama1 Lamb2 Ecm1 Col16a1 Fbln1 Col25a1 Fn1 Mfap2 Nid2 Col18a1 Aebp1 Cilp Bgn Tuft1 Abi3bp Cd4 Sbspon Nid1 Col6a1 Vwf Col13a1 Pxdn Col6a3 Lama2 Matn1 Col8a2 Lama5 Srpx Col5a3 Col3a1 Igfbp7 Col5a1 Col6a2 Emilin2 Thbs1 Col1a1 Col4a3 Pcolce Emilin1 Thbs2 Fbln5 Col5a2 Dpt Col7a1 Col6a6 Prg4 Eln Fgl2 Fbn2 Col4a2 Colq Col4a6 Col9a2 Spon1 | 8 |
| GOCC INNER MITOCHONDRIAL MEMBRANE PROTEIN COMPLEX | 1.81E-07 | 9.37E-05 | 6.90E-01 | -0.534 | -2.091 | 125 | ATP8 Ndufc1 ND4L CYTB Ndufs5 ND5 COX1 ND2 ND1 ATP6 Dmac2 Atp5mg ND3 Atp5pd Ndufab1 Cox7c Atp5me Chchd6 Mcub Ndufb4 Timm10 Sdhb Immp1l Atp5f1d Ndufa13 Timm23 Uqcrb Ndufv2 Atp5f1c Cox5a Grpel1 Ndufa7 Hspa9 Ndufs4 Uqcrc1 Pam16 Atp5mc3 Ndufs8 Afg3l2 Micos10 Ndufs7 Mtx3 Apoo Sdhc Atp5f1a Atp5po Ndufs3 Cox6a2 Ndufb2 Cox6a1 | 9 |
| GOCC RESPIRASOME | 1.91E-07 | 9.37E-05 | 6.90E-01 | -0.577 | -2.143 | 89 | COX2 Ndufc1 ND4L CYTB Ndufs5 COX3 ND5 COX1 ND2 ND1 Dmac2 ND3 Cox6b1 Ndufab1 Cox7c Oxa1l Ndufb4 Sdhb Ndufa13 Uqcrb Ndufv2 Cox5a Ndufa7 Ndufs4 Uqcrc1 Ndufs8 Ndufs7 Stmp1 Sdhc Ttc19 Higd1a Ndufs3 Cox6a2 Ndufb2 Cox6a1 | 10 |
| GOMF ELECTRON TRANSFER ACTIVITY | 4.61E-07 | 2.06E-04 | 6.75E-01 | -0.53 | -2.051 | 116 | COX2 Ndufc1 ND4L CYTB Ndufs5 Aox1 COX3 ND5 COX1 ND2 ND1 Cyp19a1 Cox11 ND3 Cox6b1 Ndufab1 Cox7c Ndufb4 Ndufaf2 Sdhb Cox6c Ndufa13 Cyb5a Ndufv2 Cox5a Ndufa7 Ndufs4 Uqcrc1 | 11 |
| GOBP RESPIRATORY ELECTRON TRANSPORT CHAIN | 5.20E-07 | 2.12E-04 | 6.59E-01 | -0.548 | -2.085 | 103 | COX2 Ndufc1 ND4L CYTB Ndufs5 COX3 ND5 COX1 ND2 ND1 ND3 Cox6b1 Ndufab1 Cox7c Pink1 Ndufb4 Slc25a18 Sdhb Cox6c Ndufa13 Uqcrb Ndufv2 Cox5a Iscu Ndufa7 Ndufs4 Uqcrc1 | 12 |
| GOBP CELLULAR RESPIRATION | 5.90E-07 | 2.22E-04 | 6.59E-01 | -0.475 | -1.931 | 166 | COX2 Ndufc1 ND4L CYTB Ndufs5 COX3 ND5 COX1 ND2 ND1 ND3 Bnip3 Cox6b1 Ndufab1 Actn3 Cox7c Nr4a3 Oxa1l Slc25a23 Pink1 Dlst Ndufb4 Slc25a18 Sdhb Coq10a Cox6c Pik3ca Atp5f1d Ndufa13 Uqcrb Ndufv2 Cox5a Iscu Ndufa7 Trex1 Ndufs4 Uqcrc1 | 13 |
| GOCC ORGANELLE INNER MEMBRANE | 1.56E-06 | 5.46E-04 | 6.44E-01 | -0.354 | -1.608 | 484 | COX2 ATP8 Ndufc1 ND4L CYTB Coq5 Ndufs5 COX3 ND5 COX1 Lrpprc ND2 ND1 Mrps17 Tmem65 ATP6 Mrpl46 Cox11 Dmac2 Atp5mg Sfxn4 Tmem177 ND3 Atp5pd Mrps35 Ecsit Slc25a36 Cox6b1 Mrps7 Slc25a29 Spns1 Atpsckmt Cds2 Ndufab1 Cox7c Emd Sfxn1 Mrpl23 Mrpl41 Oxa1l Slc25a23 Atp5me Chchd6 Mcub Pink1 Slc25a48 Mrps34 Ndufb4 Slc25a18 Tomm40 Ndufaf2 Mrpl19 Mrpl32 Timm10 Sdhb Mrps18b Mrpl43 Coq10a Atp23 Cox6c Nrm Immp1l Cps1 Mtch2 Mrpl40 Myoc Atp5f1d Tdh Ndufa13 Timm23 Clpx Uqcrb Tmx4 Lemd3 Cox18 Dele1 Ndufv2 Efhd1 Atp5f1c Coq4 Slc25a21 Maip1 Cox5a Grpel1 Endog Ndufa7 Mrpl33 Hspa9 mrpl24 Ndufs4 Mtch1 Uqcrc1 Nutf2 Sfxn5 Pam16 Mrpl54 Atad3a Hadhb Tra2b Lyn Rsad2 Atp5mc3 Ndufs8 Mrps15 Dpy19l1 Chdh Mrps14 Ccdc51 Afg3l2 Micos10 Hadha Ndufs7 Stmp1 Mtx3 Apoo Lmnb2 Sdhc Mrpl57 Slc27a1 Ttc19 Higd1a Atp5f1a Coq7 Atp5po Rcc1l Ndufs3 Slc25a43 Smad1 Pde2a Tafazzin Pgs1 Cox6a2 Slc25a42 Ndufb2 Cox6a1 | 14 |
| GOBP WOUND HEALING | 1.82E-06 | 5.93E-04 | 6.44E-01 | 0.357 | 1.634 | 422 | Wnt4 Elk3 Irf1 Ccn1 Serpine1 Fgf10 Alox5 F11r Dag1 Myof C1galt1c1 Itpr2 Prkcb Anxa1 F3 Wnt5b Dsp Dgkz Gpx1 Vav2 Pros1 Fbln1 Pdgfrb Fn1 Vcl Csrp1 Itpr1 Plscr1 Erbb2 Mical1 Adora2a Tgfb3 Cflar Scube1 Rad51c Myl9 Hnf4a Gsn Ptk2 Flna Ano6 Rhoc P2rx4 Ppara Kdr Jmjd1c Plat Mylk Ptpn6 Sdc4 Anxa6 Fzd7 Itga5 Usf1 Ocln Dock6 Tec Alox12 Itgb1 Pla2g4a Serping1 Phldb2 B4galt1 Gna14 Hspb1 Smpd1 Pdgfa Anxa5 Gata2 Rhoa Nf1 Vwf Hps4 Dock9 Cd9 Itga2 Fgfr1op2 Enpp4 Foxc2 Plaur Smoc2 Ddr1 Axl Tln1 Prkca Vegfa Clec1b Trpc7 Pear1 St3gal4 Mcam Pdgfb Dcbld2 Pdgfra Drd5 Hps6 Col3a1 Col5a1 Jak2 Ephb2 Thbs1 Dgkg Col1a1 Prcp Notch2 | 15 |
| GOCC LUMENAL SIDE OF MEMBRANE | 1.98E-06 | 5.93E-04 | 6.27E-01 | 0.715 | 2.201 | 31 | Sppl2a Hspa8 RT1-M3-1 RT1-CE5 RT1-Da RT1-Ba RT1-M6-2 RT1-Bb Tapbp RT1-CE7 RT1-CE4 RT1-M6-1 Cd74 Ctsa RT1-Db1 | 16 |
| GOBP OLEFINIC COMPOUND METABOLIC PROCESS | 2.06E-06 | 5.93E-04 | 6.27E-01 | 0.558 | 2.07 | 79 | Wnt4 Alox5 Rest Star Fads1 Gpx1 Bmp6 Gstm2 Stat5b Elovl5 Ptgds H6pd Hsd17b4 Srd5a2 Cyp2j4 Ephx1 Gsta6 Alox12 Ptges2 Pla2g4a Aldh1a3 Srd5a1 Bmp5 Cyp11b2 | 17 |
| GOBP POSITIVE REGULATION OF CELL ADHESION | 2.36E-06 | 6.43E-04 | 6.27E-01 | 0.37 | 1.665 | 343 | Wnt4 Ccn1 RT1-M3-1 Hyal1 Alox5 Cd46 F11r Dag1 Jak3 Egfl6 Tjp1 Efnb1 Anxa1 Col16a1 Rela Fn1 Dock5 RT1-Da Cr1l Erbb2 Akt1 Flna Ptpru Cyth3 Flot1 S100a10 Stat5b Kdr P4hb Stx3 Rara Ptn Epb41l5 Megf10 Ptpn6 Nrp1 Sdc4 Map3k8 Itga5 Lgals1 Dmp1 Bmp7 Dnm2 Iqgap1 Cd4 Nck1 Itpkb Nid1 Rhoa Il6st Cd3e Itga2 Arpc2 Tfrc Foxc2 Ccdc80 Plaur Egflam Cd74 Hlx Prkca Vegfa Igf2 Apbb1ip Myb Chrd Rhod Igfbp2 Lilrb3 Myadm St3gal4 Fstl3 Il23a Selenok Pdgfb Ptk2b Sfrp1 Sox12 RT1-Db1 Cbfb Il1b Sash3 Has2 Jak2 Fadd Smad7 Prkd2 Itga3 Emilin1 Adam9 Rsu1 Kif26b Ap3b1 Tfe3 Socs5 Rin2 Ilk Cd276 Csf1 Zbtb7b Ecm2 Nkap Angpt1 Itgb2 Disc1 Tek Icoslg Hspd1 Sart1 | 18 |
| GOMF STRUCTURAL CONSTITUENT OF RIBOSOME | 2.98E-06 | 7.68E-04 | 6.27E-01 | -0.486 | -1.928 | 138 | Rps2 Rps23 Mrps17 Mrpl46 Mrps35 Rplp1 Mrps7 Rpl27a Rps15a Mrpl23 Mrpl41 Mrps34 Rpl27 Rps13 Rpsa Rpl22l1 Mrpl19 Mrpl32 Mrps18b Mrpl43 Rpl28 Rpl15 Rpl12 Rps7 Rpl37a-ps1 Rps8 Srbd1 Rps20 Rpl29 Ndufa7 Mrpl33 mrpl24 Rps12 Rpl10a Mrpl54 Rsl24d1 Rpl19 Rpl3 Mrps15 Mrps14 Rps5 Rpl23a Rps27 Mrpl57 Rps3a Rpl5 Rpl36al Mrpl35 Rpl18 Rps15 Rps11 Mrpl4 Mrpl16 Rpl17 | 19 |
| GOCC PRESYNAPSE | 3.21E-06 | 7.86E-04 | 6.27E-01 | -0.354 | -1.601 | 463 | Slc6a5 Adcy8 Ush2a Hap1 Adra1a Tnn Efnb2 Kcnip3 Cntn5 Atp2b4 Nrxn2 Calb1 Rab3c Syn3 Hip1 Cbarp Calca Znrf1 Slc22a3 Ntsr1 Adcyap1 Slc8a3 Pcdh8 Cad Penk Rab10 Ntf3 Cntn6 Ptprn2 Kcnk9 Nlgn3 Syngr1 Unc13c Syt10 Baiap3 Dennd1a Snap47 Scamp1 Lin7a Apba1 Th Grm2 Hcn3 Slc6a17 Cyp46a1 Slc5a7 Slc18a2 Calb2 Slc32a1 Otof Ppp3cc Stx7 Slc1a1 Grap Cdk5r1 Kcna3 Prss12 Slc2a8 Dgki Fmr1 Cdh2 Dnajc5 Gad2 C1ql1 Grik3 Stx1b Rab3b Ntng1 P2ry4 Igsf21 Btbd8 Syt5 Rab2b Gabra2 Tmem163 Gpm6a Nufip1 C1qbp Fzd3 Nlgn1 Dnm3 Sncaip Ppt1 Cntnap4 Rims4 Kirrel3 Srcin1 Slc29a4 Sncb Syt2 Slc17a5 Rab3d Zdhhc17 Dnajc6 Nptn Syn2 Kcnc1 Nlgn2 Gad1 Grik2 Ap3d1 Vdac2 Gabbr1 Stxbp1 Svop Trappc4 Cadps2 Sri Prrt2 Lin7c Ptprd Vti1b Mme Gnb5 Crhr2 Rps27 Adam10 Arhgap44 Nrxn1 Vamp2 Sypl2 Sv2c Syt4 Vamp4 Cdk5 Hcrt Ndel1 Rims3 Vdac1 Doc2a Oprm1 Pde2a | 20 |
| GOBP UROGENITAL SYSTEM DEVELOPMENT | 4.45E-06 | 1.04E-03 | 6.11E-01 | 0.391 | 1.72 | 276 | Wnt4 Sulf2 Amer1 Col4a1 Fgf10 Enpep Bmp4 Fbn1 Dync2h1 Greb1l Lamb2 Zmpste24 Bicc1 Nphp3 Traf3ip1 Frs2 Notch1 Anxa1 Hpgd Cat Pdgfrb Jmjd6 Bmp6 Glis2 Foxc1 Itga8 Cflar Acta2 Adamts1 Tfap2b Cyp7b1 Rgn Myocd Sdc4 Tbx18 Frem2 Car2 Adamts16 Mmp9 Id3 Bmp7 Ren Foxd1 Iqgap1 Crip1 Ift88 Srd5a1 Kank2 Nid1 Pdgfa Lrp4 Gata2 Nf1 Magi2 Sall1 Foxc2 Plag1 Wnk4 Ctsh Wnt6 Vegfa Wwtr1 Fstl3 Osr1 Pdgfb Sfrp1 Aldh1a2 Rbp4 Tp73 Lama5 Stra6 Pdgfra Has2 Six2 Angpt2 Fadd Ephb2 Klhl3 Sulf1 Smad7 Col4a3 Smad6 Notch2 Itga3 | 21 |
| GOBP ELECTRON TRANSPORT CHAIN | 5.38E-06 | 1.20E-03 | 6.11E-01 | -0.472 | -1.894 | 150 | COX2 Ndufc1 ND4L CYTB Ndufs5 Aox1 COX3 ND5 COX1 ND2 ND1 Cyp19a1 Cox11 ND3 Cox6b1 Ndufab1 Cox7c Pink1 Ndufb4 Slc25a18 Ndufaf2 Sdhb Cox6c Ndufa13 Cyb5a Uqcrb Ndufv2 Cox5a Iscu Ndufa7 Ndufs4 Uqcrc1 | 22 |
| GOBP GLAND DEVELOPMENT | 6.43E-06 | 1.33E-03 | 6.11E-01 | 0.365 | 1.636 | 335 | Wnt4 Cebpb Pml Sulf2 Onecut2 Fgf10 Bmp4 Cebpg Dag1 Lama1 Eda Zmpste24 Nphp3 Frs2 Notch1 Anxa1 Msn Orai1 Robo1 Ncor2 Gpx1 Rela Xdh Foxc1 Tgfb3 Cflar Hk2 Cdo1 Akt1 Hnf4a Ak4 Cyp7b1 Rgn Stat5b Taf10 Ptn Hpn Jarid2 Edar Lmo4 Lrp6 Rps6ka1 Oxtr Tgfbr3 Cdh1 Stat5a Bmp7 Wnt10a Man2a1 Crip1 Aldh1a3 Srd5a1 Pdgfa Met Gata2 Nf1 Sall1 Itga2 Kalrn Plag1 Prmt5 Ddr1 Hlx Oas2 Vegfa Igf2 Twsg1 Atp7a Tbx1 Fstl3 Igf2r Sfrp1 Aldh1a2 Plxna1 Pnpt1 Btbd7 Hmgcs1 Lama5 Sox10 Stra6 Wdr35 Pdgfra Atp2c2 Dkk3 Jak2 Fadd Sulf1 Apln Notch2 Msx1 Rbpj Slc6a3 Sox9 Ucp2 Ephb3 Ripk3 Csf1 Sec63 Zbtb7b Raf1 Scrib Sostdc1 Hmox1 Bcl11b Lrp5 | 23 |
| GOBP ANTIGEN PROCESSING AND PRESENTATION OF ENDOGENOUS PEPTIDE ANTIGEN | 6.51E-06 | 1.33E-03 | 6.11E-01 | 0.762 | 2.142 | 21 | Erap1 RT1-M3-1 RT1-CE5 B2m RT1-Da RT1-M6-2 Tap1 Tapbp RT1-CE7 RT1-CE4 RT1-M6-1 RT1-Db1 | 24 |
| GOBP GLAND MORPHOGENESIS | 8.67E-06 | 1.70E-03 | 5.93E-01 | 0.51 | 1.955 | 97 | Wnt4 Cebpb Pml Sulf2 Fgf10 Bmp4 Dag1 Lama1 Eda Frs2 Notch1 Msn Tgfb3 Cflar Cyp7b1 Ptn Hpn Edar Lrp6 Rps6ka1 Bmp7 Crip1 Pdgfa Plag1 Ddr1 Twsg1 Sfrp1 Plxna1 Btbd7 Lama5 Sulf1 Notch2 Sox9 Csf1 Scrib Sostdc1 Lrp5 Ceacam1 Mdk Wnt5a Rarg Tgm2 Tnc Cav3 Fgf1 Epha2 Cav1 Tgfbr2 Tnfaip3 Tbx2 Fgfr2 | 25 |
